# Comparative genomics of *Chlamydomonas*

**DOI:** 10.1101/2020.06.13.149070

**Authors:** Rory J. Craig, Ahmed R. Hasan, Rob W. Ness, Peter D. Keightley

## Abstract

Despite its fundamental role as a model organism in plant sciences, the green alga *Chlamydomonas reinhardtii* entirely lacks genomic resources for any closely related species, obstructing its development as a study system in several fields. We present highly contiguous and well-annotated genome assemblies for the two closest known relatives of the species, *Chlamydomonas incerta* and *Chlamydomonas schloesseri*, and a third more distantly related species, *Edaphochlamys debaryana*. We find the three *Chlamydomonas* genomes to be highly syntenous with similar gene contents, although the 129.2 Mb *C. incerta* and 130.2 Mb *C. schloesseri* assemblies are more repeat-rich than the 111.1 Mb *C. reinhardtii* genome. We identify the major centromeric repeat in *C. reinhardtii* as an L1 LINE transposable element homologous to Zepp (the centromeric repeat in *Coccomyxa subellipsoidea*) and infer that centromere locations and structure are likely conserved in *C. incerta* and *C. schloesseri*. We report extensive rearrangements, but limited gene turnover, between the minus mating-type loci of the *Chlamydomonas* species, potentially representing the early stages of mating-type haplotype reformation. We produce an 8-species whole-genome alignment of unicellular and multicellular volvocine algae and identify evolutionarily conserved elements in the *C. reinhardtii* genome. We find that short introns (<~100 bp) are extensively overlapped by conserved elements, and likely represent an important functional class of regulatory sequence in *C. reinhardtii*. In summary, these novel resources enable comparative genomics analyses to be performed for *C. reinhardtii*, significantly developing the analytical toolkit for this important model system.

## Introduction

With the rapid increase in genome sequencing over the past two decades, comparative genomics analyses have become a fundamental tool in biological research. As the first sets of genomes for closely related eukaryotic species became available, pioneering comparative studies led to refined estimates of gene content and orthology, provided novel insights into the evolution of genome architecture and the extent of genomic synteny between species, and enabled the proportions of genomes evolving under evolutionary constraint to be estimated for the first time (Mouse Genome Sequencing Consortium 2002; Cliften et al. 2003; Stein et al. 2003; Richards et al. 2005). As additional genomes were sequenced it became possible to produce multiple species whole-genome alignments (WGA) and to identify conserved elements (CEs) in noncoding regions for several of the most well-studied lineages (Siepel et al. 2005; Stark et al. 2007; Gerstein et al. 2010; Lindblad-Toh et al. 2011). Many of these conserved noncoding sequences overlap regulatory elements, and the identification of CEs has proved to be among the most accurate approaches for discovering functional genomic sequences (Alföldi and Lindblad-Toh 2013). As a result, CEs have frequently been used to enhance genome annotation projects and to study several aspects of regulatory sequence evolution (Mikkelsen et al. 2007; Lowe et al. 2011; Halligan et al. 2013; Williamson et al. 2014).

The ability to perform comparative analyses is contingent on the availability of genome assemblies for species that span a range of appropriate evolutionary distances. While this state has been achieved for the majority of model organisms, there remain several species of high biological significance that entirely lack genomic resources for any closely related species. Hiller et al. (2013) described such cases as ‘phylogenetically isolated genomes’, specifically referring to species for which the most closely related genomes belong to species divergent by one or more substitutions, on average, per neutrally evolving site. At this scale of divergence an increasingly negligible proportion of the genome can be aligned at the nucleotide-level (Margulies et al. 2006), limiting comparative analyses to the protein-level and impeding the development of such species as model systems in numerous research areas.

The unicellular green alga *Chlamydomonas reinhardtii* is a long-standing model organism across several fields, including cell biology, plant physiology and molecular biology, and algal biotechnology (Salomé and Merchant 2019). Because of its significance, the ~110 Mb haploid genome of *C. reinhardtii* was among the earliest eukaryotic genomes to be sequenced (Grossman et al. 2003; Merchant et al. 2007), and both the genome assembly and annotation are actively being developed and improved (Blaby et al. 2014). Despite its quality and extensive application, the *C. reinhardtii* genome currently meets the ‘phylogenetically isolated’ definition. The closest confirmed relatives of *C. reinhardtii* that have genome assemblies belong to the clade of multicellular algae that includes *Volvox carteri*, the *Tetrabaenaceae-Goniaceae-Volvocaceae*, or TGV clade. Collectively, *C. reinhardtii* and the TGV clade are part of the highly diverse order Volvocales, and the more taxonomically limited clades *Reinhardtinia* and core-*Reinhardtinia* (Nakada et al. 2008; Nakada et al. 2016). Although these species are regularly considered close relatives, multicellularity likely originated in the TGV clade over 200 million years ago (Herron et al. 2009), and *C. reinhardtii* and *V. carteri* are more divergent from one another than human is to chicken (Prochnik et al. 2010).

Without a comparative genomics framework, the wider application of *C. reinhardtii* as a model system is severely impeded. While this broadly applies to the general functional annotation of the genome as outlined above (e.g. refinement of gene models and annotation of CEs), it is particularly relevant to the field of molecular evolution. Although the evolutionary biology of *C. reinhardtii* has not been widely studied, the species has several features that have attracted recent attention to its application in this field. Its haploid state, high genetic diversity (~2% genome-wide (Craig et al. 2019)) and experimental tractability make it an excellent system to study the fundamental evolutionary processes of mutation (Ness et al. 2015; Ness et al. 2016), recombination (Liu et al. 2018; Hasan and Ness 2020), and selection (Böndel et al. 2019). However, without genomic resources for closely related species it is currently impossible to perform several key analyses, such as the comparison of substitution rates at synonymous and non-synonymous sites of protein-coding genes (i.e. calculating dN/dS), and the inference of ancestral states at polymorphic sites (a requirement of several population and quantitative genetics models (Keightley and Jackson 2018)).

Furthermore, *V. carteri* and its relatives in the TGV clade are extensively used to study the evolution of multicellularity and other major evolutionary transitions (e.g. isogamy to anisogamy), and five genomes of multicellular species spanning a range of organismal complexities have now been assembled (Prochnik et al. 2010; Hanschen et al. 2016; Featherston et al. 2018; Hamaji et al. 2018). These studies have often included analyses of gene family evolution, reporting expansions in families thought to be functionally related to multicellularity. While these analyses have undoubtedly made important contributions, they are nonetheless limited in their phylogenetic robustness, as *C. reinhardtii* is the only unicellular relative within hundreds of millions of years available for comparison. Thus, the availability of annotated genomes for unicellular relatives of *C. reinhardtii* will also serve as an important resource towards reconstructing the ancestral core-*Reinhardtinia* gene content, potentially providing novel insights into the major evolutionary transitions that have occurred in this lineage.

Here we present highly contiguous and well-annotated genome assemblies for the two closest known relatives of *C. reinhardtii*, namely *Chlamydomonas incerta* and *Chlamydomonas schloesseri*, and a more distantly related unicellular species, *Edaphochlamys debaryana*. Via comparison to the genomes of *C. reinhardtii* and the TGV clade species we present the first insights into the comparative genomics of *Chlamydomonas*, focussing specifically on the conservation of genome architecture between species and the landscape of sequence conservation in *C. reinhardtii*. While forming only one of the initial steps in this process, by providing the first comparative genomics framework for the species we anticipate that these novel genomic resources will greatly aid in the continued development of *C. reinhardtii* as a model organism.

## Results & Discussion

### The closest known relatives of Chlamydomonas reinhardtii

Although the genus *Chlamydomonas* consists of several hundred unicellular species it is highly polyphyletic (Pröschold et al. 2001), and *C. reinhardtii* is more closely related to the multicellular TGV clade than the majority of *Chlamydomonas* species. Given their more conspicuous morphology, the TGV clade contains ~50 described species (Herron et al. 2009), while the unicellular lineage leading to *C. reinhardtii* includes only two other confirmed species, *C. incerta* and *C. schloesseri* (Pröschold et al. 2005; Pröschold et al. 2018). As *C. reinhardtii* is the type species of *Chlamydomonas*, these three species collectively comprise the monophyletic genus (fig.1a, b, c), and *Chlamydomonas* will be used specifically to refer to this clade throughout.

*C. incerta* is the closest known relative of *C. reinhardtii*, and a small number of comparative genetics analyses have been performed between the two species (Ferris et al. 1997; Popescu et al. 2006; Smith and Lee 2008). *C. incerta* is known from only two isolates, and we selected the original isolate SAG 7.73 for sequencing. Unfortunately, although *C. incerta* SAG 7.73 is nominally from Cuba, the geographic origin of the isolate is uncertain due to a proposed historical culture replacement with *C. globosa* SAG 81.72 from the Netherlands (Harris et al. 1991). As the direction of replacement is unknown, the accepted taxonomic name of the species also remains undecided. *C. schloesseri* was recently described by Pröschold et al. (2018), and three isolates from a single site in Kenya exist in culture. We selected the isolate CCAP 11/173 for sequencing.

Beyond *Chlamydomonas* there are a substantial number of unicellular core-*Reinhardtinia* species with uncertain phylogenetic relationships (i.e. that may be part of the lineage including *Chlamydomonas*, the lineage including the TGV clade, or outgroups to both). Among these, the best studied species is *E. debaryana*, which was recently renamed from *Chlamydomonas debaryana* (Pröschold et al. 2018). Unlike the three described *Chlamydomonas* species, *E. debaryana* appears to be highly abundant in nature, with more than 20 isolates from across the Northern Hemisphere in culture, suggesting that it could be developed as a model for studying algal population structure and biogeography via the collection of further isolates. Draft genomes of the isolates NIES-2212 from Japan (Hirashima et al. 2016) and WS7 from the USA (Nelson et al. 2019) were recently assembled, while we selected the isolate CCAP 11/70 from Czechia for sequencing (fig. 1d).

**Figure 1.**
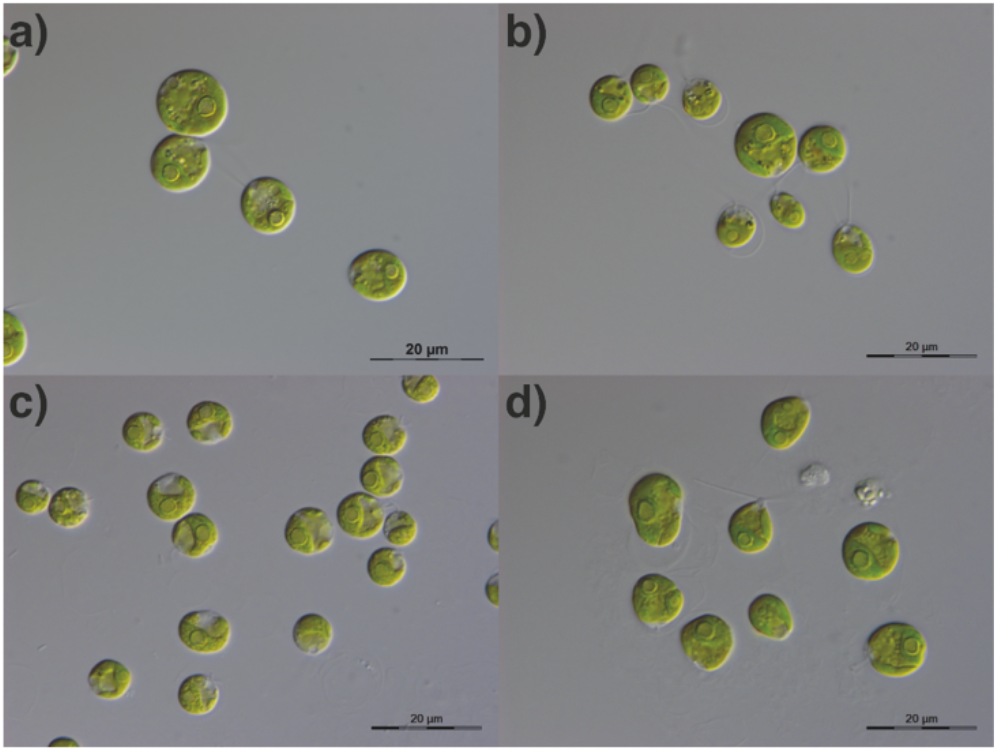
Images of unicellular species. a) *Chlamydomonas reinhardtii.* b) *Chlamydomonas incerta* SAG 7.73. c) *Chlamydomonas schloesseri* SAG 2486 (=CCAP 11/173). d) *Edaphochlamys debaryana* SAG 11.73 (=CCAP 11/70). All images taken by Thomas Pröschold.

### The genomes of Chlamydomonas incerta, Chlamydomonas schloesseri & Edaphochlamys debaryana

Using a combination of Pacific Biosciences (PacBio) long-read sequencing for *de novo* assembly (40-49x coverage, table S1) and Illumina short-read sequencing for error correction (43-86x coverage, table S2), we produced contig-level genome assemblies for *C. incerta*, *C. schloesseri* and *E. debaryana*. All three assemblies were highly contiguous, with N50s of 1.6 Mb (*C. incerta*), 1.2 Mb (*C. schloesseri*) and 0.73 Mb (*E. debaryana*), and L50s of 24, 30 and 56 contigs, respectively (table 1). Genome-mode BUSCO scores supported a high-level of assembly completeness, with the percentage of universal chlorophyte single-copy orthologs identified in each genome ranging from 95.9% to 98.1%. These metrics compare favourably to the best existing core-*Reinhardtinia* assemblies (table 1). Although the *C. reinhardtii* and *V. carteri* assemblies have greater scaffold-level N50 values than the three new assemblies, they are both considerably more fragmented at the contig level, with N50s of 215 kb and 85 kb, respectively. While this is not surprising given our application of long-read sequencing, it nonetheless demonstrates that these important model genomes could be substantially improved by additional sequencing effort. The contig-level N50s of the three new assemblies also exceeded those of the *Gonium pectorale* assembly (Hanschen et al. 2016), and the Pacbio-based assemblies of *Yamagishiella unicocca* and *Eudorina sp. 2016-703-Eu-15* (hereafter *Eudorina sp.*) (Hamaji et al. 2018), and thus they currently represent the most contiguous assemblies in terms of uninterrupted sequence in the core-*Reinhardtinia*, and indeed the entire Volvocales (table S3).

**Table 1.** Genome assembly metrics for eight high-quality core-*Reinhardtinia* genome assemblies.

| Species | *Chlamydomonas reinhardtii* v5 | *Chlamydomonas incerta* | *Chlamydomonas schloesseri* | *Edaphochlamys debaryana* | *Gonium pectorale* | *Yamagishiella unicocca* | *Eudorina. sp.* 2016-703-Eu-15 | *Volvox carteri* v2 |
| --- | --- | --- | --- | --- | --- | --- | --- | --- |
| Assembly level | chromosome | contig | contig | contig | scaffold | contig | scaffold | scaffold |
| Genome size (Mb) | 111.10 | 129.24 | 130.20 | 142.14 | 148.81 | 134.23 | 184.03 | 131.16 |
| Number of contigs/scaffolds | 17* | 453 | 457 | 527 | 2373 | 1461 | 3180 | 434 |
| N50 (Mb) | 7.78 | 1.58 | 1.21 | 0.73 | 1.27 | 0.67 | 0.56 | 2.60 |
| Contig N50 (Mb) | 0.22 | 1.58 | 1.21 | 0.73 | 0.02 | 0.67 | 0.30 | 0.09 |
| L50 | 7 | 24 | 30 | 56 | 30 | 53 | 83 | 15 |
| Contig L50 | 141 | 24 | 30 | 56 | 1871 | 53 | 155 | 410 |
| GC (%) | 64.1 | 66.0 | 64.4 | 67.1 | 64.5 | 61.0 | 61.4 | 56.1 |
| TEs & satellites (Mb / %) | 15.33 / 13.80 | 26.75 / 20.70 | 27.48 / 21.11 | 20.05 / 14.11 | 11.65 / 7.83 | 29.57 / 22.03 | 46.81 / 25.43 | 22.22 / 16.94 |
| Simple & low complexity repeats (Mb / %) | 8.71 / 7.84 | 8.57 / 7.72 | 10.19 / 9.17 | 6.40 / 5.76 | 4.15 / 3.74 | 6.55 / 4.88 | 15.15 / 8.23 | 6.45 / 5.80 |
| BUSCO genome mode (complete % / fragmented %) | 96.5 / 1.7 | 96.5 / 1.6 | 96.1 / 1.7 | 94.0 / 1.9 | 86.3 / 4.5 | 95.9 / 2.2 | 94.7 / 2.7 | 95.9 / 2.4 |

\*17 chromosomes + 37 unassembled scaffolds.
BUSCO was run using the Chlorophyta odb10 dataset. See table S3 for complete BUSCO results.

Assembled genome size varied moderately across the eight species, ranging from 111.1 Mb (*C. reinhardtii*) to 184.0 Mb (*Eudorina sp.*) (table 1). Both *C. incerta* (129.2 Mb) and *C. schloesseri* (130.2 Mb) had consistently larger genomes than *C. reinhardtii*, and *E. debaryana* (142.1 Mb) had a larger genome than both *Y. unicocca* and *V. carteri*. Although additional genome assemblies will be required to fully explore genome size evolution in the core-*Reinhardtinia*, these results suggest that *C. reinhardtii* may have undergone a recent reduction in genome size. Furthermore, while earlier comparisons between multicellular species and *C. reinhardtii* led to the observation that certain metrics of genomic complexity (e.g. gene density and intron length, see below) correlate with organismal complexity, these results indicate that genome size, at least for these species, does not. Conversely, as proposed by Hanschen et al. (2016), GC content does appear to decrease with increasing cell number, with genome-wide values ranging from 64.1 to 67.1% for the unicellular species and from 64.5 to 56.1% in the TGV clade (table 1).

The larger genome sizes of the unicellular species, relative to *C. reinhardtii,* can largely be attributed to differences in the content of transposable elements (TEs) and large satellite sequences (defined as those with monomers >10 bp), with all three species containing greater total amounts (20.1-27.5 Mb) and higher genomic proportions (14.1-21.1%) of complex repetitive sequence than *C. reinhardtii* (15.3 Mb and 13.8%) (table 1). As discussed below, the larger genome size of *E. debaryana* can also be partly attributed to the substantially higher number of genes encoded by the species. For all three assemblies, repeat content was relatively consistent across contigs, with the exception of small contigs (<~100 kb), which exhibited highly variable repeat contents and likely represent fragments of complex genomic regions that have resisted assembly (fig. S1). The higher repeat contents of the three assemblies were broadly consistent across TE subclasses (fig. S2), although a direct comparison of the TEs present in each genome is complicated by phylogenetic bias in repeat masking and classification. The existence of a curated repeat library for *C. reinhardtii* directly contributes to masking and can improve homology-based classification of repeats in related species, however this effect will become increasingly negligible as divergence increases. This is likely to at least partly explain the lower repeat content and higher proportion of “unknown” classifications observed for *E. debaryana* compared to *C. incerta* and *C. schloesseri* (table 1, fig. S2).

Nonetheless, based on manual curation of the most abundant TE families in each species, a qualitative comparison is possible. All curated TEs belonged to subclasses and superfamilies that are present in one or both of *C. reinhardtii* and *V. carteri* (the two species with existing repeat libraries), suggesting a largely common repertoire of TEs across the core-*Reinhardtinia*. Alongside the more widely recognised L1 LINE and Gypsy LTR elements, all species contained families of the comparatively obscure Dualen LINE elements (Kojima and Fujiwara 2005), PAT-like DIRS elements (Poulter and Butler 2015) and Helitron2 rolling-circle elements (Bao and Jurka 2013). We also identified families of Zisupton and Kyakuja DNA transposons, both of which were reported as potentially present in *C. reinhardtii* upon their recent discovery (Böhne et al. 2012; Iyer et al. 2014). These superfamilies are greatly understudied, and there are currently no Kyakuja elements deposited in either the Repbase (https://www.girinst.org/repbase/) or Dfam (https://www.dfam.org) repositories. Although not the main focus of this study, the annotation of elements from such understudied superfamilies highlights the importance of performing manual TE curation in phylogenetically diverse lineages. Alongside improving our understanding of TE biology, these elements are expected to contribute towards more effective repeat masking/classification and gene model annotation in related species, which will be of increasing importance given the large number of chlorophyte genome projects currently in progress (Blaby-Haas and Merchant 2019).

### Phylogenomics of the core-Reinhardtinia and Volvocales

Due to the low number of available genomes and gene annotations, the phylogenetics of the Volvocales has almost exclusively been studied using ribosomal and plastid marker genes. These analyses have successfully delineated several broad clades (e.g. *Reinhardtinia*, *Moewusinia*, *Dunaliellinia*) (Nakada et al. 2008), but often yielded inconsistent topologies for more closely related taxa. Utilising both our own and several recently published genomic resources, we further explored the phylogenomic structure of the *Reinhardtinia* and Volvocales. As several genomes currently lack gene annotations, we first used an annotation-free approach based on the identification of chlorophyte single-copy orthologous genes with BUSCO (Waterhouse et al. 2018). This dataset consisted of 1,624 genes, present in at least 15 of the 18 included species (12 *Reinhardtinia*, three other Volvocales, and three outgroups from the order Sphaeropleales, table S3). For the 11 species with gene annotations (table S4), we produced a second dataset based on the orthology clustering of each species’ proteome, which yielded 1,681 single-copy orthologs shared by all species. For both datasets, we performed maximum-likelihood (ML) analyses using IQ-TREE (Nguyen et al. 2015). Analyses were performed on both concatenated protein alignments (producing a species-tree) and individual alignments of each ortholog (producing gene trees), which were then summarised as a species-tree using ASTRAL-III (Zhang et al. 2018).

All four of the resulting phylogenies exhibited entirely congruent topologies, with near maximal-support values at all nodes (fig. 2, S3). Rooting the tree on the Sphaeropleales species, the monophyly of the Volvocales, *Reinhardtinia*, and core-*Reinhardtinia* clades were recovered. *Chlamydomonas* was recovered with the expected branching order (Pröschold et al. 2018), as was the monophyly and expected topology of the TGV clade (Nakada et al. 2019). The most contentious phylogenetic relationships are those of the remaining unicellular core-*Reinhardtinia*, which include *E. debaryana* and also the recently published genomes of *Chlamydomonas sphaeroides* (Hirashima et al. 2016) and *Chlamydomonas sp. 3112* (Nelson et al. 2019). In the most gene-rich analysis to date, *E. debaryana* was grouped in a weakly-supported clade with *Chlamydomonas* (termed metaclade C), while *C. sphaeroides* grouped with a small number of other unicellular species on the lineage including the TGV clade (Nakada et al. 2019). In our analysis, *E. debaryana* and *C. sphaeroides* were recovered as sister taxa on the lineage including *Chlamydomonas*, meeting the prior definition of metaclade C as the sister clade of the TGV clade and its unicellular relatives. Due to its recent discovery, *C. sp. 3112* has not been included in previous phylogenetic analyses. This species is a member of the core-*Reinhardtinia* based on sequence similarity of ribosomal and plastid genes, and is likely a close relative of the described species *Chlamydomonas zebra* (table S5). Given its basal phylogenetic position relative to metaclade C and the TGV clade, species such as *C. sp. 3112* could prove particularly useful in future efforts to reconstruct the ancestral gene content of the core-*Reinhardtinia*.

**Figure 2.**
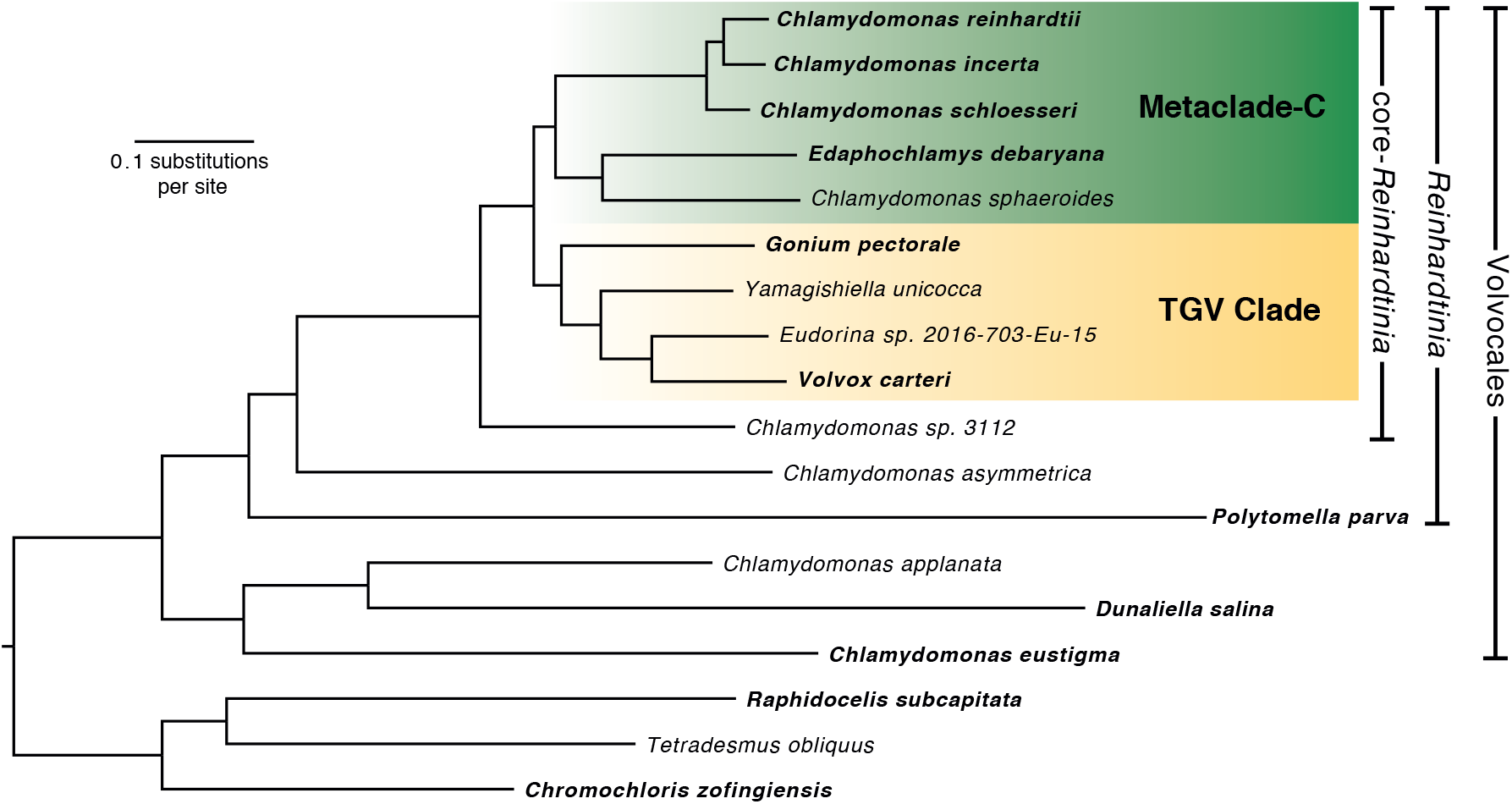
ML phylogeny of 15 Volvocales species and three outgroups inferred using LG+F+R6 model and concatenated protein alignment of 1,624 chlorophyte BUSCO genes. All bootstrap values >=99%. Species in bold have gene model annotations and were included in the OrthoFinder-based phylogenies (fig. S3b, c).

### Synteny and conserved genome architecture in Chlamydomonas

Almost nothing is known about karyotype evolution and the rate of chromosomal rearrangements in *Chlamydomonas* and the core-*Reinhardtinia*. Prochnik et al. (2010) reported that the syntenic genomic segments identified between *C. reinhardtii* and *V. carteri* contained fewer genes than human and chicken syntenic segments, in part due to a greater number of small inversions disrupting synteny. As the longest contigs in our assemblies were equivalent in length to *C. reinhardtii* chromosome arms (6.4, 4.5 and 4.2 Mb for *C. incerta*, *C. schloesseri* and *E. debaryana*, respectively), and given the closer evolutionary relationships of the unicellular species, we explored patterns of synteny between the three species and *C. reinhardtii*. We used SynChro (Drillon et al. 2014) to identify syntenic segments, which first uses protein sequence reciprocal best-hits to anchor syntenic segments, before extending segments via the inclusion of homologs that are syntenic but not reciprocal best-hits. All three *Chlamydomonas* genomes were highly syntenous, with 99.5 Mb (89.5%) of the *C. reinhardtii* genome linked to 315 syntenic segments spanning 108.1 Mb (83.6%) of the *C. incerta* genome, and 98.5 Mb (88.6%) of the *C. reinhardtii* genome linked to 409 syntenic segments spanning 108.1 Mb (83.1%) of the *C. schloesseri* genome.

Given the high degree of synteny, it was possible to order and orientate the contigs of *C. incerta* and *C. schloesseri* relative to the assembled chromosomes of *C. reinhardtii* (fig. 3). A substantial proportion of the *C. reinhardtii* karyotype appeared to be conserved in *C. incerta*, with six of the 17 chromosomes (1, 3, 4, 7, 14 and 16) showing no evidence of inter-chromosomal rearrangements, and a further three (5, 13 and 15) showing evidence for only minor translocations <150 kb in length (fig. 3a). Consistent with its greater divergence from *C. reinhardtii*, *C. schloesseri* exhibited such one-to-one conservation between only four chromosomes (5, 7, 11 and 14) (fig. 3b). For both species, patterns of synteny indicated at least one inter-chromosomal rearrangement affecting the remaining chromosomes, although without additional scaffolding of contigs it is difficult to comment on the overall effect of such rearrangements on karyotype. Furthermore, by direct comparison to *C. reinhardtii* chromosomes, we may have overestimated karyotype conservation due to undetected chromosome fusion/fission events (i.e. if a *C. reinhardtii* chromosome is present as two chromosomes in one of the related species). Across both *C. incerta* and *C. schloesseri*, all chromosomes (except chromosome 15 in the *C. incerta* comparison) contained intra-chromosomal rearrangements relative to *C. reinhardtii*, with small inversions <100 kb in length comprising the vast majority (fig. S4a, b). Synteny was far weaker between *C. reinhardtii* and *E. debaryana*, with 58.6 Mb (52.8%) of the *C. reinhardtii* genome linked to 1,975 syntenic segments spanning 64.8 Mb (45.6%) of the *E. debaryana* genome (fig. S4c). Taken together with the previous assessment of synteny between *C. reinhardtii* and *V. carteri*, these results suggest that karyotype evolution in the core-*Reinhardtinia* is expected to be dynamic, with very high levels of synteny but a non-negligible rate of inter-chromosomal rearrangements present between closely related species, and likely far greater karyotypic diversity present between more distantly related species.

**Figure 3.**
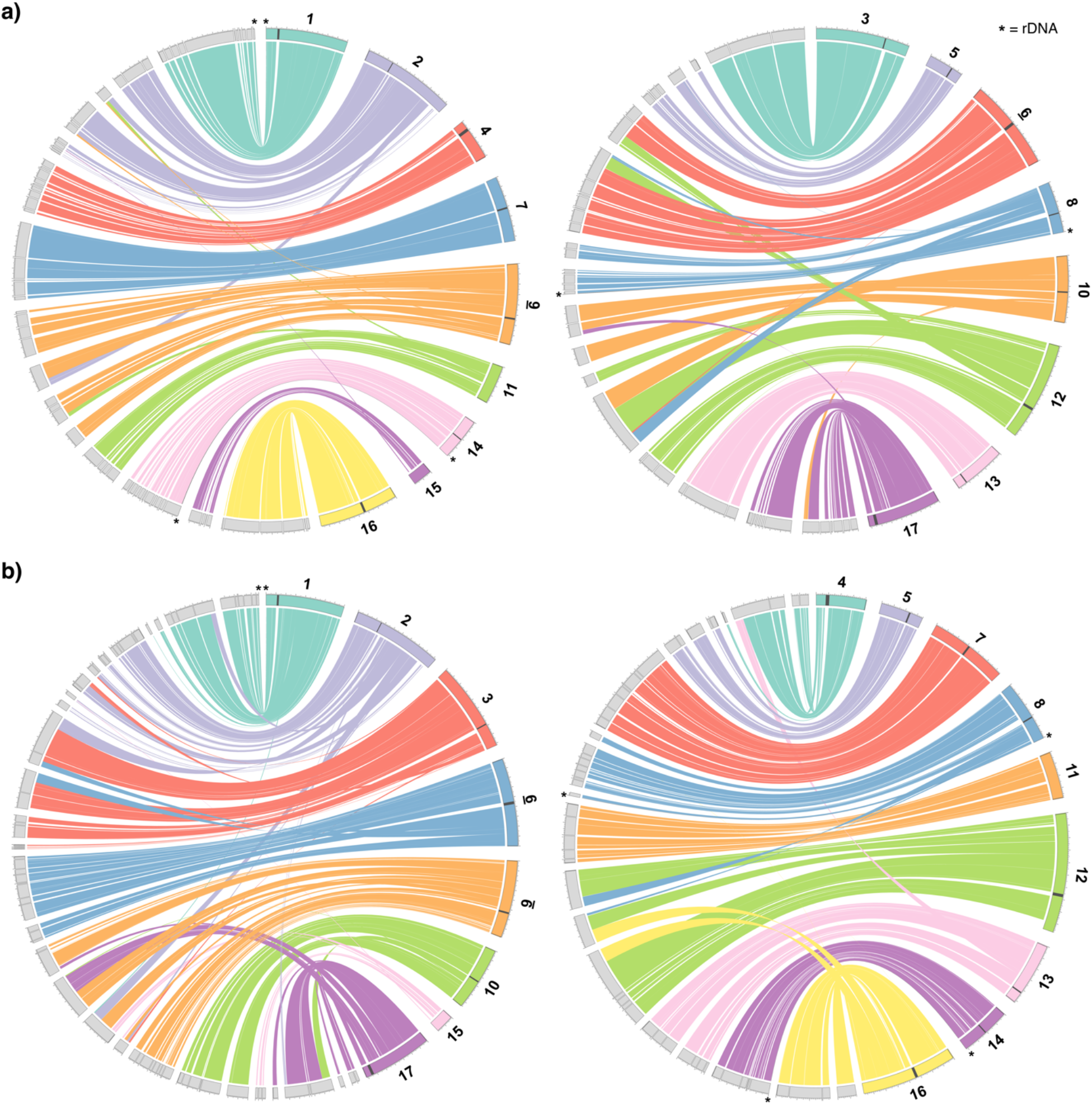
Circos plot (Krzywinski *et al*. 2009) representation of synteny blocks shared between a) *C. reinhardtii* and *C. incerta*, and b) *C. reinhardtii* and *C. schloesseri*. *C. reinhardtii* chromosomes are represented as coloured bands, and *C. incerta* / *C. schloesseri* contigs as grey bands. Contigs are arranged and orientated relative to *C. reinhardtii* chromosomes, and adjacent contigs with no signature of rearrangement relative to *C. reinhardtii* are plotted without gaps. Dark grey bands highlight putative *C. reinhardtii* centromeres, and asterisks represent rDNA. Note that colours representing specific chromosomes differ between a) and b).

Given the high-contiguity and synteny of the assemblies, it was possible to assess several complex features of genome architecture that regularly resist assembly in short-read assemblies. Telomeric repeats were observed in all three assemblies, with six *C. incerta* and 19 *C. schloesseri* contigs terminating in the satellite (TTTTAGGG)n, and 15 *E. debaryana* contigs terminating in (TTTAGGG)n (table S6). The *Arabidopsis-*type sequence (TTTAGGG)n is ancestral to green algae and was previously confirmed as the telomeric repeat present in *E. debaryana*, while the derived *Chlamydomonas-*type sequence (TTTTAGGG)n is found in both *C. reinhardtii* and *V. carteri* (Fulnečková et al. 2012). Given the phylogenetic relationships presented above (fig. 2), this implies either two independent transitions to the derived sequence or a reversion to the ancestral sequence in the lineage including *E. debaryana*, providing further evidence for the relatively frequent transitions that have produced extensive variation in telomere composition in green algae and land plants (Peska and Garcia 2020).

Ribosomal DNA repeats (rDNA) were assembled as part of three larger contigs in both *C. incerta* and *C. schloesseri*, but were found only as fragmented contigs entirely consisting of rDNA in *E. debaryana*. Although poorly assembled in *C. reinhardtii*, the rDNA arrays are located at subtelomeric locations on chromosomes 1, 8 and 14, where cumulatively they are estimated to be present in 250-400 tandem copies (Howell 1972; Marco and Rochaix 1980). The assembled *C. incerta* and *C. schloesseri* rDNA arrays (which are also not complete and are tandemly repeated at most five times) were entirely syntenous with those of *C. reinhardtii*, suggesting conservation of subtelomeric rDNA organisation in *Chlamydomonas* (fig. 3). rDNA arrays are commonly located in subtelomeric regions across several taxa, where among several other factors their location may be important for genomic stability (Dvořáčková et al. 2015).

Finally, we were able to assess the composition and potential synteny of centromeres in *Chlamydomonas*. The centromeric locations of 15 of the 17 *C. reinhardtii* chromosomes were recently mapped by Lin et al. (2018), who observed that these regions were characterised by multiple copies of genes encoding reverse-transcriptase. Upon inspection of these regions, we found that these genes are generally encoded by copies of the L1 LINE element L1-1_CR. Although these regions are currently not well-enough assembled to conclusively define the structure of centromeric repeats, L1-1_CR is present in multiple copies at all 15 putative centromeres and appears to be the major centromeric component (with chromosome-specific contributions from other TEs, especially Dualen LINE elements) (table S7, fig S5a). Remarkably, phylogenetic analysis of all curated L1 elements from green algae indicated that L1-1_CR is more closely related to the Zepp elements of *Coccomyxa subellipsoidea* than to any other L1 elements annotated in *C. reinhardtii* (fig. 4a). The divergence of the classes Trebouxiophyceae (to which *C. subellipsoidea* belongs) and Chlorophyceae (to which *C. reinhardtii* belongs) occurred in the early Neoproterozoic era (i.e. 700-1,000 million years ago) (Del Cortona et al. 2020), suggesting that L1-1_CR has been evolving independently from all other *C. reinhardtii* L1 elements for more than half a billion years. Zepp elements are of particular interest as they are thought to constitute the centromeres in *C. subellipsoidea*, where they are strictly present as one cluster per chromosome (Blanc et al. 2012). The clustering pattern of Zepp elements arises due to a nested insertion mechanism that targets existing copies, creating tandem arrays consisting mostly of the 3’ end of the elements (due to frequent 5’ truncations upon insertion) (Higashiyama et al. 1997). Chromosome-specific clustering of L1-1_CR was also evident in *C. reinhardtii*, with highly localised clusters observed at all 15 of the putative centromeres (fig. 4b). The double-peaks in L1-1_CR density present on chromosomes 2, 3 and 8, and the single sub-telomeric cluster present on chromosome 5, are all the result of the incorrect scaffolding of contigs in these highly repetitive regions (unpublished data). Thus, outside the putative centromeres, L1-1_CR appears to be entirely absent from the *C. reinhardtii* genome.

**Figure 4.**
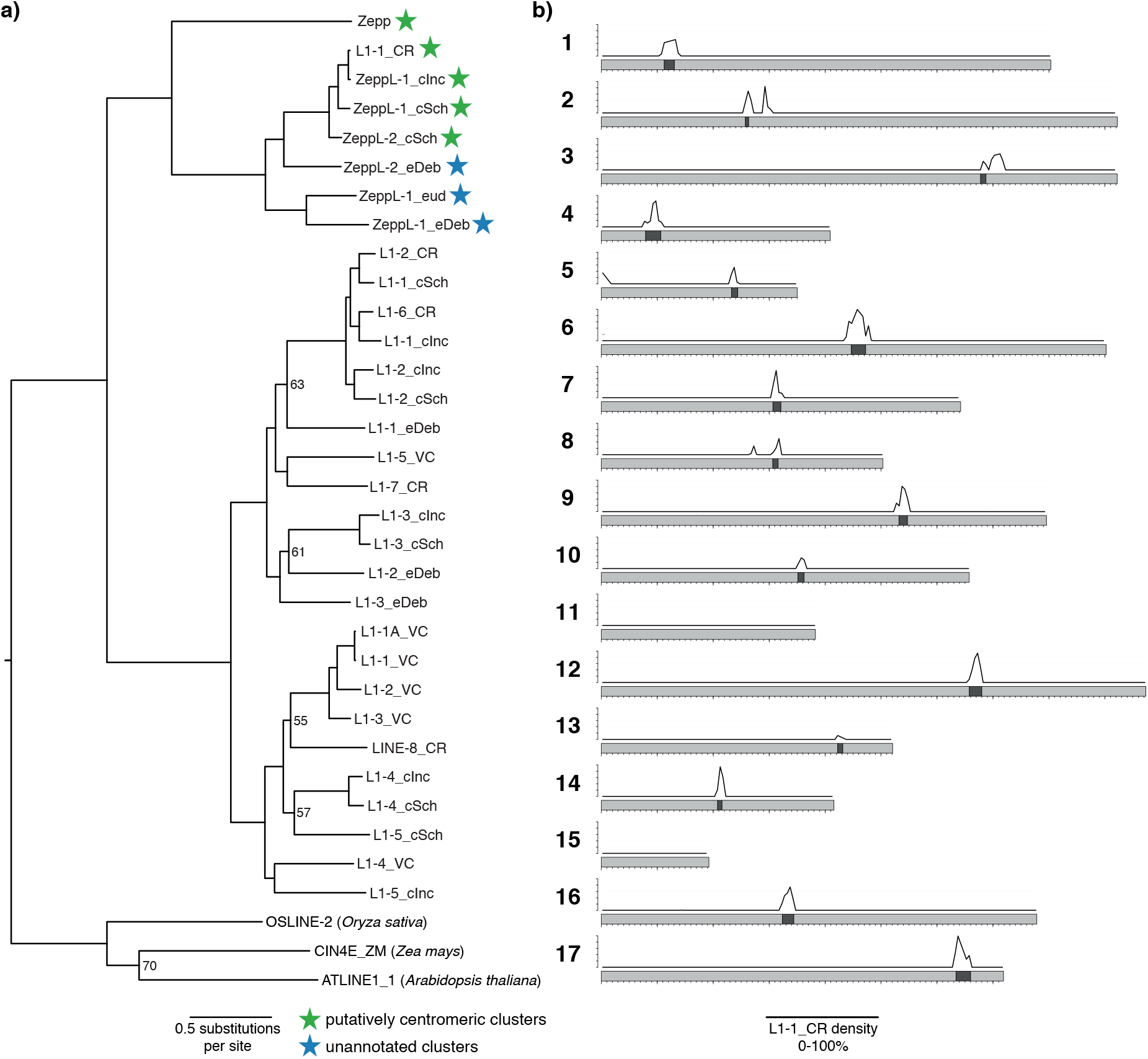
a) ML phylogeny of chlorophyte L1 LINE elements inferred using the LG+F+R6 model and alignment of endonuclease and reverse transcriptase protein domains. Bootsrap values <=70% are shown. Species are given by the element name suffix as follows: _CR = *C. reinhardtii*, _VC = *V. carteri*, _cInc = *C. incerta*, _cSch = *C. schloesseri*, _eDeb = *E. debaryana*, _eud = *Eudorina* sp. *2016-703-Eu-15*. b) Density (0-100%) of L1-1_CR in 50 kb windows across *C. reinhardtii* chromosomes. Dark bands represent putative centromeres, x-axis ticks represent 100 kb increments and y-axis ticks represent 20% increments. Plot produced using karyoploteR (Gel and Serra 2017).

Every centromeric location in *C. reinhardtii* coincided with breaks in syntenic segments and the termination of contigs in *C. incerta* and *C. schloesseri* (fig. 3), suggesting that these regions are also likely to be repetitive in both species. The phylogenetic analysis revealed the presence of one and two L1-1_CR homologs in *C. incerta* and *C. schloesseri,* respectively, which we term Zepp-like (ZeppL) elements (fig. 4a). Of the 30 contig ends associated with the 15 *C. reinhardtii* centromeres, 28 contigs in both species contained a ZeppL element within their final 20 kb (fig. S5b, c), and genome-wide the ZeppL elements exhibited similarly localised clustering to that observed in *C. reinhardtii* (fig. S6a, b). Thus, it appears that both the location and composition of the *C. reinhardtii* centromeres are likely to be conserved in both *C. incerta* and *C. schloesseri*. We further identified two families of ZeppL elements in the *E. debaryana* genome and one family of ZeppL elements in the *Eudorina sp.* genome, although we did not find any evidence for ZeppL elements in either *Y. unicocca* or *V. carteri*. Given the lack of synteny between *C. reinhardtii* and *E. debaryana* it was not possible to assign putatively centromeric contigs. Nonetheless, highly localised genomic clustering of ZeppL elements was observed for both *E. debaryana* and *Eudorina sp.* (fig. S6c, d), suggesting that these elements may play a similar role to that in *Chlamydomonas*.

As sequencing technologies advance it is becoming increasingly clear that TEs, alongside satellite DNA, contribute substantially to centromeric sequence in many species (Chang et al. 2019; Fang et al. 2020). Given the evolutionary distance between *C. subellipsoidea* and *Chlamydomonas,* it is tempting to predict that ZeppL elements may be present at the centromeres of many other species of green algae. However, it is unlikely that centromeres are conserved between species from the Trebouxiophyceae and Chlorophyceae. Firstly, it has recently been shown that the centromeric repeats in the Chlorophyceae species *Chromochloris zofingiensis* consist of LTR elements from the Copia superfamily (Roth et al. 2017). Secondly, the apparent absence of ZeppL elements from *Y. unicocca* and *V. carteri* suggest that these elements are not required for centromere formation in these species. Instead, it is possible that the propensity for Zepp and ZeppL elements to form clusters may play a role in their recruitment as centromeric sequences, which is likely to have happened independently in *C. subellipsoidea* and *Chlamydomonas*. As more highly contiguous chlorophyte assemblies become available, it will be important to search these genomes for ZeppL clusters to assess whether these elements can be used more generally as centromeric markers.

### Gene and gene family evolution in the core-Reinhardtinia

Gene model annotation was performed for each species using 7.4-8.2 Gb of stranded RNA-seq (table S8). Protein mode BUSCO scores supported a high level of annotation completeness across all three species (97.0-98.1% chlorophyte genes present), although relative to genome mode scores there was an increase in the proportion of genes identified as fragmented (4.0-5.9%) (table 2). *C. incerta* and *C. schloesseri* had comparable gene counts to *C. reinhardtii*, although lower gene densities as a result of their larger genomes. With 19,228 genes, the *E. debaryana* genome contained substantially more genes than any other currently annotated core-*Reinhardtinia* species. As reported by Hanschen et al. (2016), several metrics appeared to correlate with organismal complexity. Relative to the unicellular species, gene density was lower, and median intergenic and intronic lengths were longer, in *G. pectorale* and *V. carteri*. Presumably this is at least partly due to an increase in the amount of regulatory sequence in these genomes, although this has not yet been explored.

**Table 2.** Gene annotation metrics for core-*Reinhardtinia* species.

| Species | *Chlamydomonas reinhardtii* v5.6* | *Chlamydomonas incerta* | *Chlamydomonas schloesseri* | *Edaphochlamys debaryana* | *Gonium pectorale* | *Volvox carteri* v2.1 |
| --- | --- | --- | --- | --- | --- | --- |
| **Number of genes** | 16,656 | 16,350 | 15,571 | 19,228 | 16,290 | 14,247 |
| **Number of transcripts** | 18,311 | 16,957 | 16,268 | 20,450 | 16,290 | 16,075 |
| **Gene coverage (Mb / %)** | 91.22 / 82.10 | 94.42 / 73.06 | 94.29 / 73.42 | 103.13 / 72.55 | 65.04 / 43.71 | 84.00 / 64.04 |
| **UTR coverage (Mb / %)** | 3.88 / 15.59 | 4.04 / 11.22 | 3.37 / 9.23 | 4.24 / 9.31 | 0 / 0 | 3.00 / 11.55 |
| **Mean intron number** | 7.81 | 8.58 | 7.67 | 9.31 | 6.15 | 6.73 |
| **Median intron length (bp)** | 229 | 225 | 244 | 198 | 310 | 343 |
| **Median intergenic distance (bp)** | 134 | 341 | 408 | 555 | 2372 | 905 |
| **BUSCO protein mode (complete % / fragmented %)** | 96.1 / 2.3 | 91.1 / 5.9 | 94.7 / 3.0 | 94.1 / 4.0 | 81.5 / 12.9 | 94.7 / 2.0 |

\* *C. reinhardtii* annotation is based on a customised repeat-filtered version of the v5.6 annotation (see Methods).
Intron metrics are based only on introns within coding sequence, to avoid differences caused by the quality of UTR annotation. BUSCO was run using the Chlorophyta odb10 dataset. See table S4 for complete BUSCO results.

Across all species, both mean intron lengths (discussed below) and intron numbers per gene were very high for such compact genomes. For the unicellular species, the mean number of introns per gene coding sequence ranged from 7.7-9.3, with slightly lower counts in *G. pectorale* (6.2) and *V. carteri* (6.7). These numbers are more comparable to vertebrates such as human (8.5) than to other model organisms with similar genomes sizes, such as *Caenorhabditis elegans* (5.1), *Drosophila melanogaster* (3.0), and *Arabidopsis thaliana* (4.1). Modelling of intron evolution across the breadth of eukaryota has predicted that a major expansion of introns occurred early in chlorophyte evolution (prior to the divergence of Chlorophyceae and Trebouxiophyceae), and that high intron densities have since been maintained in certain lineages by a balance between intron loss and gain (Csuros et al. 2011). It has been hypothesised that the relative roles of DNA double-strand break repair pathways play a major role in the dynamics of intron gain and loss, as the homologous recombination (HR) pathway is thought to cause intron deletion, while non-homologous end-joining (NHEJ) may result in both intron gain and loss (Farlow et al. 2011). In is interesting to note that HR occurs at an extremely low rate in *C. reinhardtii* (Plecenikova et al. 2014), and if this is shared across *Chlamydomonas* and the core-*Reinhardtinia*, it may contribute to the maintenance of such high intron numbers. Alternatively, introns could be maintained by other forces, such as selection. Intron gains and losses caused by NHEJ are expected to possess specific genomic signatures (Farlow et al. 2011; Sun et al. 2015), and thus it should now be possible to test this hypothesis by exploring patterns of intron gain and loss across the three *Chlamydomonas* species.

To explore gene family evolution in the core-*Reinhardtinia,* we performed orthology clustering using the six available high-quality gene annotations (98,342 total protein-coding genes), which resulted in the delineation of 13,728 orthogroups containing 86,446 genes (fig. 5). The majority of orthogroups (8,532) were shared by all species, with the second most abundant category (excluding genes unique to a single species) being those present in all species except *G. pectorale* (868 orthogroups). Given the lower BUSCO score observed for *G. pectorale* (table 2) it is likely that a proportion of these orthogroups are also universal to core-*Reinhardtinia* species. The next most abundant category was the 859 orthogroups present only in *Chlamydomonas*. Unfortunately, essentially nothing is known about the biology and ecology of *C. incerta* and *C. schloesseri*, and even for *C. reinhardtii* we have a minimal understanding of its biology in natural environments (Sasso et al. 2018; Craig et al. 2019). Although many of the *Chlamydomonas*-specific orthogroups are associated with functional domains, this scenario currently precludes the formation of any clear hypotheses to test. In contrast to *Chlamydomonas*, only 51 orthogroups were unique to the two multicellular species. This may be an underestimate due to the relative incompleteness of the *G. pectorale* annotation, and it will be important to re-visit this analysis as more annotations become available (e.g. for *Y. unicocca* and *Eudorina sp.*). Nonetheless, the availability of our three new high-quality annotations for unicellular species will provide a strong comparative framework to explore the relative roles of gene family birth versus expansions in existing gene families in the transition to multicellularity.

**Figure 5.**
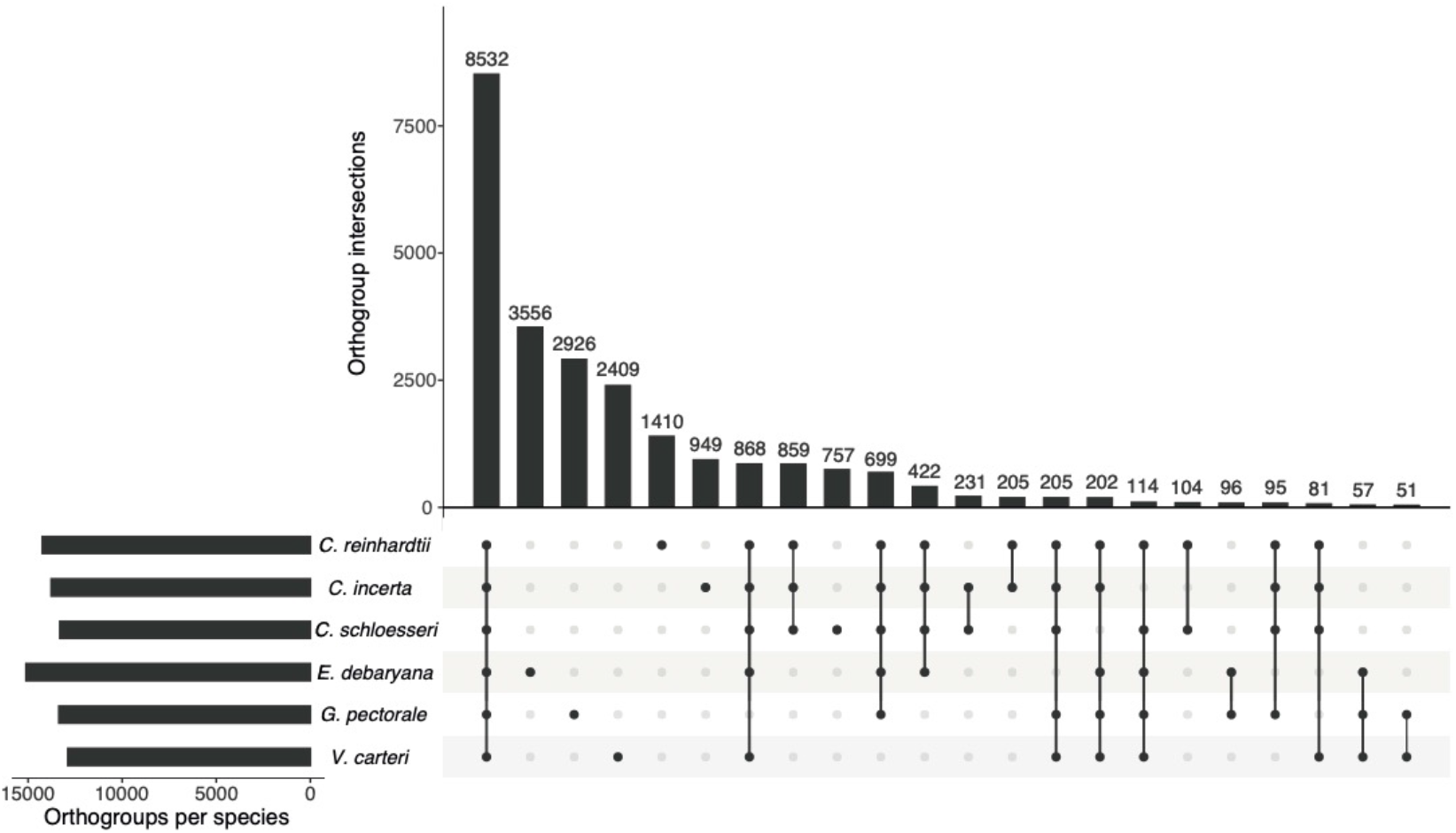
Upset plot (Lex et al. 2014) representing the intersection of orthogroups between six core-*Reinhardtinia* species. Numbers above bars represent the number of orthogroups shared by a given intersection of species.

Finally, we explored the contribution of gene family expansions to the high gene number observed in *E. debaryana*. The *E. debaryana* genome contained more species-specific genes (3,556) than any other species, however this figure was not substantially higher than the unassigned gene counts for *G. pectorale* and *V. carteri* (fig. 5). We quantified *E. debaryana* gene family expansion and contraction by calculating per orthogroup log2-transformed ratios of the *E. debaryana* gene count and the mean gene count for the other species. Arbitrarily defining an expansion as a log2-transformed ratio >1 (i.e. a given orthogroup containing more than twice as many *E. debaryana* genes than the mean of the other species) and a contraction as a ratio <−1, we identified *E. debaryana*-specific expansions in 294 orthogroups and contractions in 112. Although 192 of the expanded orthogroups were associated with functional domains (table S9), it is once again very difficult to interpret these results without any knowledge of the biological differences between unicellular species. Given that *E. debaryana* has been found across the Northern Hemisphere it is possible that the species experiences a greater range of environments than the *Chlamydomonas* species, however this is currently entirely speculative.

### Evolution of the mating-type locus in Chlamydomonas

Across core-*Reinhardtinia* species, sex is determined by a haploid mating-type locus (MT) with two alleles, termed MT+ or female, and MT- or male, in isogamous and anisogamous species, respectively. The *C. reinhardtii* MT is located on chromosome 6, spanning >400 kb and consisting of three domains, the T (telomere-proximal), R (rearranged) and C (centromere-proximal) domains. While both the T and C domains exhibit high synteny between the MT alleles, the R domain contains the only MT limited genes (Ferris and Goodenough 1997) and harbours substantial structural variation, featuring several inversions and rearrangements (Ferris et al. 2002; De Hoff et al. 2013). Crossover events are suppressed across the entire MT, although genetic differentiation between gametologs is reduced as a result of widespread gene conversion (De Hoff et al. 2013; Hasan et al. 2019). Comparative analyses of MT+/female and MT-/male haplotypes between core-*Reinhardtinia* species, and particularly between TGV clade species, have revealed highly dynamic MT evolution, with extensive gene turnover and structural variation resulting in a complex and discontinuous evolutionary history of haplotype reformation (Ferris et al. 2010; Hamaji et al. 2016b; Hamaji et al. 2018). This is most strikingly illustrated by the MT male R domains of *V. carteri* and *Eudorina sp.*, the former being ~1.1 Mb in length and relatively repeat-rich, while the latter is just 7 kb and contains only three genes (Hamaji et al. 2018). Only one MT limited gene is common to all species, the minus dominance gene *MID*, which determines MT-/male gametic differentiation (Ferris and Goodenough 1997; Yamamoto et al. 2017).

To explore whether MT evolution is similarly dynamic between the more closely related *Chlamydomonas* species, we used a reciprocal best-hit approach to identify *C. reinhardtii* MT orthologs in *C. incerta* and *C. schloesseri*. The sequenced isolates of both species were determined to be MT-based on the presence of *MID*, as was previously reported for *C. incerta* (Ferris et al. 1997). Although we were able to map the entire *C. reinhardtii* MT-haplotype to single contigs in both the *C. incerta* and *C. schloesseri* assemblies, it is important to state that it is currently impossible to define the R domain boundaries for either species without sequencing their MT+ alleles. Unfortunately, it is currently unknown if any of the one (*C. incerta*) or two (*C. schloesseri*) other isolates are MT+, and as no isolate from either species has been successfully crossed it is not even known if they are sexually viable (Pröschold et al. 2005). We also determined the sequenced isolate of *E. debaryana* to be MT-via the identification of *MID*, although we did not explore MT evolution further given the evolutionary distance to *C. reinhardtii*. At least one heterothallic mating pair of *E. debaryana* are in culture, and a future comprehensive study of MT in the species is therefore possible.

In *C. incerta,* gene order was entirely syntenic across the C domain, with the exception of *MT0828*, which did not yield a hit anywhere in the genome. Conversely, both T and R domain genes have undergone several rearrangements and inversions relative to *C. reinhardtii* MT-(fig. 6a). Furthermore, the T domain genes *SPP3* and *HDH1* were present on separate contigs in *C. incerta* and do not appear to be MT-linked (table S10). Synteny otherwise continued well into the adjacent autosomal sequence, in line with the genome-wide patterns of synteny described above. We observed even less synteny between *C. reinhardtii* and *C. schloesseri* MT-genes, with both the T and C domains showing two large inversions each (fig. 6b). However, gene order in the surrounding autosomal sequence was also largely collinear. As in *C. incerta*, *SPP3* was located elsewhere in the *C. schloesseri* assembly, suggesting a relatively recent translocation to the T domain in *C. reinhardtii*. Finally, the C domain gene *97782* was also located on a different contig, while the genes *MT0796, MT0828* and *182389* did not yield hits anywhere in the *C. schloesseri* genome. While the *C. reinhardtii* MT-limited gene *MTD1* was found in both *C. incerta* and *C. schloesseri*, we found no hits for the MT+ limited genes *FUS1* and *MTA1*, suggesting that these genes (assuming they exist) are also expected to be MT+ limited in both species.

**Figure 6.**
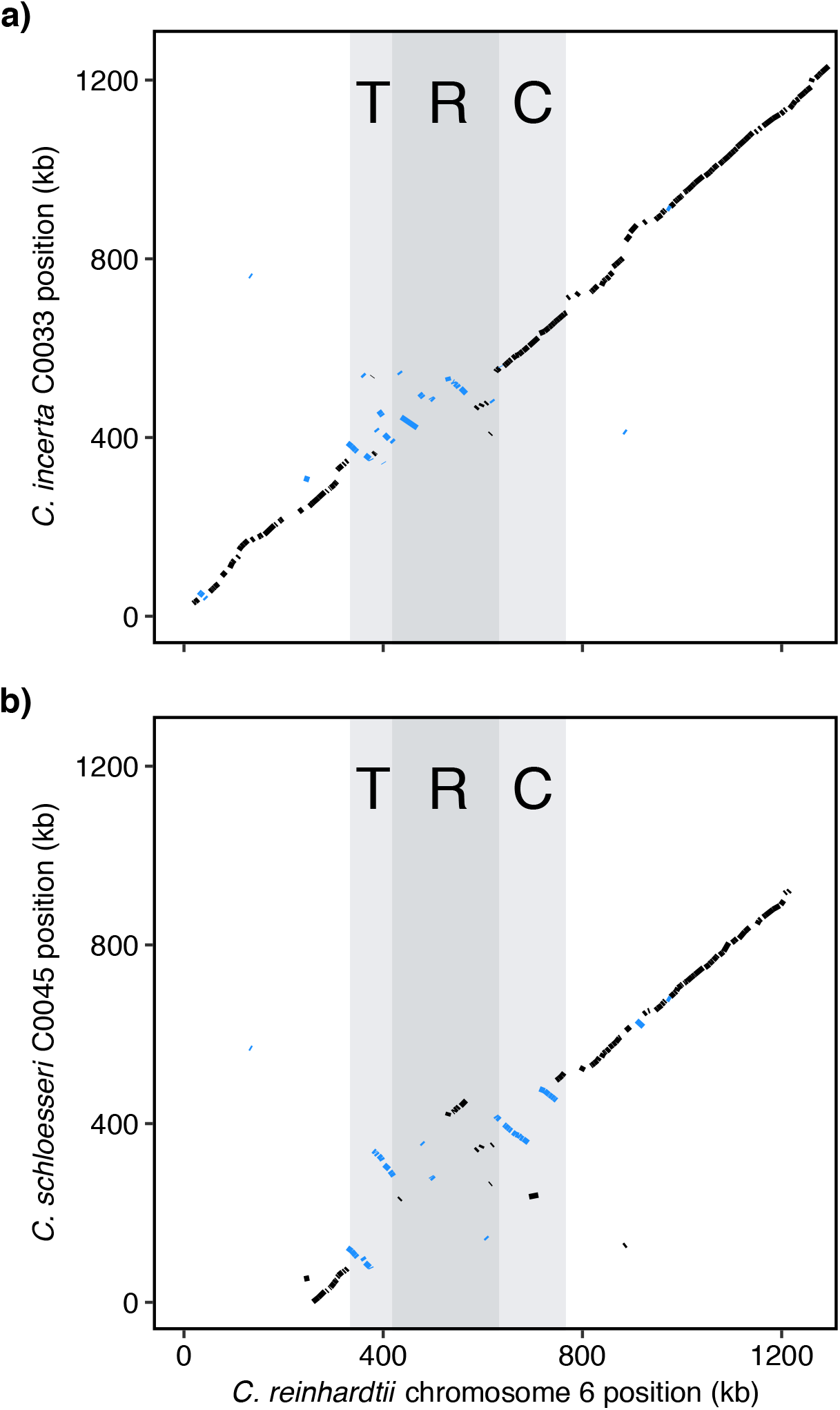
Synteny representation between genes of the *C. reinhardtii* MT-haplotype and flanking autosomal sequence, and a) inferred *C. incerta* MT-haplotype and flanking sequence genes (contig C0033), or b) inferred *C. schloesseri* MT-haplotype and flanking sequence genes (contig C0045). The T, R and C domains of the *C. reinhardtii* MT-are highlighted. Note that C0045 does not contain the initial ~260 kb of chromosome 6.

The lack of collinearity relative to the *C. reinhardtii* T domain may be indicative of an extended R domain in these species, especially in *C. schloesseri*, where we observe multiple rearrangements in all three domains. We do not, however, observe dramatic variation in MT size; whereas *C. reinhardtii* MT-is ~422 kb, if *NIC7* and *MAT3* are taken as the boundaries of the locus (De Hoff et al. 2013), *C. incerta* MT-is ~329 kb and *C. schloesseri* MT-is ~438 kb. In all, while we do find evidence of MT-haplotype reformation within *Chlamydomonas*, this is mostly limited to rearrangements, with far less gene turnover and MT size variation than has been observed between more distantly related core-*Reinhardtinia* species. While MT evolution has previously been explored in the context of transitions from unicellularity to multicellularity and isogamy to anisogamy, our data suggest that MT haplotype reformation is still expected to occur between closely related isogamous species, albeit at a reduced scale.

### Alignability and estimates of neutral divergence

In order to facilitate the identification of conserved elements (CEs), we produced an 8-species core-*Reinhardtinia* whole-genome alignment (WGA) using Cactus (Armstrong et al. 2019). Based on the alignment of *C. reinhardtii* four-fold degenerate (4D) sites extracted from the WGA, we estimated putatively neutral branch lengths across the topology connecting the eight species using the GTR substitution model (fig. 7a). Divergence between *C. reinhardtii* and *C. incerta,* and *C. reinhardtii* and *C. schloesseri*, was estimated as 34% and 45%, respectively. Divergence between *C. reinhardtii* and *E. debaryana* was estimated as 98%, while all four TGV clade species were saturated relative to *C. reinhardtii* (i.e. on average, each 4D site is expected to have experienced more than one substitution). To put these estimates within a more recognisable framework, divergence across *Chlamydomonas* is approximately on the scale of human-rodent divergence (Lindblad-Toh et al. 2011), while divergence between *Chlamydomonas* and the TGV clade is approximately equivalent to that of mammals and sauropsids (birds and reptiles), which diverged ~320 million years ago (Alföldi et al. 2011). Our estimates corroborate a previous estimate of synonymous divergence between *C. reinhardtii* and *C. incerta* of 37% (averaged over 67 orthologous genes) (Popescu et al. 2006), and are broadly in line with the divergence time estimate of ~230 million years ago between the TGV clade and their unicellular ancestors (Herron et al. 2009). Finally, it is important to note that we have likely underestimated neutral divergence, as 4D sites are unlikely to be evolving neutrally due to selection acting on codon usage, which has been shown to decrease divergence between *C. reinhardtii* and *C. incerta* (Popescu et al. 2006).

**Figure 7.**
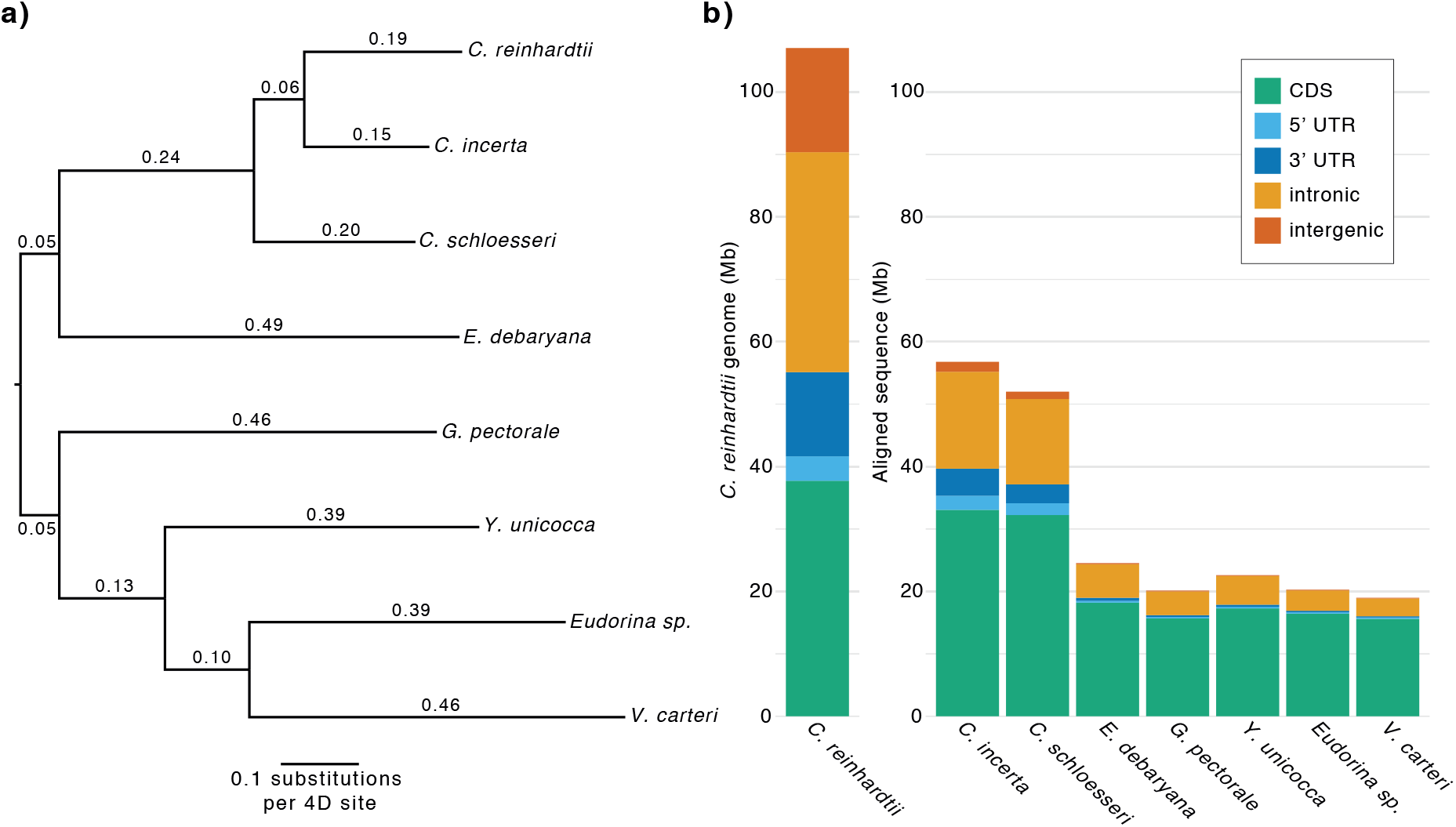
a) Estimates of putatively neutral divergence under the GTR model, based on the topology of figure 2 and 1,552,562 *C. reinhardtii* 4D sites extracted from the Cactus WGA. b) A representation of the *C. reinhardtii* genome by site class, and the number of aligned sites per *C. reinhardtii* site class for each other species in the Cactus WGA.

As expected, genome-wide alignability (i.e. the proportion of bases aligned between *C. reinhardtii* and a given species in the WGA) decreased substantially with increasing divergence, with 53.0% of the *C. reinhardtii* genome aligned to *C. incerta*, 48.6% to *C. schloesseri*, and on average only 19.9% to the remaining five species (fig. 7b). The majority of *C. reinhardtii* CDS was alignable within *Chlamydomonas* (87.7% and 85.5% to *C. incerta* and *C. schloesseri*, respectively), indicating that it will be possible to perform molecular evolutionary analyses on CDS between the three species. CDS also constituted the majority of the aligned sequence to the other five species, comprising on average 78.3% of the aligned bases despite CDS forming only 35.2% of the *C. reinhardtii* genome. In contrast, far less non-exonic sequence was alignable, especially at evolutionary distances beyond *Chlamydomonas*. Substantial proportions of intronic bases were aligned to *C. incerta* (44.1%) and *C. schloesseri* (38.8%), with on average 11.3% aligned to the other five species. Less than 10% of intergenic sequence was aligned to any one species, and on average less than 1% was aligned to non-*Chlamydomonas* species. Distributions of intergenic tract lengths across the core-*Reinhardtinia* are highly skewed (fig. S7), so that in *C. reinhardtii* tracts shorter than 250 bp constitute 63.5% of tracts but just 5.5% of total intergenic sequence. The sequence content of tracts >250 bp is highly repetitive (total repeat content 63.4%), while tracts <250 bp are relatively free of repeats (4.3% repeat content) and as a result are far more alignable to *C. incerta* and *C. schloesseri* (40.8% and 32.0% of bases aligned, respectively). This suggests that at least for introns and short intergenic tracts it is feasible to explore the landscape of non-exonic evolutionary constraint, primarily utilising alignment data from *Chlamydomonas*, supplemented by what is likely the alignment of only the most conserved sites at greater evolutionary distances.

### Missing genes in Chlamydomonas reinhardtii

One of the major successes of comparative genomics has been the refinement of gene annotations and identification of novel gene models and exons (e.g. Lin et al. (2008); Mudge et al. (2019)). Prior to identifying conserved sequences and classifying them as coding or noncoding, we attempted to identify novel *C. reinhardtii* genes absent from the latest annotation (v5.6) using patterns of *Chlamydomonas* synteny and the core-*Reinhardtinia* WGA. A *de novo C. reinhardtii* gene annotation yielded 433 novel gene models, 142 of which were retained based on the presence of a syntenic homolog in one or both of the *Chlamydomonas* species, and/or a phyloCSF (Lin et al. 2011) score >100. PhyloCSF assesses the protein-coding potential of a multi-species alignment using patterns of substitution at putative synonymous and non-synonymous sites, and so is not reliant on gene annotations in other species. Of the 142 supported genes, 90 had significant BLASTp hits (e-value <1×10^−5^, >=80% protein length) to *C. reinhardtii* proteins from annotation version 4.3 and likely represent models that were lost during the transition from v4 to v5 of the genome.

### The genomic landscape of sequence conservation in Chlamydomonas reinhardtii

Based on the core-*Reinhardtinia* WGA, we identified 265,006 CEs spanning 33.8 Mb or 31.5% of the *C. reinhardtii* genome. The majority of CE bases overlapped CDS (70.6%), with the remaining bases overlapping 5’ UTRs (2.9%), 3’ UTRs (4.4%), introns (20.0%) and intergenic sites (2.0%) (table 3). Relative to the site class categories themselves, 63.1% of CDS bases, 24.8% of 5’ UTR bases, 11.0% of 3’ UTR bases, and 19.2% of intronic bases were overlapped by CEs. Only 4.1% of intergenic bases were covered by CEs, however when splitting intergenic tracts into those <250 bp (short tracts) and >250 bp (long tracts), a more appreciable proportion of short tract bases (14.1%) were covered by CEs. As would be predicted given the expectation that CEs contain functional sequences, *C. reinhardtii* genetic diversity (π) was 39.5% lower for CEs (0.0134) than non-CE bases (0.0220), a result that was relatively consistent across site classes with the exception of long intergenic tracts (table 3). It is, however, important to state that the CEs we have identified are likely to contain a proportion of non-constrained sites. While this is always to be expected to some extent (e.g. CDS is generally included in CEs despite the presence of synonymous sites), given a mean length of 128 bp our CE dataset should be cautiously interpreted as regions containing elevated proportions of constrained sites.

**Table 3.** Overlap between conserved elements and *C. reinhardtii* genomic site classes.

| Site class | CE overlap (Mb) | Proportion of CE bases (%) | Proportion of site class (%) | Genetic diversity all sites ($\pi$) | Genetic diversity CE sites ($\pi$) | Genetic diversity non-CE sites ($\pi$) |
| --- | --- | --- | --- | --- | --- | --- |
| CDS | 23.85 | 70.64 | 63.10 | 0.0144 | 0.0112 | 0.0204 |
| 5' UTR | 0.97 | 2.86 | 24.76 | 0.0189 | 0.0138 | 0.0208 |
| 3' UTR | 1.48 | 4.38 | 10.97 | 0.0205 | 0.0151 | 0.0213 |
| intronic | 6.76 | 20.01 | 19.15 | 0.0248 | 0.0216 | 0.0256 |
| intergenic <250 bp | 0.13 | 0.38 | 14.07 | 0.0229 | 0.0194 | 0.0235 |
| intergenic >=250 bp | 0.56 | 1.65 | 3.55 | 0.0137 | 0.0134 | 0.0138 |

Given the compactness of the *C. reinhardtii* genome (82.1% genic, median intergenic tract length 134 bp), it is expected that a high proportion of regulatory sequence will be concentrated in UTRs and intergenic sequences immediately upstream of genes (i.e. promoter regions). Relatively little is known about the genome-wide distribution of regulatory elements in *C. reinhardtii*, although analyses based on motif modelling have identified putative *cis*-regulatory elements enriched in these regions (Ding et al. 2012; Hamaji et al. 2016a). Presumably many CEs overlapping UTRs and promoter regions harbour regulatory elements, and the CEs we have identified could be used in future studies to validate potential functional motifs. Due to our inability to align longer intergenic tracts, it remains an open question whether functional elements are present at non-negligible abundances in these regions. Although the lack of alignment could in itself be taken for a lack of constraint, the highly repetitive nature of these regions may disrupt the alignment of functional sequences present among repeats. It is noticeable that *C. reinhardtii* genetic diversity is lower in long intergenic tracts than all site classes except CDS (table 3), which could be due the presence of functional sequences or alternatively an as of yet unknown evolutionary mechanism.

All six annotated core-*Reinhardtinia* species contained conspicuously long introns (median lengths 198-343 bp, table 2). As reported previously for *C. reinhardtii* (Merchant et al. 2007), the distribution of intron lengths for core-*Reinhardtinia* species lacked the typical peak in intron lengths at 60-110 bp that is present in several model organisms with similarly compact genomes (fig. 8a, b). In *D. melanogaster*, short introns (<80 bp) appear to largely consist of neutrally evolving sequence, while longer introns that form the tail of the length distribution contain sequences evolving under evolutionary constraint (Halligan and Keightley 2006). To explore the relationship between intron length and sequence conservation in *C. reinhardtii*, we ordered introns by length and divided them into 50 bins, so that each bin contained an approximately equal number (~2,667) of introns. Mean intron length per bin was significantly negatively correlated with the proportion of bases overlapped by CEs (Pearson’s *r =* −0.626, p <0.01) (fig. 8c). This was particularly pronounced for introns <100 bp (~5% of introns), for which 48.1% of bases were overlapped by CEs, compared to 18.5% for longer introns. Therefore, it appears that in a reverse of the situation found in *D. melanogaster*, the minority of introns in *C. reinhardtii* are short and potentially functionally important, while the majority of introns are longer and contain far fewer conserved bases. The tight peak in the distribution of intron lengths combined with the lack of sequence constraint in *D. melanogaster* short introns led Halligan and Keightley (2006) to hypothesise that intron length was under selection, but not the intronic sequence itself, and that introns had essentially evolved to be as short as possible. It is possible that *C. reinhardtii* introns >100 bp are similarly evolving under selection to be bounded within certain length constraints, although the selective advantage of maintaining intron lengths substantially longer than the minimum remains unknown. Given that atypical intron length distributions are common to all core-*Reinhardtinia* species, whatever mechanism is driving intron length is likely to be evolutionarily ancient.

**Figure 8.**
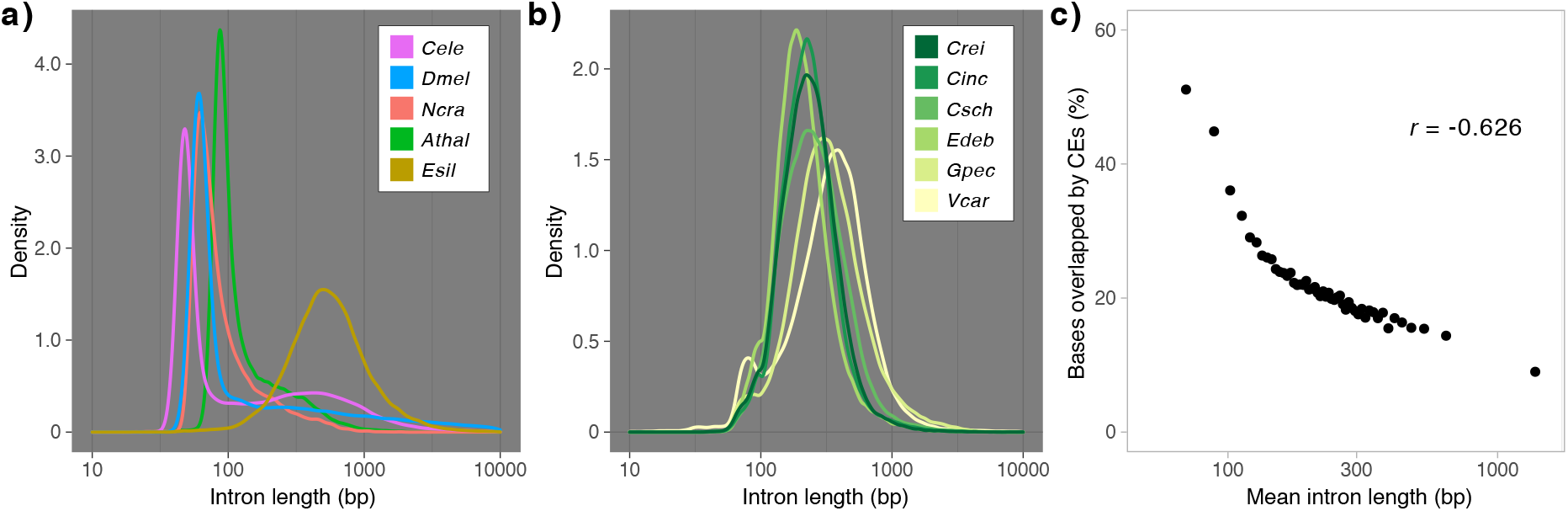
a) Intron length distributions for five model organisms (*Cele* = *C. elegans*, *Dmel* = *D. melanogaster*, *Ncra* = *Neurospora crassa*, *Athal* = *A. thaliana*, *Esil* = *Ectocarpus siliculosus*). The brown alga *E. siliculosus* is included as an example of an atypical distribution. b) Intron length distributions for six core-*Reinhardtinia* species, note different y-axis scale (*Crei* = *C. reinhardtii*, *Cinc* = *C. incerta*, *Csch* = *C. schloesseri*, *Edeb* = *Edaphochlamys debaryana*, *Gpec* = *G. pectorale*, *Vcar* = *V. carteri*). c). Correlation between mean intron length per bin and the proportion of bases overlapped by CEs. Introns were ordered by length and separated into 50 bins, so that each bin contained the same number of introns.

There are at least two leading explanations for why shorter *C. reinhardtii* introns may be functionally important. Firstly, intron retention (IR) has been shown to occur significantly more frequently in shorter genes (median = 181 bp) (Raj-Kumar et al. 2017). IR is the most common form of alternative splicing (AS) detected in *C. reinhardtii* (~30% of AS events), although AS in the species has not yet been extensively characterised and only ~1% of introns are currently annotated as alternatively retained. Furthermore, as only ~20% of IR events produce a functional protein (Raj-Kumar et al. 2017), not all retained introns are expected to be evolving under coding constraint. It is therefore difficult to assess the overall contribution of IR to intronic sequence conservation. Secondly, short introns may be enriched for regulatory sequences. Short introns <100 bp represent the first intron in a gene approximately four-fold more frequently (44.6%) than longer introns (10.3%) (fig. S8a). Introns <100 bp were also significantly more likely to occur closer to the transcription start site (mean intron positions relative to transcript length for introns <100 bp = 24.2% and introns >100 bp = 39.5%; independent-samples t-test t=-54.0, p<0.01) (fig. S8b). For many genes, introns within the first 1 kb have strong regulatory effects on gene expression (Rose 2018), and in *C. reinhardtii* it has been shown that the addition of a specific first intron to transgenes substantially increases their expression (Baier et al. 2018). It is also important to emphasise that a non-negligible proportion of sites in introns >100 bp are overlapped by CEs (18.5%), and thus longer introns are also likely to harbour some functional sites. Alongside regulatory sequences there are several possible explanations for this, including the presence of intronic RNA genes (Chen et al. 2008; Valli et al. 2016) and other categories of AS (e.g. alternative acceptor or donor splice sites) (Raj-Kumar et al. 2017).

Finally, we further identified 5,611 ultraconserved elements (UCEs) spanning 356.0 kb of the *C. reinhardtii* genome, defined as sequences >=50 bp exhibiting 100% sequence conservation across the three *Chlamydomonas* species. A subset of just 55 UCEs exhibited >=95% sequence conservation across all eight species, indicating that hardly any sequence is expected to be conserved to this level across the core-*Reinhardtinia*. The vast majority of UCE bases (96.0%) overlapped CDS, indicating constraint at both nonsynonymous and synonymous sites. There are several reasons why synonymous sites may be subject to such strong constraint, including interactions with RNA processing or formation of RNA secondary structures, the presence of exonic regulatory elements, or selection for optimal codon usage. Noticeably, 15 of the 55 core-*Reinhardtinia* UCEs overlapped ribosomal protein genes, which are often used as a standard for identifying optimal codons given their extremely high gene expression (Sharp and Li 1987), and several of the other genes overlapped by UCEs are also expected to be very highly expressed (e.g. elongation factors) (table S11). Although considered to be a very weak evolutionary force, this indicates that coordinated selection for optimal codons across the core-*Reinhardtinia* may be driving extreme sequence conservation. UCEs have proved to be excellent phylogenetic markers across several taxa (Faircloth et al. 2012; Faircloth et al. 2015). Given the lack of nuclear markers and the current difficulty in determining phylogenetic relationships in the core-*Reinhardtinia*, the 55 deeply conserved elements could potentially be used to provide additional phylogenetic resolution.

## Conclusions

Via the assembly of highly contiguous and well-annotated genomes for three of *C. reinhardtii*’s unicellular relatives, we have presented the first nucleotide-level comparative genomics framework for this important model organism. These resources are expected to enable the continued development of *C. reinhardtii* as a model system for molecular evolution. Furthermore, by providing insights into the gene content and genomic architecture of unicellular core-*Reinhardtinia* species, they are also expected to advance our understanding of the genomic changes that have occurred during the transition to multicellularity in the TGV clade.

Despite such advances, these genome assemblies have only now raised *C. reinhardtii* to a standard that had been achieved for many other model organisms ten or more years ago. Many of the analyses we have performed could be greatly enhanced by the inclusion of additional *Chlamydomonas* species, but addressing this is a question of taxonomy rather than sequencing effort. This is somewhat analogous to the past situation for *Caenorhabditis*, where only very recent advances in ecological knowledge have led to a rapid increase in the number of sampled species and sequenced genomes (Stevens et al. 2019). We hope that this study will encourage *Chlamydomonas* researchers to increase sampling efforts for new species, fully enabling the power of comparative genomics analyses to be realised for the species.

## Methods

### Nucleic acid extraction and sequencing

Isolates were obtained from the SAG or CCAP culture centres, cultured in Bold’s Basal Medium, and where necessary made axenic via serial dilution, plating on agar, and isolation of single algal colonies. High molecular weight DNA was extracted using a customised extension of an existing CTAB/phenol-chloroform protocol (file S1). One SMRTbell library (sheared to ~20 kb, with 15-50 kb size selection) was prepared per species, and each library was sequenced on a single SMRTcell on the PacBio Sequel platform. PacBio library preparation and sequencing were performed by Edinburgh Genomics.

DNA for Illumina sequencing was extracted using a phenol-chloroform protocol (Ness et al. 2012). Across all species a variety of library preparations, read lengths, insert sizes and sequencing platforms were used (table S2). RNA was extracted from four-day liquid cultures using Zymo Research TRI Reagent (product ID: R2050) and the Direct-zol RNA Miniprep Plus kit (product ID: R2070) following user instructions. One stranded RNA-seq library was prepared for each species using TruSeq reagents, and sequencing was performed on the Illumina HiSeq X platform (*Chlamydomonas incerta* 150 bp paired-end, *Chlamydomonas schloesseri* and *Edaphochlamys debaryana* 100 bp paired-end). All Illumina sequencing and library preparations were performed by BGI Hong Kong.

### De novo genome assembly

Detailed per-species methods and command line options are detailed in file S2. We first identified and removed reads derived from any contaminants by producing taxon-annotated GC-coverage plots with BlobTools v1.0 (Laetsch and Blaxter 2017a). Assemblies were produced using Canu v1.7.1 (Koren et al. 2017), with three iterative round of error-correction performed with the PacBio reads and the GenomicConsensus module Arrow v2.3.2 (https://github.com/PacificBiosciences/GenomicConsensus). All available Illumina data for each species was subsequently used to perform three iterative rounds of polishing using Pilon v1.22 (Walker et al. 2014). Assemblies of the plastid and mitochondrial genomes were produced independently and will be described elsewhere.

### Annotation of genes and repetitive elements

A preliminary repeat library was produced for each species with RepeatModeler v1.0.11 (Smit and Hubley 2008-2015). Repeat models with homology to *Chlamydomonas reinhardtii* v5.6 and/or *Volvox carteri* v2.1 transcripts (e-values <10^−3^, megablast (Camacho et al. 2009)) were filtered. The genomic abundance of each repeat model was estimated by providing RepeatMasker v4.0.9 (Smit et al. 2013-2015) with the filtered RepeatModeler output as a custom library, and any TEs with a cumulative total >100 kb were selected for manual curation, following Suh et al. (2014). Briefly, multiple copies of a given TE were retrieved by querying the appropriate reference genome using megablast, before each copy was extended at both flanks and aligned using MAFFT v7.245 (Katoh and Standley 2013). Alignments were then manually inspected, consensus sequences were created, and TE families were classified following Wicker et al. (2007) and Kapitonov and Jurka (2008). This procedure was also performed exhaustively for *C. reinhardtii* (i.e. curating all repeat models regardless of genomic abundance), which will be described in detail elsewhere. Final repeat libraries were made by combining the RepeatModeler output for a given species with all novel curated TEs and *V. carteri* repeats from Repbase (Bao et al. 2015) (files S3 and S4). TEs and satellites were soft-masked by providing RepeatMasker with the above libraries. In line with the most recent *C. reinhardtii* annotation (Blaby et al. 2014), low-complexity and simple repeats were not masked as the high GC-content of genuine coding sequence can result in excessive masking.

Adapters and low-quality bases were trimmed from each RNA-seq dataset using Trimmomatic v0.38 (Bolger et al. 2014) with the parameters optimised by Macmanes (2014). Trimmed reads were mapped to repeat-masked assemblies with the 2-pass mode of STAR v2.6.1a (Dobin et al. 2013). Gene annotation was performed with BRAKER v2.1.2 (Hoff et al. 2016; Hoff et al. 2019), an automated pipeline that combines the gene prediction tools Genemark-ET (Lomsadze et al. 2014) and AUGUSTUS (Stanke et al. 2006; Stanke et al. 2008). Read pairs mapping to the forward and reverse strands were extracted using samtools v1.9 (Li et al. 2009) and passed as individual BAM files to BRAKER, which was run with the “--UTR=on” and “--stranded=+,-” flags to perform UTR annotation. Resulting gene models were filtered for genes with internal stop codons, protein sequences <30 amino acids, or CDS overlapped by >=30% TEs/satellites or >=70% low-complexity/simple repeats.

Proteins were functionally annotated via upload to the Phycocosm algal genomics portal (https://phycocosm.jgi.doe.gov). Phycocosm uses an array of tools to add detailed annotation (gene ontology terms, Pfam domains, etc.), and additionally provides a genome browser interface to enable visualisation.

### Phylogenomics analyses

Genome and gene annotations for all available *Reinhardtinia* species and selected outgroups (tables S3, S4) were accessed from either Phytozome (if available) or NCBI. For annotation based analyses, protein clustering analysis was performed with OrthoFinder v2.2.7 (Emms and Kelly 2015), using the longest isoform for each gene, the modified BLASTp options “-seq yes, −soft_masking true, −use_sw_tback” (following Moreno-Hagelsieb and Latimer (2008)) and the default inflation value of 1.5. Protein sequences from orthogroups containing a single gene in all 11 included species (i.e. putative single copy-orthologs) were aligned with MAFFT and trimmed for regions of low-quality alignment using trimAl v1.4.rev15 (“-automated1”) (Capella-Gutiérrez et al. 2009). A ML species-tree was produced using concatenated gene alignments with IQ-TREE v1.6.9 (Nguyen et al. 2015), run with ModelFinder (“-m MFP”) (Kalyaanamoorthy et al. 2017) and ultrafast bootstrapping (“-bb 1000”) (Hoang et al. 2018). ASTRAL-III v5.6.3 (Zhang et al. 2018) was used to produce an alternative species-tree from individual gene-trees, which were themselves produced for each aligned single copy-ortholog using IQ-TREE as described above, with any branches with bootstrap support <10% contracted as recommended.

Annotation-free phylogenies were produced from a dataset of single-copy orthologous genes identified by BUSCO v3.0.2 (Waterhouse et al. 2018) run in genome mode with the pre-release Chlorophyta odb10 dataset (allowing missing data in up to three species). For each BUSCO gene, proteins were aligned and trimmed, and two species-trees were produced as described above.

### General comparative genomics and synteny analyses

Basic genome assembly metrics were generated using QUAST v5.0.0 (Gurevich et al. 2013). Repeat content was estimated by performing repeat masking on all genomes as described above (i.e. supplying RepeatMasker with the RepeatModeler output for a given species + manually curated repeats from all species). Assembly completeness was assessed by running BUSCO in genome mode with the Eukaryota odb9 and Chlorophyta odb10 datasets. Each species was run with *C. reinhardtii* (-sp chlamy2011) and *V. carteri* (-sp volvox) AUGUSTUS parameters, and the run with the most complete BUSCO genes was retained.

Synteny segments were identified between *C. reinhardtii* and the three novel genomes using SynChro (Drillon et al. 2014) with a block stringency value (delta) of 2. To create the input file for *C. reinhardtii*, we combined the repeat-filtered v5.6 gene annotation (see below) with the centromere locations for 15 of the 17 chromosomes, as defined by Lin et al. (2018). The resulting synteny blocks were used to check the *C. incerta* and *C. schloesseri* genomes for misassemblies, by manually inspecting breakpoints between synteny blocks on a given contig that resulted in a transition between *C. reinhardtii* chromosomes (see file S2). This resulted in four *C. incerta* and two *C. schloesseri* contigs being split due to likely misassembly.

A ML phylogeny of L1 LINE elements was produced from the endonuclease and reverse transcriptase domains (i.e. ORF2) of all known chlorophyte L1 elements. Protein sequences were aligned, trimmed and analysed with IQ-TREE as described above. All *C. incerta*, *C. schloesseri* and *E. debaryana* elements were manually curated as part of the annotation of repeats (see above). The *Yamagishiella unicocca, Eudorina sp.*, and *V. carteri* genomes were searched using tBLASTn with the L1-1_CR protein sequence as query, and the best hits were manually curated to assess the presence or absence of ZeppL elements in these species.

### Gene annotation metrics and gene family evolution

The *C. reinhardtii* v5.6 gene models were manually filtered based on overlap with the novel repeat library (files S3 and S4), which resulted in the removal of 1,085 putative TE genes. For all species, annotation completeness was assessed by protein mode BUSCO analyses using the Eukaryota odb9 and Chlorophyta odb10 datasets. Gene families were identified using OrthoFinder as described above with the six core-*Reinhardtinia* species with gene annotations (*C. reinhardtii*, *C. incerta*, *C. schloesseri*, *E. debaryana*, *G. pectorale* and *V. carteri*). Protein sequences for all species were annotated with InterPro domain IDs using InterProScan v5.39-77.0 (Jones et al. 2014). Domain IDs were assigned to orthogroups by KinFin v1.0 (Laetsch and Blaxter 2017b) if a particular ID was assigned to at least 20% of the genes and present in at least 50% of the species included in the orthogroup.

### Mating-type locus evolution

As the three novel genomes are all MT- and the *C. reinhardtii* reference genome is MT+, we first obtained the *C. reinhardtii* MT-locus and proteins from NCBI (accession GU814015.1) and created a composite chromosome 6 with an MT-haplotype. A reciprocal best hit approach with BLASTp was used to identify orthologs, supplemented with tBLASTn queries to search for genes not present in the annotations. To visualise synteny, we used the MCscan pipeline from the JCVI utility libraries v0.9.14 (Tang et al. 2008), which performs nucleotide alignment with LAST (Kiełbasa et al. 2011) to identify orthologs. We applied a C-score of 0.99, which filters LAST hits to only reciprocal best hits, while otherwise retaining default parameters. We manually confirmed that the LAST reciprocal hits were concordant with our BLASTp results. Scripts and data for this analysis are available at: https://github.com/aays/MT_analysis

### Core-Reinhardtinia whole-genome alignment and estimation of putatively neutral divergence

An 8-species core-*Reinhardtinia* WGA was produced using Cactus (Armstrong et al. 2019) with all available high-quality genomes (*C. reinhardtii* v5, *C. incerta*, *C. schloesseri*, *E. debaryana*, *Gonium pectorale, Y. unicocca, Eudorina sp.* and *V. carteri* v2). The required guide phylogeny was produced by extracting alignments of 4D sites from single-copy orthologs identified by BUSCO (genome mode, Chlorophyta odb10 dataset). Protein sequences of 1,543 BUSCO genes present in all eight species were aligned with MAFFT and subsequently back-translated to nucleotide sequences. Sites where the aligned codon in all eight species contained a 4D site were then extracted (250,361 sites), and a guide-phylogeny was produced by supplying the 4D site alignment and topology (extracted from the Volvocales species-tree, see above) to phyloFit (PHAST v1.4) (Siepel et al. 2005), which was run with default parameters (i.e. GTR substitution model).

Where available the R domain of the MT locus not included in a given assembly was appended as an additional contig (extracted from the following NCBI accessions: *C. reinhardtii* MT-GU814015.1, *G. pectorale* MT+ LC062719.1, *Y. unicocca* MT-LC314413.1, *Eudorina sp.* MT male LC314415.1, *V. carteri* MT male GU784916.1). All genomes were softmasked for repeats as described above, and Cactus was run using the guide-phylogeny and all genomes set as reference quality. Post-processing was performed by extracting a multiple alignment format (MAF) alignment with *C. reinhardtii* as the reference genome from the resulting hierarchical alignment (HAL) file, using the HAL tools command hal2maf (v2.1) (Hickey et al. 2013), with the options –onlyOrthologs and –noAncestors. Paralogous alignments were reduced to one sequence per species by retaining the sequence with the highest similarity to the consensus of the alignment block, using mafDuplicateFilter (mafTools suite v0.1) (Earl et al. 2014).

Final estimates of putatively neutral divergence were obtained using a method adopted from Green et al. (2014). For each *C. reinhardtii* protein-coding gene, the alignment of each exon was extracted and concatenated. For the subsequent CDS alignments, a site was considered to be 4D if the codon in *C. reinhardtii* included a 4D site, and all seven other species had a triplet of aligned bases that also included a 4D site at the same position (i.e. the aligned triplet was assumed to be a valid codon, based on its alignment to a *C. reinhardtii* codon). The resulting alignment of 1,552,562 sites were then passed to phyloFit with the species tree, as described above.

### Identification of novel Chlamydomonas reinhardtii genes

*De novo* gene annotation was performed on the *C. reinhardtii* v5 genome using BRAKER (without UTR annotation) and all RNA-seq datasets produced by Strenkert et al. (2019). Potential novel genes were defined as those without any overlap with CDS of v5.6 genes. To determine if any novel predictions had syntenic homologs within *Chlamydomonas*, SynChro was re-run against *C. incerta* and *C. schloesseri* using updated *C. reinhardtii* input files containing the potential novel genes. Coding potential was assessed by passing CDS alignments extracted from the WGA to phyloCSF (Lin et al. 2011), which was run in “omega” mode using the neutral branch length tree from phyloFit.

### Identification and analyses of conserved elements

CEs were identified from the 8-species WGA using phastCons (Siepel et al. 2005) with the phyloFit neutral model (described above) and the standard UCSC parameters “--expected-length=45, --target-coverage=0.3, --rho=0.31”. Parameter tuning was attempted, but it proved difficult to achieve a balance between overly long CEs containing too many non-constrained bases at one extreme, and overly fragmented CEs at the other, and the standard parameters were found to perform as adequately as others.

*C. reinhardtii* site classes were delineated using the repeat-filtered v5.6 annotation, augmented with the 142 novel genes identified (file S5). To assess the genomic distribution of conserved bases, site classes were called uniquely in a hierarchical manner, so that if a site was annotated as more than one site class it was called based on the following hierarchy: CDS, 5’ UTR, 3’ UTR, intronic, intergenic. Overlaps between site classes and CEs were calculated using BEDtools v2.26.0 (Quinlan and Hall 2010). Genetic diversity was calculated from re-sequencing data of 17 *C. reinhardtii* field isolates from Quebec, as described by Craig et al. (2019). For analyses of intron length and conservation, all introns were called based on longest isoforms as they appear in the annotation (i.e. no hierarchical calling was performed as described above).

## Supporting information

All supplementary files

## Supplementary files

supplementary_tables.xlsx

file_S1.pdf: high molecular weight DNA extraction protocol for *Chlamydomonas.*

file_S2.pdf: detailed genome assembly methods.

file_S3.fa: Volvocales curated TE library.

file_S4.xlsx: Volvocales curated TE annotation notes.

file_S5.gff3: *C. reinhardtii* v5.6 gene annotation, filtered for TE/repeat genes and with newly identified genes added.

file_S6.txt: OrthoFinder gene clustering used for phylogenomics analyses.

file_S7.fa: aligned and trimmed OrthoFinder single-copy orthologs used for phylogenomics analyses.

file_S8.nwk: IQ-TREE phylogeny produced from OrthoFinder single-copy orthologs.

file_S9.nwk: ASTRAL-III phylogeny produced from OrthoFinder single-copy orthologs.

file_S10.fa: aligned and trimmed chlorophyte BUSCO genes used for phylogenomics analyses.

file_S11.nwk: IQ-TREE phylogeny produced from chlorophyte BUSCO genes.

file_S12.nwk: ASTRAL-III phylogeny produced from chlorophyte BUSCO genes.

file_S13.tsv: *C. reinhardtii* – *C. incerta* synteny blocks.

file_S14.tsv: *C. reinhardtii* – *C. incerta* syntenic orthologs.

file_S15.tsv: *C. reinhardtii* – *C. schloesseri* synteny blocks.

file_S16.tsv: *C. reinhardtii* – *C. schloesseri* syntenic orthologs.

file_S17.tsv: *C. reinhardtii* – *E. debaryana* synteny blocks.

file_S18.tsv: *C. reinhardtii* – *E. debaryana* syntenic orthologs.

file_S19.fa: chlorophyte L1 LINE proteins.

file_S20.fa: aligned and trimmed chlorophyte L1 LINE proteins.

file_S21.nwk: IQ-TREE phylogeny of chlorophyte L1 LINE proteins.

file_S22.txt: OrthoFinder gene clustering of six core-*Reinhardtinia* species.

file_S23.txt: InterProScan raw output for genes of six core-*Reinhardtinia* species.

file_S24.tsv: InterPro domains associated with core-*Reinhardtinia* orthogroups.

file_S25.bed: phastCons conserved elements in *C. reinhardtii* v5 coordinates.

file_S26.bed: ultraconserved elements in *C. reinhardtii* v5 coordinates.

## Data availability

The 8-species core-*Reinhardtinia* Cactus WGA and all genome assemblies and annotations are available from the Edinburgh Datashare repository (doi: https://doi.org/10.7488/ds/2847). All sequencing reads, genome assemblies and gene annotations will shortly be available from NCBI under the BioProject PRJNA633871. Code and bioinformatic pipelines are available at: https://github.com/rorycraig337/Chlamydomonas_comparative_genomics

## Acknowledgements

We are indebted to Lewis Stevens, Dominik Laetsch and Mark Blaxter for their guidance and advice on all genomics matters. We thank Alexander Suh and Valentina Peona for their invaluable guidance on TE curation, Thomas Pröschold for providing isolate images and for useful discussions on taxonomy, and Olivier Vallon for useful discussions on chlorophyte genomics. We thank Susanne Kraemer and Jack Hearn for their work on an earlier version of the *C. incerta* genome. Rory Craig is supported by a BBSRC EASTBIO Doctoral Training Partnership grant. PacBio sequencing was funded by a NERC Biomolecular Analysis Facility Pilot Project Grant (NBAF1123).

## Author contributions

RJC performed analyses and wrote the first draft of the manuscript, with the exception of the analyses and manuscript section on mating-type evolution, which were performed and written by ARH. RJC and RWN performed laboratory work. RJC, RWN and PDK conceived the study. All authors read and commented on the final draft version of the manuscript.

---

**Figure S1.**
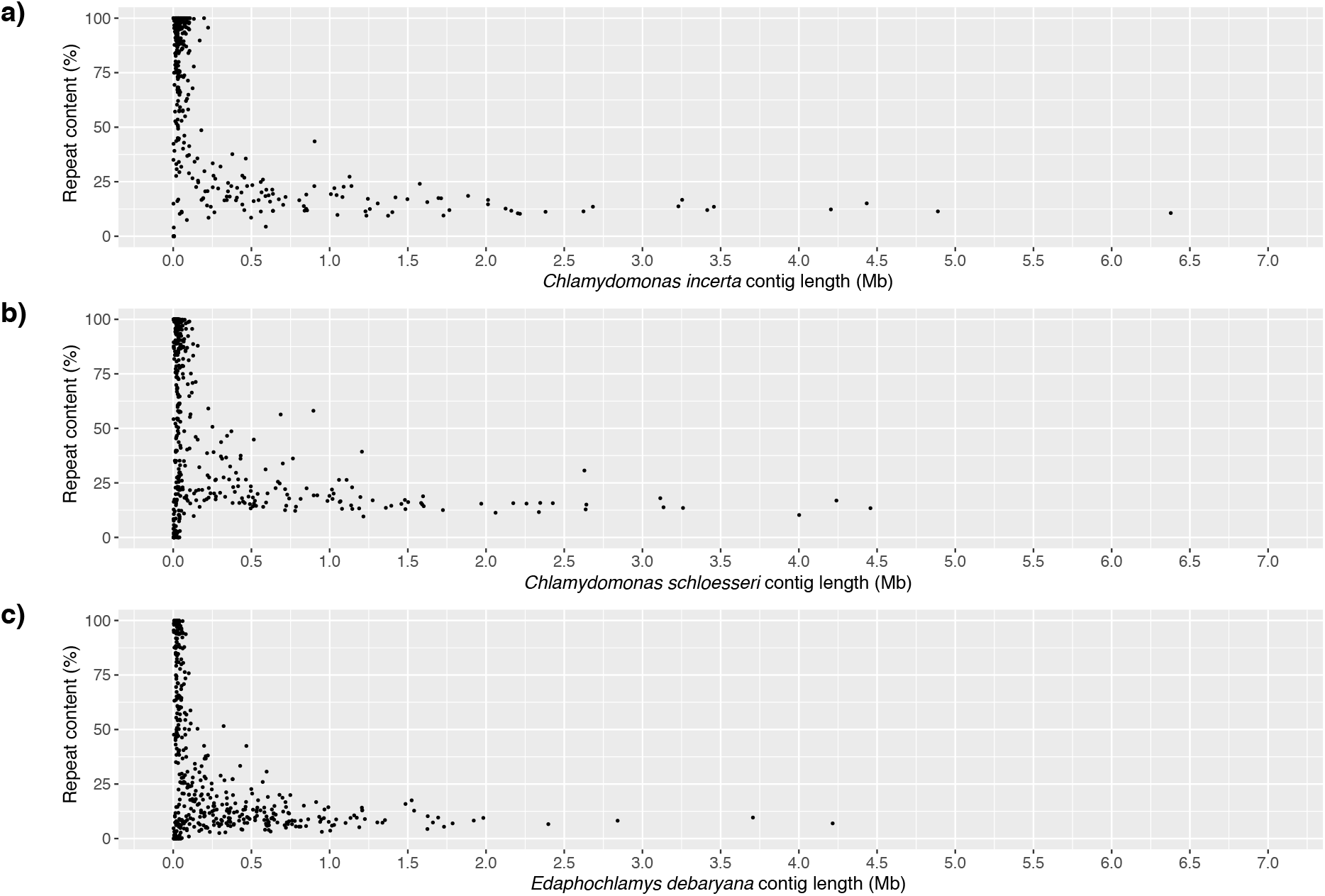
Total repeat content per contig (TEs, satellites and simple/low-complexity repeats) plotted by contig length for a) *Chlamydomonas incerta*, b) *Chlamydomonas schloesseri*, and c) *Edaphochlamys debaryana*.

**Figure S2.**
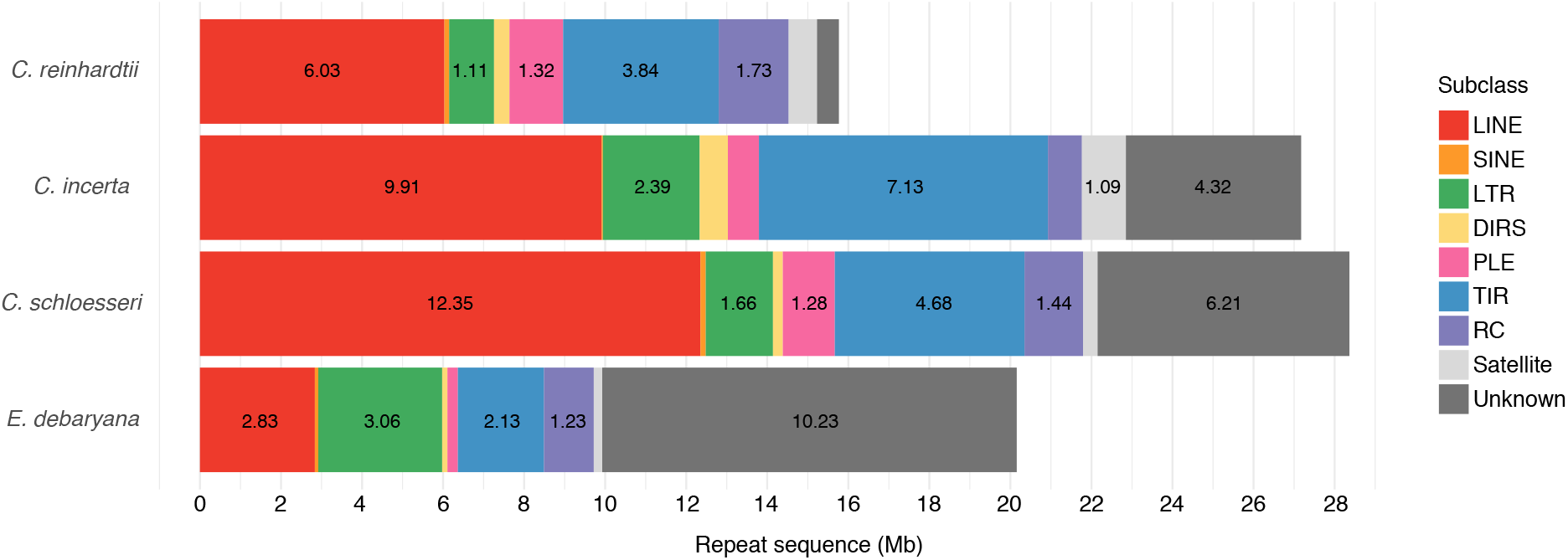
Repeat content per species by repeat subclass. Numbers within bars represent total sequence per subclass in Mb. LINE = long interspersed nuclear element, SINE = short interspersed nuclear element, LTR = long terminal repeat, DIRS = tyrosine recombinase encoding retrotransposons, PLE = Penelope-like elements, TIR = terminal inverted repeat (i.e. DNA transposons), RC = rolling-circle elements.

**Figure S3.**
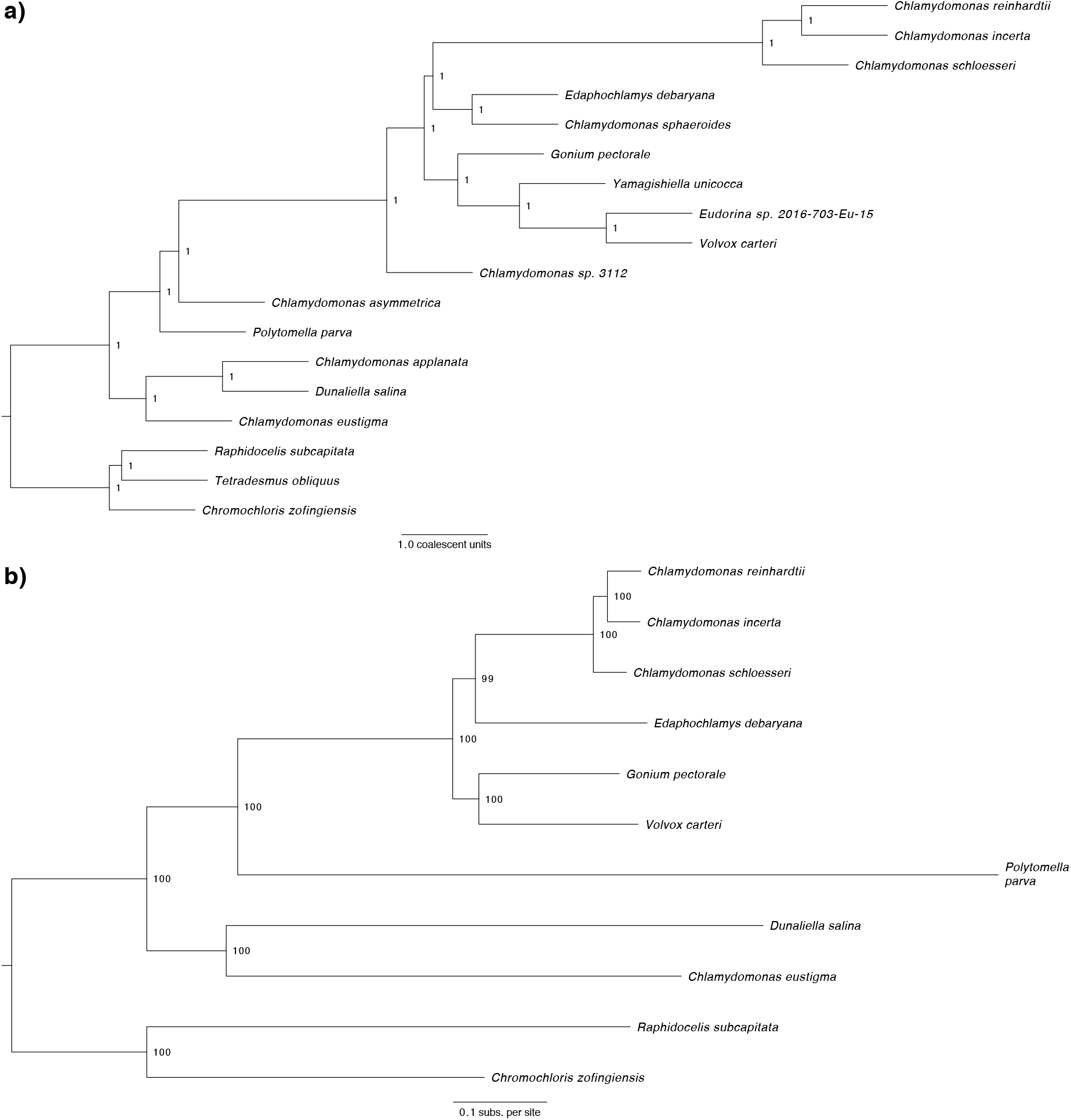
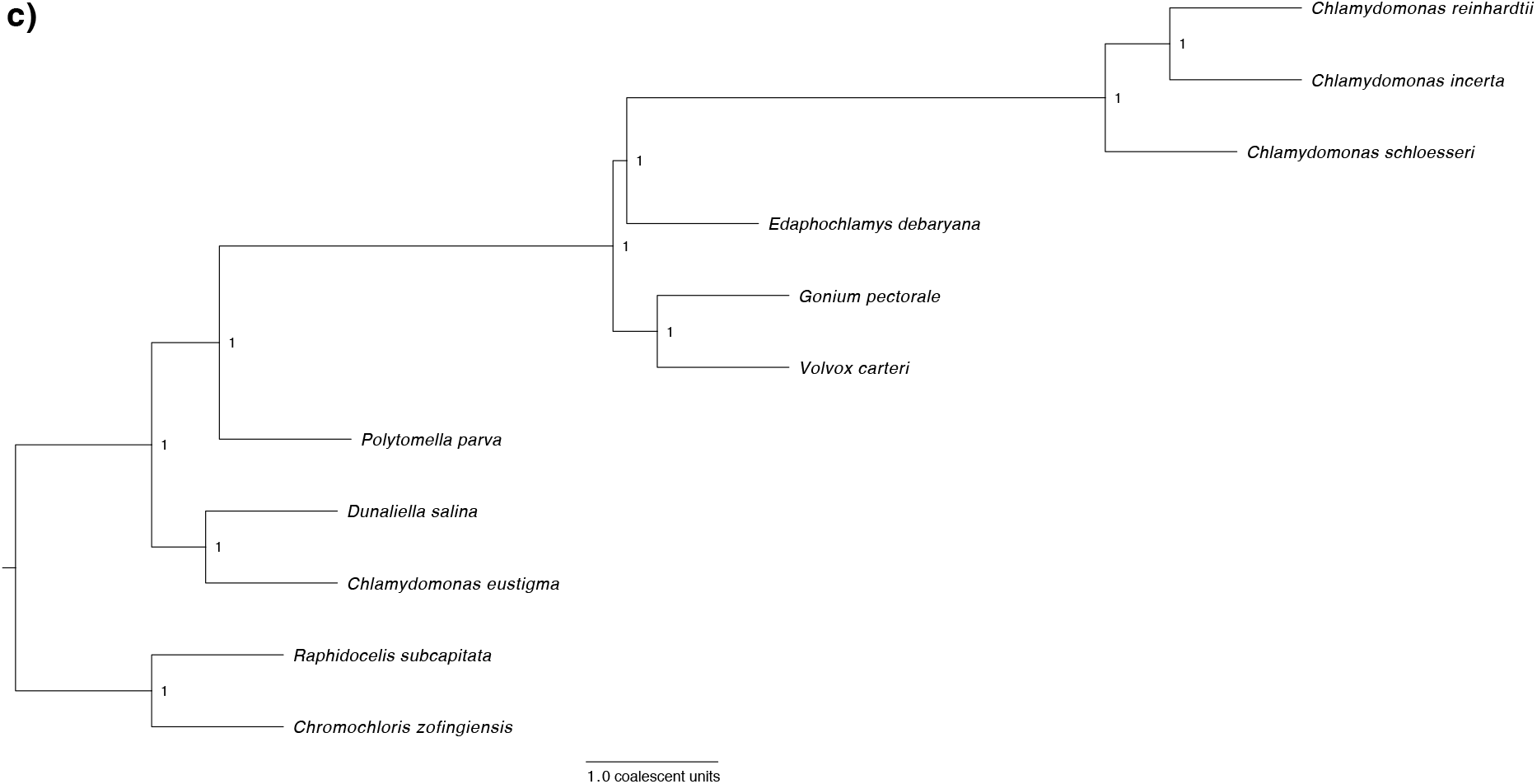
Phylogenomic analyses. a) ASTRAL-III species tree (15 Volvocales species and three outgroups) summarising 1,624 gene trees produced from individual protein alignments of chlorophyte BUSCO genes. b) ML phylogeny of nine Volvocales species and two outgroups inferred using LG+F+R5 model and a concatenated protein alignment of 1,681 putative single-copy orthologs identified by OrthoFinder. c) ASTRAL-III species tree summarising 1,681 gene trees produced from individual protein alignments of the OrthoFinder single-copy genes.

**Figure S4.**
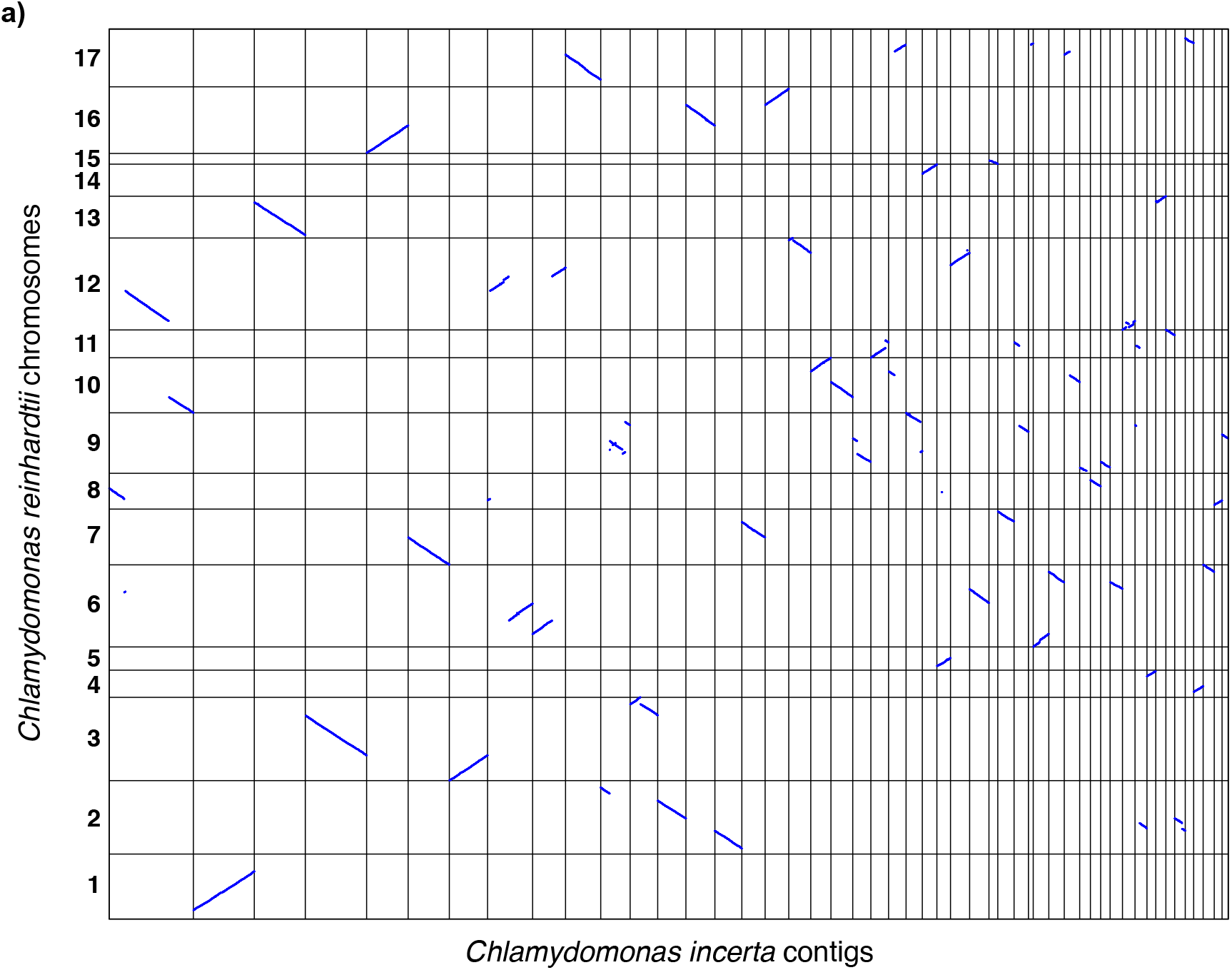
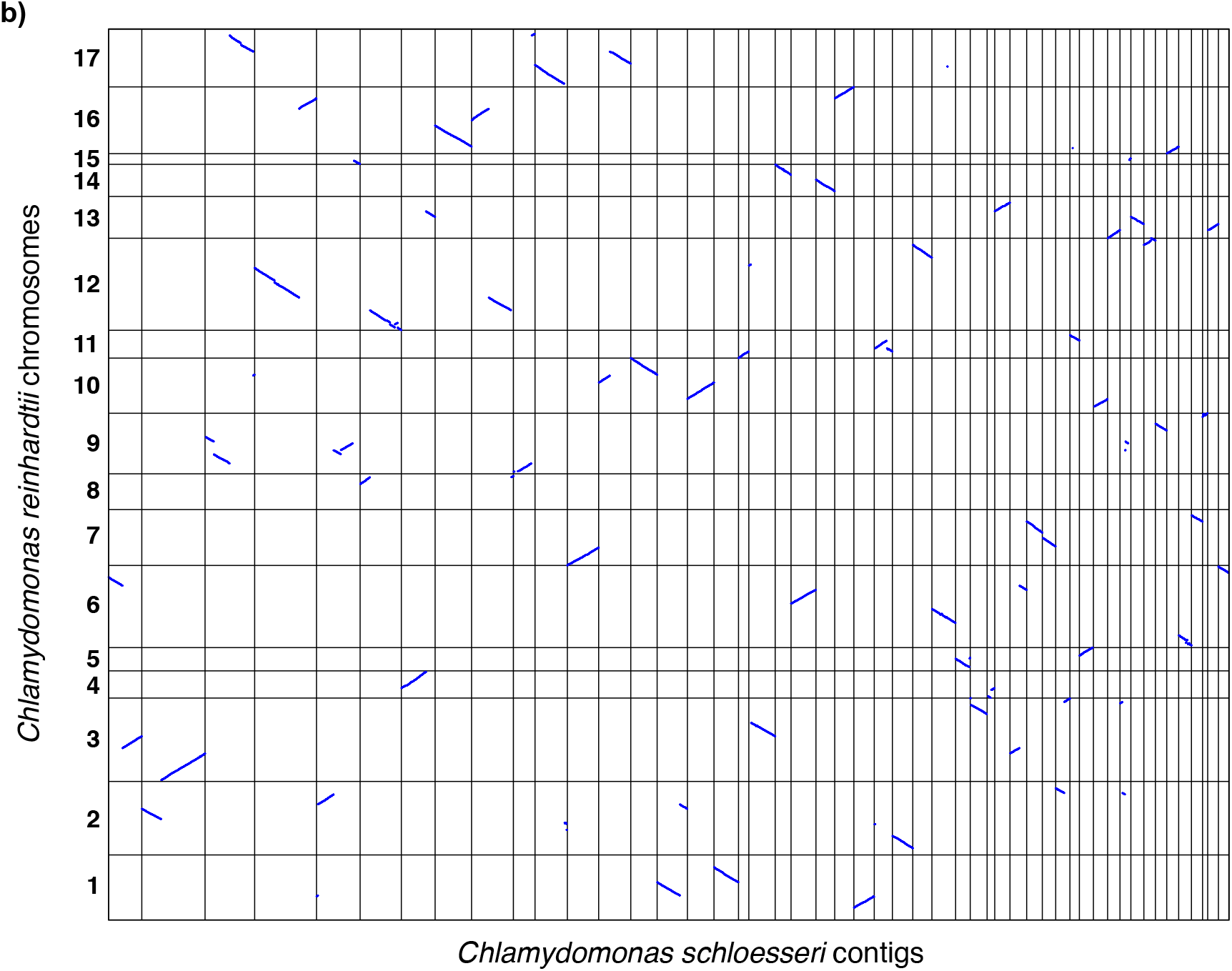
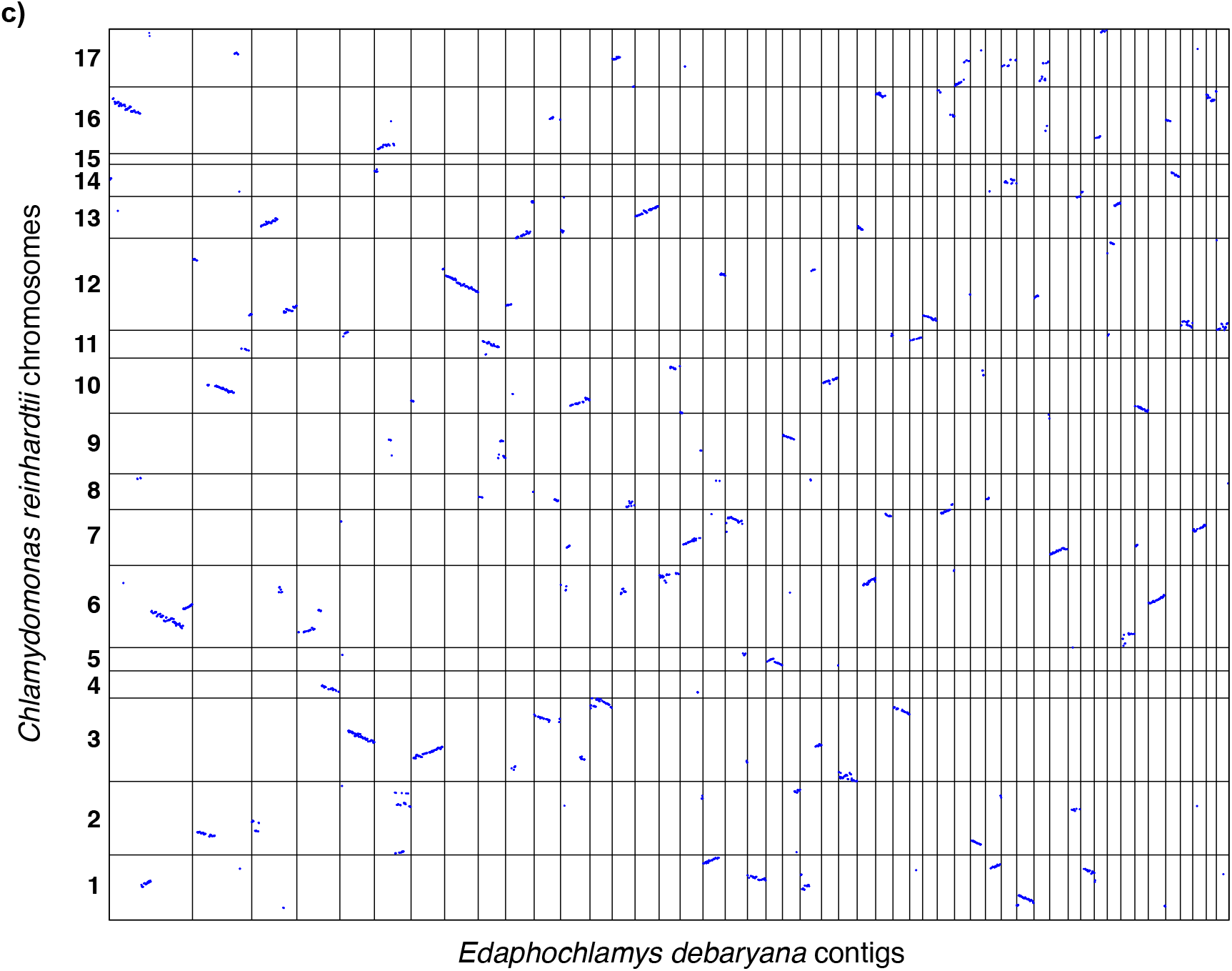
Dotplots representing syntenic genomic segments identified between *C. reinhardtii* and 50 largest contigs of a) *Chlamydomonas incerta*, b) *Chlamydomonas schloesseri*, and c) *Edaphochlamys debaryana*.

**Figure S5.**
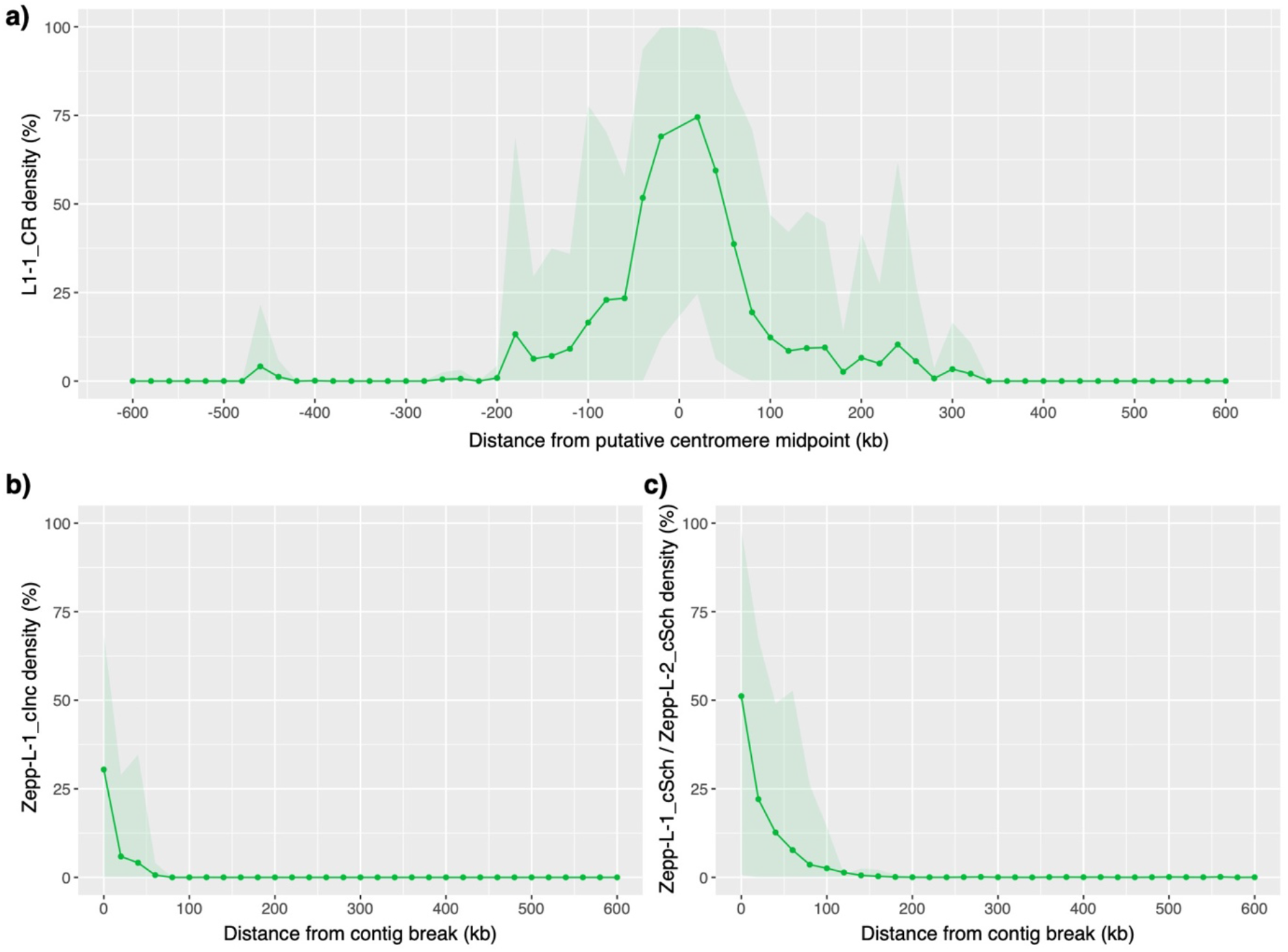
Mean densities of Zepp-like L1 LINE elements per 20 kb windows averaged over relevant chromosomes/contigs. Shaded areas represent 95% quantiles. a) Density of L1-1_CR elements relative to midpoint of 15 putative *C. reinhardtii* centromeres. b) Density of ZeppL-1_cInc elements relative to *C. incerta* contig ends syntenic to *C. reinhardtii* putative centromeres. c) Density of ZeppL-1_cSch and ZeppL-2_cSch elements relative to *C. schloesseri* contig ends syntenic to *C. reinhardtii* putative centromeres.

**Figure S6.**
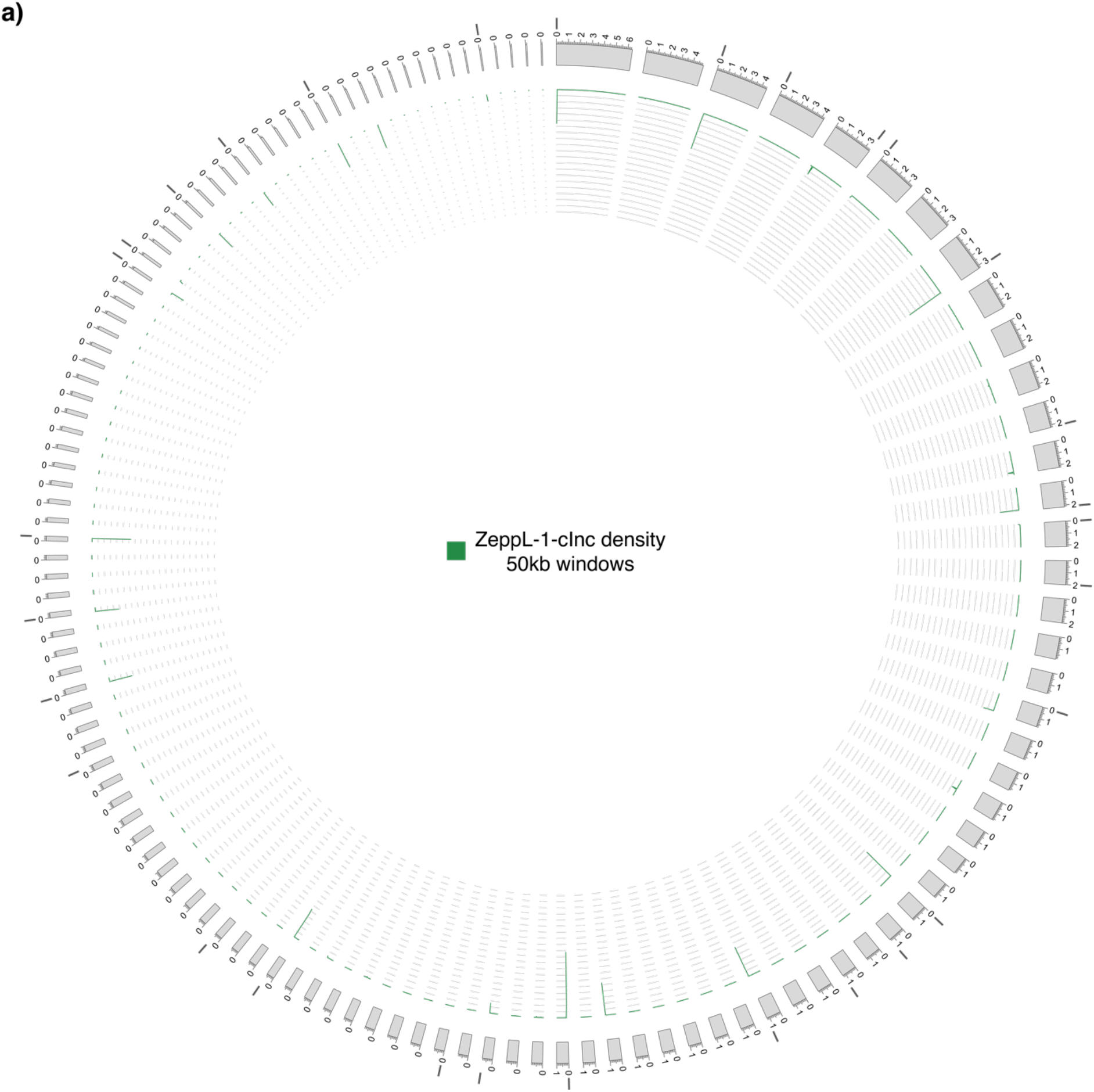
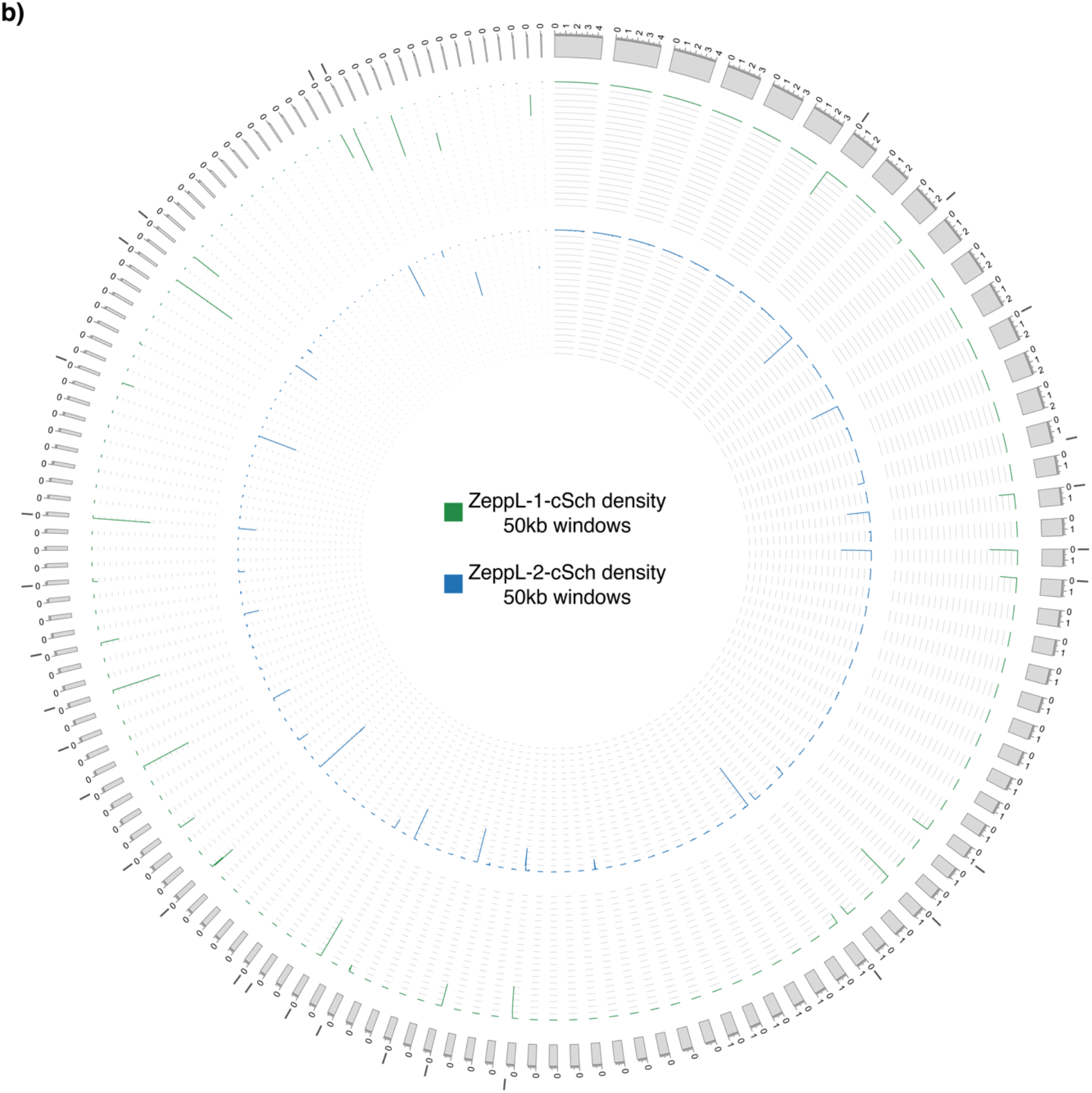
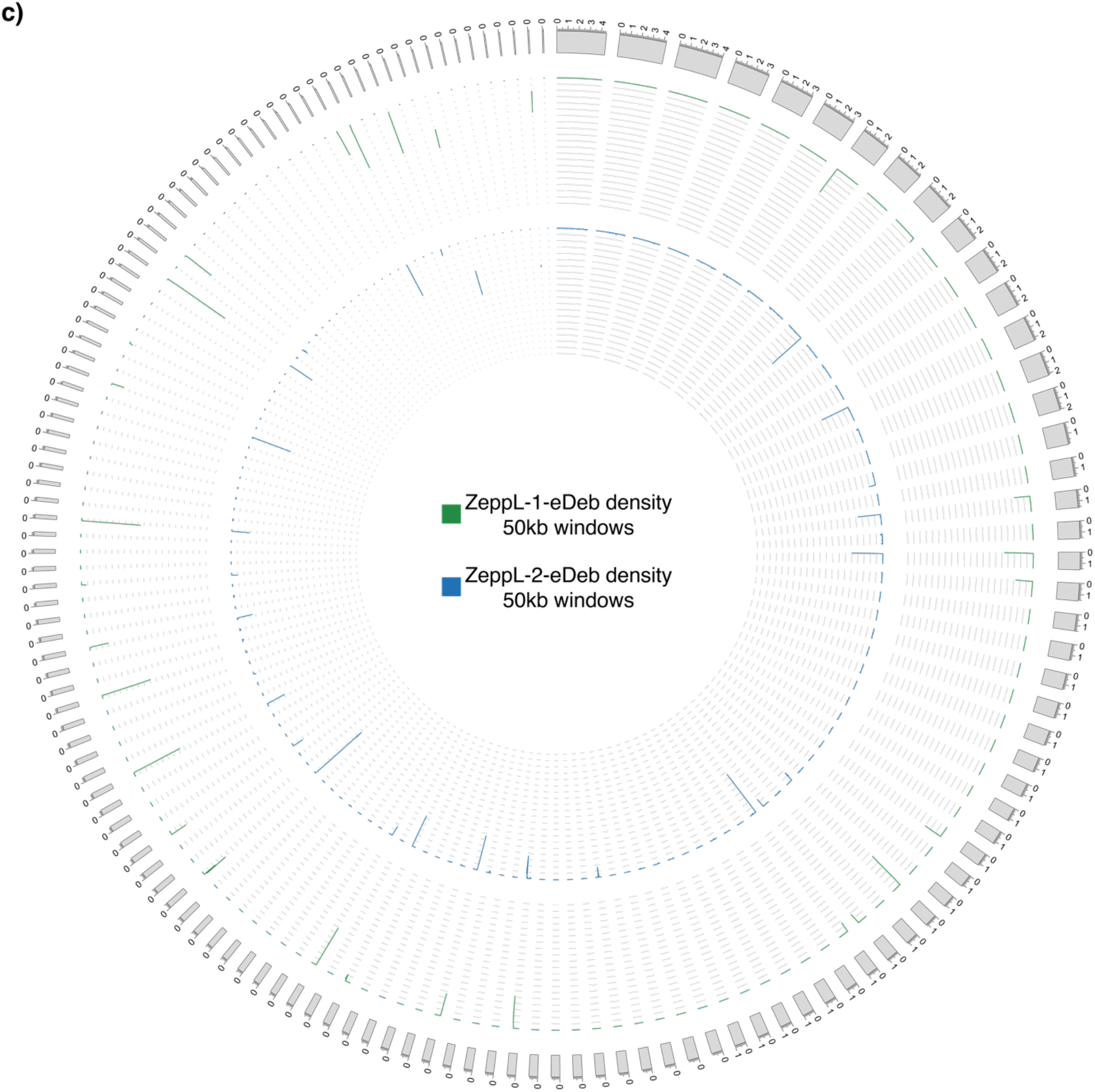
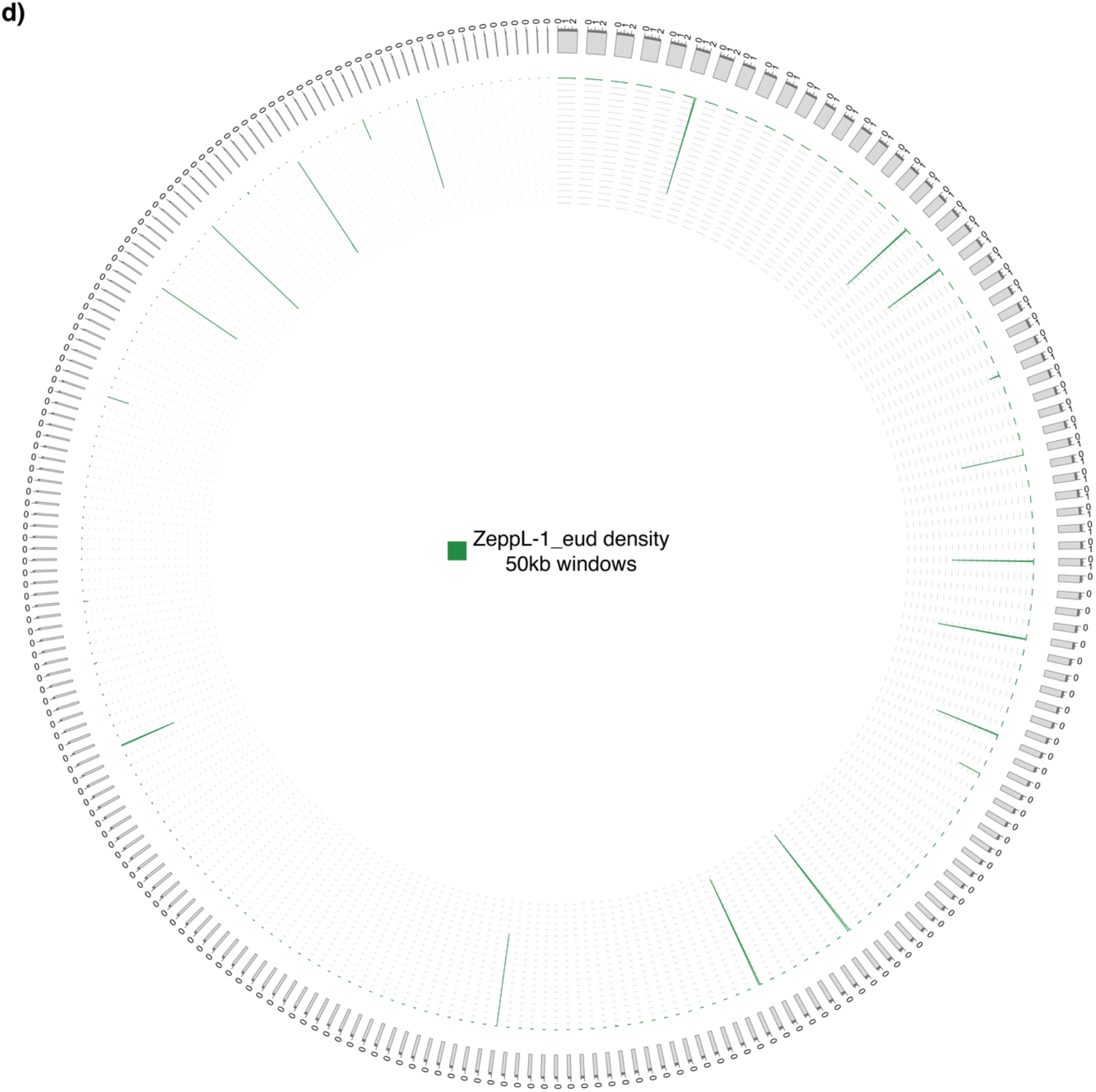
Genome-wide density of Zepp-like elements. Contigs are represented by grey bands and ordered by size. Dark grey bars above/below contigs represent contig ends inferred as syntenic with *C. reinhardtii* centromeres. Axis ranges from 0-100%. a) *Chlamydomonas incerta*. b) *Chlamydomonas schloesseri*. c) *Edaphochlamys debaryana*. d) *Eudorina sp. 2016-703-Eu-15*.

**Figure S7.**
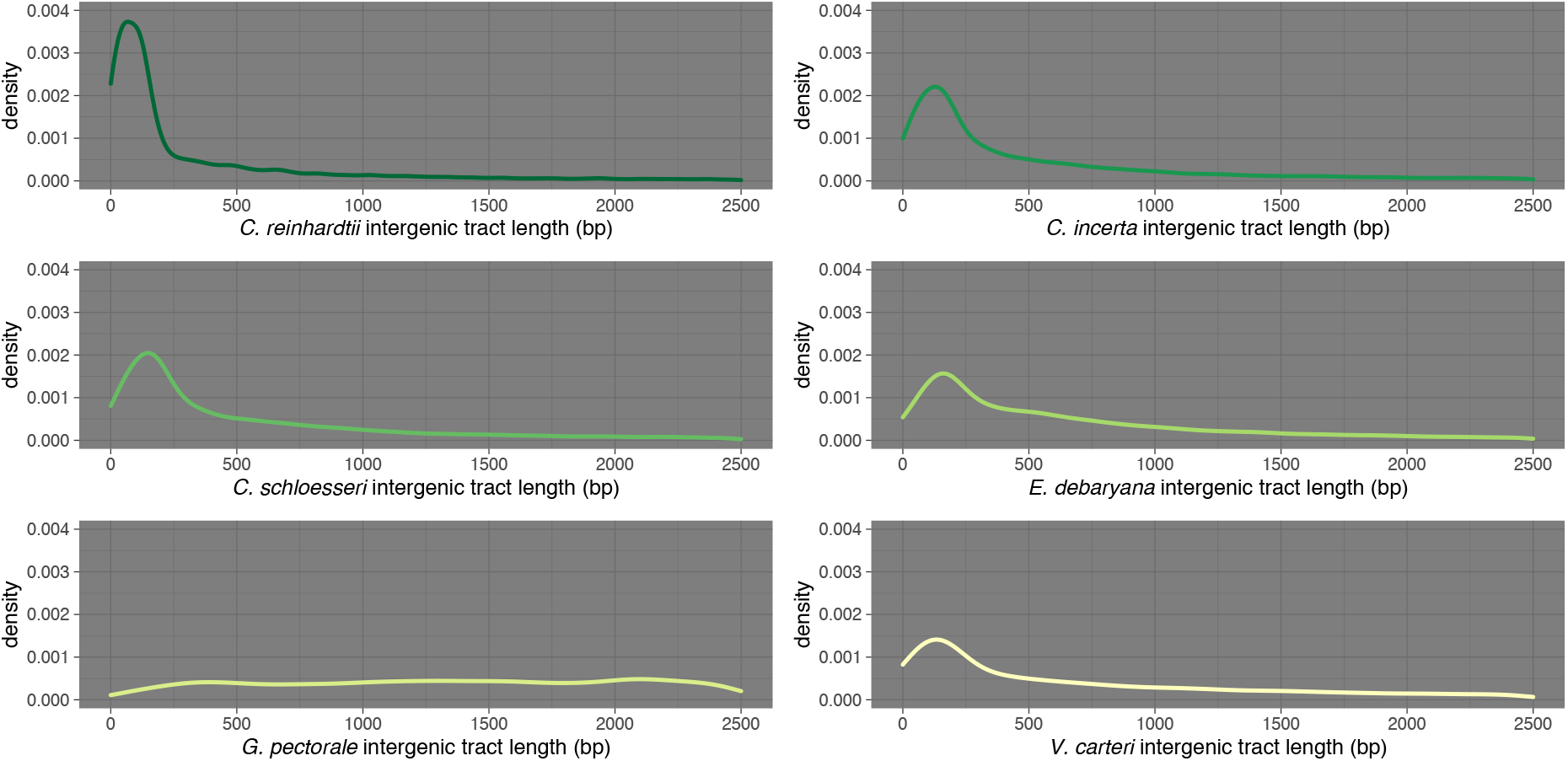
Distribution of intergenic tract lengths across six core-Reinhardtinia species. G. pectorale distribution differs due to the lack of UTR annotation for this species.

**Figure S8.**
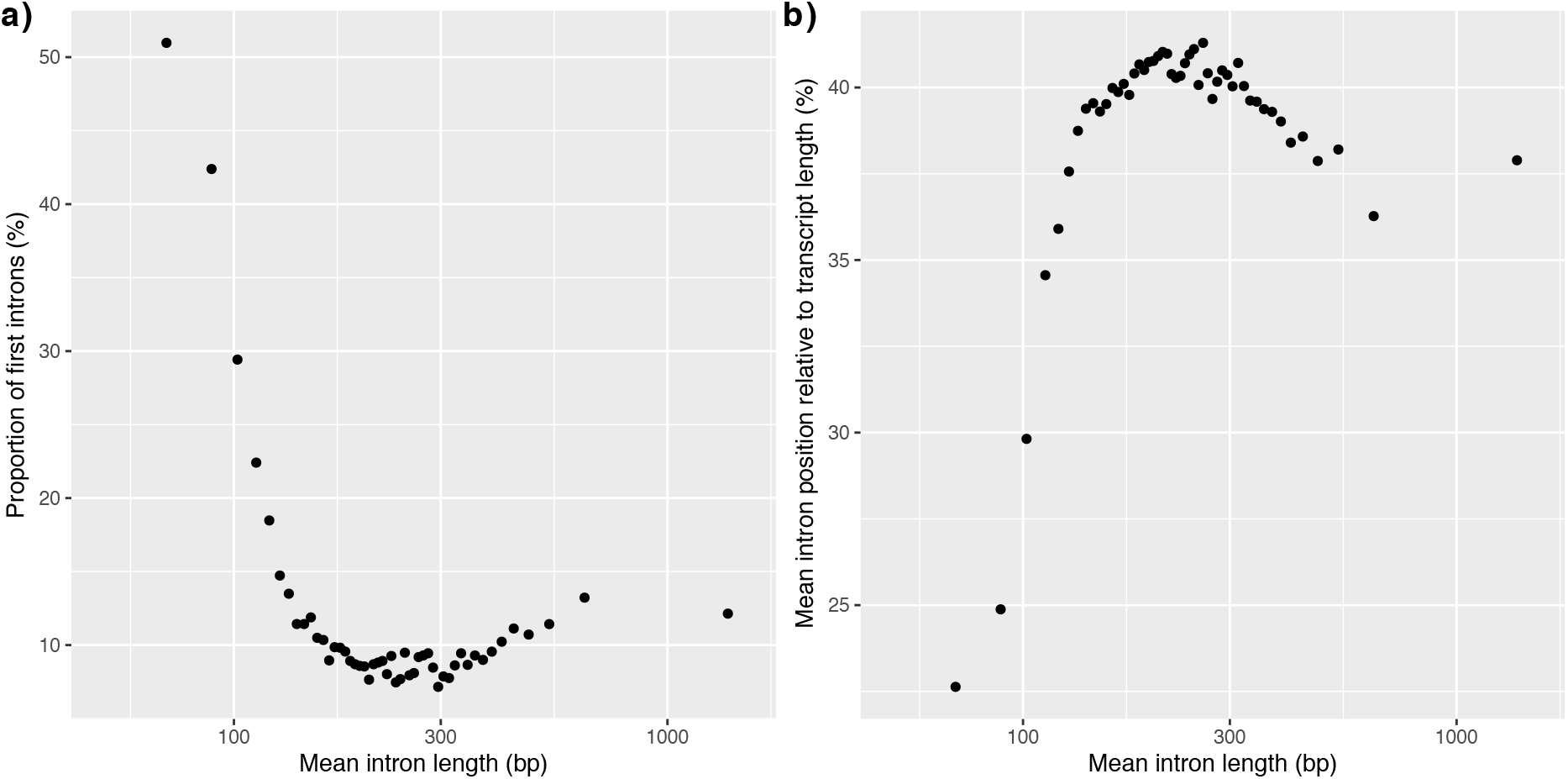
a) Relationship between the proportion of introns that are the first intron of a gene and the mean intron length per bin (see main text). b) The relationship between the mean intron position relative to transcript length (e.g. an intron at position 500 of a 2000 bp transcript equals 25%) and mean intron length per bin.

## Notes

### Competing Interest Statement

The authors have declared no competing interest.

## References

Alföldi J, Di Palma F, Grabherr M, Williams C, Kong LS, Mauceli E, Russell P, Lowe CB, Glor RE, Jaffe JD et al. 2011. The genome of the green anole lizard and a comparative analysis with birds and mammals. Nature 477: 587–591.

Alföldi J, Lindblad-Toh K. 2013. Comparative genomics as a tool to understand evolution and disease. Genome Res 23: 1063–1068.

Armstrong J, Hickey G, Diekhans M, Deran A, Fang Q, Xie D, Feng S, Stiller J, Genereux D, Johnson J et al. 2019. Progressive alignment with Cactus: a multiple-genome aligner for the thousand-genome era. bioRxiv.

Baier T, Wichmann J, Kruse O, Lauersen KJ. 2018. Intron-containing algal transgenes mediate efficient recombinant gene expression in the green microalga *Chlamydomonas reinhardtii*. Nucleic Acids Res 46: 6909–6919.

Bao W, Jurka J. 2013. Homologues of bacterial TnpB_IS605 are widespread in diverse eukaryotic transposable elements. Mob DNA 4: 12.

Bao W, Kojima KK, Kohany O. 2015. Repbase Update, a database of repetitive elements in eukaryotic genomes. Mobile DNA 6: 11.

Blaby IK, Blaby-Haas CE, Tourasse N, Hom EF, Lopez D, Aksoy M, Grossman A, Umen J, Dutcher S, Porter M et al. 2014. The *Chlamydomonas* genome project: a decade on. Trends in Plant Science 19: 672–680.

Blaby-Haas CE, Merchant SS. 2019. Comparative and functional algal genomics. Annual Review of Plant Biology 70: 605–638.

Blanc G, Agarkova I, Grimwood J, Kuo A, Brueggeman A, Dunigan DD, Gurnon J, Ladunga I, Lindquist E, Lucas S et al. 2012. The genome of the polar eukaryotic microalga *Coccomyxa subellipsoidea* reveals traits of cold adaptation. Genome Biol 13.

Böhne A, Zhou Q, Darras A, Schmidt C, Schartl M, Galiana-Arnoux D, Volff JN. 2012. Zisupton—a novel superfamily of DNA transposable elements recently active in fish. Mol Biol Evol 29: 631–645.

Bolger AM, Lohse M, Usadel B. 2014. Trimmomatic: a flexible trimmer for Illumina sequence data. Bioinformatics 30: 2114–2120.

Böndel KB, Kraemer SA, Samuels T, McClean D, Lachapelle J, Ness RW, Colegrave N, Keightley PD. 2019. Inferring the distribution of fitness effects of spontaneous mutations in *Chlamydomonas reinhardtii*. PLoS Biol 17: e3000192.

Camacho C, Coulouris G, Avagyan V, Ma N, Papadopoulos J, Bealer K, Madden TL. 2009. BLAST+: architecture and applications. BMC Bioinformatics 10: 421.

Capella-Gutiérrez S, Silla-Martínez JM, Gabaldón T. 2009. trimAl: a tool for automated alignment trimming in large-scale phylogenetic analyses. Bioinformatics 25: 1972–1973.

Chang CH, Chavan A, Palladino J, Wei XL, Martins NMC, Santinello B, Chen CC, Erceg J, Beliveau BJ, Wu CT et al. 2019. Islands of retroelements are major components of *Drosophila* centromeres. PLoS Biol 17.

Chen CL, Chen CJ, Vallon O, Huang ZP, Zhou H, Qu LH. 2008. Genomewide analysis of box C/D and box H/ACA snoRNAs in *Chlamydomonas reinhardtii* reveals an extensive organization into intronic gene clusters. Genetics 179: 21–30.

Cliften P, Sudarsanam P, Desikan A, Fulton L, Fulton B, Majors J, Waterston R, Cohen BA, Johnston M. 2003. Finding functional features in *Saccharomyces* genomes by phylogenetic footprinting. Science 301: 71–76.

Craig RJ, Böndel KB, Arakawa K, Nakada T, Ito T, Bell G, Colegrave N, Keightley PD, Ness RW. 2019. Patterns of population structure and complex haplotype sharing among field isolates of the green alga *Chlamydomonas reinhardtii*. Mol Ecol 28: 3977–3993.

Csuros M, Rogozin IB, Koonin EV. 2011. A detailed history of intron-rich eukaryotic ancestors inferred from a global survey of 100 complete genomes. PLoS Comput Biol 7.

De Hoff PL, Ferris P, Olson BJSC, Miyagi A, Geng S, Umen JG. 2013. Species and population level molecular profiling reveals cryptic recombination and emergent asymmetry in the dimorphic mating locus of *C. reinhardtii*. PLoS Genet 9.

Del Cortona A, Jackson CJ, Bucchini F, Van Bel M, D’hondt S, Skaloud P, Delwiche CF, Knoll AH, Raven JA, Verbruggen H et al. 2020. Neoproterozoic origin and multiple transitions to macroscopic growth in green seaweeds. Proc Natl Acad Sci U S A 117: 2551–2559.

Ding J, Li X, Hu H. 2012. Systematic prediction of cis-regulatory elements in the *Chlamydomonas reinhardtii* genome using comparative genomics. Plant Physiol 160: 613–623.

Dobin A, Davis CA, Schlesinger F, Drenkow J, Zaleski C, Jha S, Batut P, Chaisson M, Gingeras TR. 2013. STAR: ultrafast universal RNA-seq aligner. Bioinformatics 29: 15–21.

Drillon G, Carbone A, Fischer G. 2014. SynChro: a fast and easy tool to reconstruct and visualize synteny blocks along eukaryotic chromosomes. PLoS One 9: e92621.

Dvořáčková M, Fojtová M, Fajkus J. 2015. Chromatin dynamics of plant telomeres and ribosomal genes. Plant J 83: 18–37.

Earl D, Nguyen N, Hickey G, Harris RS, Fitzgerald S, Beal K, Seledtsov I, Molodtsov V, Raney BJ, Clawson H et al. 2014. Alignathon: a competitive assessment of whole-genome alignment methods. Genome Res 24: 2077–2089.

Emms DM, Kelly S. 2015. OrthoFinder: solving fundamental biases in whole genome comparisons dramatically improves orthogroup inference accuracy. Genome Biol 16: 157.

Faircloth BC, Branstetter MG, White ND, Brady SG. 2015. Target enrichment of ultraconserved elements from arthropods provides a genomic perspective on relationships among Hymenoptera. Mol Ecol Resour 15: 489–501.

Faircloth BC, McCormack JE, Crawford NG, Harvey MG, Brumfield RT, Glenn TC. 2012. Ultraconserved elements anchor thousands of genetic markers spanning multiple evolutionary timescales. Syst Biol 61: 717–726.

Fang Y, Coelho MA, Shu H, Schotanus K, Thimmappa BC, Yadav V, Chen H, Malc EP, Wang J, Mieczkowski PA et al. 2020. Long transposon-rich centromeres in an oomycete reveal divergence of centromere features in Stramenopila-Alveolata-Rhizaria lineages. PLoS Genet 16: e1008646.

Farlow A, Meduri E, Schlotterer C. 2011. DNA double-strand break repair and the evolution of intron density. Trends Genet 27: 1–6.

Featherston J, Arakaki Y, Hanschen ER, Ferris PJ, Michod RE, Olson B, Nozaki H, Durand PM. 2018. The 4-Celled *Tetrabaena socialis* nuclear genome reveals the essential components for genetic control of cell number at the origin of multicellularity in the volvocine lineage. Mol Biol Evol 35: 855–870.

Ferris P, Olson BJ, De Hoff PL, Douglass S, Casero D, Prochnik S, Geng S, Rai R, Grimwood J, Schmutz J et al. 2010. Evolution of an expanded sex-determining locus in *Volvox*. Science 328: 351–354.

Ferris PJ, Armbrust EV, Goodenough UW. 2002. Genetic structure of the mating-type locus of *Chlamydomonas reinhardtii*. Genetics 160: 181–200.

Ferris PJ, Goodenough UW. 1997. Mating type in *Chlamydomonas* is specified by *mid*, the minus-dominance gene. Genetics 146: 859–869.

Ferris PJ, Pavlovic C, Fabry S, Goodenough UW. 1997. Rapid evolution of sex-related genes in *Chlamydomonas*. Proc Natl Acad Sci U S A 94: 8634–8639.

Fulnečková J, Hasíková T, Fajkus J, Lukešová A, Eliáš M, Sýkorová E. 2012. Dynamic evolution of telomeric sequences in the green algal order Chlamydomonadales. Genome Biol Evo 4: 248–264.

Gel B, Serra E. 2017. karyoploteR: an R/Bioconductor package to plot customizable genomes displaying arbitrary data. Bioinformatics 33: 3088–3090.

Gerstein MB Lu ZJ Van Nostrand EL Cheng C Arshinoff BI Liu T Yip KY Robilotto R Rechtsteiner A Ikegami K et al. 2010. Integrative analysis of the *Caenorhabditis elegans* genome by the modENCODE project. Science 330: 1775–1787.

Green RE, Braun EL, Armstrong J, Earl D, Nguyen N, Hickey G, Vandewege MW, St John JA, Capella-Gutierrez S, Castoe TA et al. 2014. Three crocodilian genomes reveal ancestral patterns of evolution among archosaurs. Science 346: 1254449.

Grossman AR, Harris EE, Hauser C, Lefebvre PA, Martinez D, Rokhsar D, Shrager J, Silflow CD, Stern D, Vallon O et al. 2003. *Chlamydomonas reinhardtii* at the crossroads of genomics. Eukaryot Cell 2: 1137–1150.

Gurevich A, Saveliev V, Vyahhi N, Tesler G. 2013. QUAST: quality assessment tool for genome assemblies. Bioinformatics 29: 1072–1075.

Halligan DL, Keightley PD. 2006. Ubiquitous selective constraints in the *Drosophila* genome revealed by a genome-wide interspecies comparison. Genome Res 16: 875–884.

Halligan DL, Kousathanas A, Ness RW, Harr B, Eory L, Keane TM, Adams DJ, Keightley PD. 2013. Contributions of protein-coding and regulatory change to adaptive molecular evolution in murid rodents. PLoS Genet 9: e1003995.

Hamaji T, Kawai-Toyooka H, Uchimura H, Suzuki M, Noguchi H, Minakuchi Y, Toyoda A, Fujiyama A, Miyagishima S, Umen JG et al. 2018. Anisogamy evolved with a reduced sex-determining region in volvocine green algae. Communications Biology 1.

Hamaji T, Lopez D, Pellegrini M, Umen J. 2016a. Identification and characterization of a *cis*-regulatory element for zygotic gene expression in *Chlamydomonas reinhardtii*. G3 (Bethesda) 6: 1541–1548.

Hamaji T, Mogi Y, Ferris PJ, Mori T, Miyagishima S, Kabeya Y, Nishimura Y, Toyoda A, Noguchi H, Fujiyama A et al. 2016b. Sequence of the *Gonium pectorale* mating locus reveals a complex and dynamic history of changes in volvocine algal mating haplotypes. G3 (Bethesda) 6: 1179–1189.

Hanschen ER, Marriage TN, Ferris PJ, Hamaji T, Toyoda A, Fujiyama A, Neme R, Noguchi H, Minakuchi Y, Suzuki M et al. 2016. The *Gonium pectorale* genome demonstrates co-option of cell cycle regulation during the evolution of multicellularity. Nat Commun 7: 11370.

Harris EH, Boynton JE, Gillham NW, Burkhart BD, Newman SM. 1991. Chloroplast genome organization in *Chlamydomonas*. Arch Protistenkd 139: 183–192.

Hasan AR, Duggal JK, Ness RW. 2019. Consequences of recombination for the evolution of the mating type locus in *Chlamydomonas reinhardtii*. New Phytologist doi:10.1111/nph.16003.

Hasan AR, Ness RW. 2020. Recombination rate variation and infrequent sex influence genetic diversity in *Chlamydomonas reinhardtii*. Genome Biol Evol doi:10.1093/gbe/evaa057.

Herron MD, Hackett JD, Aylward FO, Michod RE. 2009. Triassic origin and early radiation of multicellular volvocine algae. Proc Natl Acad Sci U S A 106: 3254–3258.

Hickey G, Paten B, Earl D, Zerbino D, Haussler D. 2013. HAL: a hierarchical format for storing and analyzing multiple genome alignments. Bioinformatics 29: 1341–1342.

Higashiyama T, Noutoshi Y, Fujie M, Yamada T. 1997. Zepp, a LINE-like retrotransposon accumulated in the *Chlorella* telomeric region. EMBO J 16: 3715–3723.

Hiller M, Agarwal S, Notwell JH, Parikh R, Guturu H, Wenger AM, Bejerano G. 2013. Computational methods to detect conserved non-genic elements in phylogenetically isolated genomes: application to zebrafish. Nucleic Acids Res 41: e151.

Hirashima T, Tajima N, Sato N. 2016. Draft genome sequences of four species of *Chlamydomonas* containing phosphatidylcholine. Microbiol Resour Ann 4.

Hoang DT, Chernomor O, von Haeseler A, Minh BQ, Vinh LS. 2018. UFBoot2: improving the ultrafast bootstrap approximation. Mol Biol Evol 35: 518–522.

Hoff KJ, Lange S, Lomsadze A, Borodovsky M, Stanke M. 2016. BRAKER1: unsupervised RNA-Seq-based genome annotation with GeneMark-ET and AUGUSTUS. Bioinformatics 32: 767–769.

Hoff KJ, Lomsadze A, Borodovsky M, Stanke M. 2019. Whole-genome annotation with BRAKER. Methods in Molecular Biology 1962: 65–95.

Howell SH. 1972. The differential synthesis and degradation of ribosomnal DNA during the vegetative cell-cycle in *Chlamydomonas reinhardi*. Nature New Biology 240: 264–267.

Iyer LM, Zhang DP, de Souza RF, Pukkila PJ, Rao A, Aravind L. 2014. Lineage-specific expansions of TET/JBP genes and a new class of DNA transposons shape fungal genomic and epigenetic landscapes. Proc Natl Acad Sci U S A 111: 1676–1683.

Jones P, Binns D, Chang HY, Fraser M, Li W, McAnulla C, McWilliam H, Maslen J, Mitchell A, Nuka G et al. 2014. InterProScan 5: genome-scale protein function classification. Bioinformatics 30: 1236–1240.

Kalyaanamoorthy S, Minh BQ, Wong TKF, von Haeseler A, Jermiin LS. 2017. ModelFinder: fast model selection for accurate phylogenetic estimates. Nat Methods 14: 587–589.

Kapitonov VV, Jurka J. 2008. A universal classification of eukaryotic transposable elements implemented in Repbase. Nat Rev Genet 9: 411–412; author reply 414.

Katoh K, Standley DM. 2013. MAFFT multiple sequence alignment software version 7: improvements in performance and usability. Mol Biol Evol 30: 772–780.

Keightley PD, Jackson BC. 2018. Inferring the probability of the derived vs. the ancestral allelic state at a polymorphic Site. Genetics 209: 897–906.

Kiełbasa SM, Wan R, Sato K, Horton P, Frith MC. 2011. Adaptive seeds tame genomic sequence comparison. Genome Res 21: 487–493.

Kojima KK, Fujiwara H. 2005. An extraordinary retrotransposon family encoding dual endonucleases. Genome Res 15: 1106–1117.

Koren S, Walenz BP, Berlin K, Miller JR, Bergman NH, Phillippy AM. 2017. Canu: scalable and accurate long-read assembly via adaptive k-mer weighting and repeat separation. Genome Res 27: 722–736.

Krzywinski M, Schein J, Birol I, Connors J, Gascoyne R, Horsman D, Jones SJ, Marra MA. 2009. Circos: an information aesthetic for comparative genomics. Genome Res 19: 1639–1645.

Laetsch DR, Blaxter M. 2017a. BlobTools: interrogation of genome assemblies. F1000Research 6.

Laetsch DR, Blaxter ML. 2017b. KinFin: software for taxon-aware analysis of clustered protein sequences. G3 (Bethesda) 7: 3349–3357.

Lex A, Gehlenborg N, Strobelt H, Vuillemot R, Pfister H. 2014. UpSet: visualization of intersecting sets. IEEE Trans Vis Comput Graph 20: 1983–1992.

Li H, Handsaker B, Wysoker A, Fennell T, Ruan J, Homer N, Marth G, Abecasis G, Durbin R, Genome Project Data Processing S. 2009. The Sequence Alignment/Map format and SAMtools. Bioinformatics 25: 2078–2079.

Lin H, Cliften PF, Dutcher SK. 2018. MAPINS, a highly efficient detection method that identifies insertional mutations and complex DNA rearrangements. Plant Physiol 178: 1436–1447.

Lin MF, Deoras AN, Rasmussen MD, Kellis M. 2008. Performance and scalability of discriminative metrics for comparative gene identification in 12 *Drosophila* genomes. PLoS Comput Biol 4: e1000067.

Lin MF, Jungreis I, Kellis M. 2011. PhyloCSF: a comparative genomics method to distinguish protein coding and non-coding regions. Bioinformatics 27: i275–282.

Lindblad-Toh K, Garber M, Zuk O, Lin MF, Parker BJ, Washietl S, Kheradpour P, Ernst J, Jordan G, Mauceli E et al. 2011. A high-resolution map of human evolutionary constraint using 29 mammals. Nature 478: 476–482.

Liu H, Huang J, Sun X, Li J, Hu Y, Yu L, Liti G, Tian D, Hurst LD, Yang S. 2018. Tetrad analysis in plants and fungi finds large differences in gene conversion rates but no GC bias. Nat Ecol Evol 2: 164–173.

Lomsadze A, Burns PD, Borodovsky M. 2014. Integration of mapped RNA-Seq reads into automatic training of eukaryotic gene finding algorithm. Nucleic Acids Res 42: e119.

Lowe CB, Kellis M, Siepel A, Raney BJ, Clamp M, Salama SR, Kingsley DM, Lindblad-Toh K, Haussler D. 2011. Three periods of regulatory innovation during vertebrate evolution. Science 333: 1019–1024.

Macmanes MD. 2014. On the optimal trimming of high-throughput mRNA sequence data. Frontiers in Genetics 5: 13.

Marco Y, Rochaix JD. 1980. Organization of the nuclear ribosomal DNA of *Chlamydomonas reinhardii*. Mol Gen Genet 177: 715–723.

Margulies EH, Chen CW, Green ED. 2006. Differences between pair-wise and multi-sequence alignment methods affect vertebrate genome comparisons. Trends Genet 22: 187–193.

Merchant SS Prochnik SE Vallon O Harris EH Karpowicz SJ Witman GB Terry A Salamov A Fritz-Laylin LK Marechal-Drouard L et al. 2007. The *Chlamydomonas* genome reveals the evolution of key animal and plant functions. Science 318: 245–250.

Mikkelsen TS, Wakefield MJ, Aken B, Amemiya CT, Chang JL, Duke S, Garber M, Gentles AJ, Goodstadt L, Heger A et al. 2007. Genome of the marsupial *Monodelphis domestica* reveals innovation in non-coding sequences. Nature 447: 167–177.

Moreno-Hagelsieb G, Latimer K. 2008. Choosing BLAST options for better detection of orthologs as reciprocal best hits. Bioinformatics 24: 319–324.

Mouse Genome Sequencing Consortium. 2002. Initial sequencing and comparative analysis of the mouse genome. Nature 420: 520–562.

Mudge JM, Jungreis I, Hunt T, Gonzalez JM, Wright JC, Kay M, Davidson C, Fitzgerald S, Seal R, Tweedie S et al. 2019. Discovery of high-confidence human protein-coding genes and exons by whole-genome PhyloCSF helps elucidate 118 GWAS loci. Genome Res 29: 2073–2087.

Nakada T, Ito T, Tomita M. 2016. 18S ribosomal RNA gene phylogeny of a colonial volvocalean lineage (*Tetrabaenaceae-Goniaceae-Volvocaceae, Volvocales, Chlorophyceae*) and its close relatives. The Journal of Japanese Botany 91: 345–354.

Nakada T, Misawa K, Nozaki H. 2008. Molecular systematics of Volvocales (Chlorophyceae, Chlorophyta) based on exhaustive 18S rRNA phylogenetic analyses. Molecular Phylogenetics and Evolution 48: 281–291.

Nakada T, Tsuchida Y, Tomita M. 2019. Improved taxon sampling and multigene phylogeny of unicellular chlamydomonads closely related to the colonial volvocalean lineage Tetrabaenaceae-Goniaceae-Volvocaceae (Volvocales, Chlorophyceae). Molecular Phylogenetics and Evolution 130: 1–8.

Nelson DR, Chaiboonchoe A, Fu W, Hazzouri KM, Huang Z, Jaiswal A, Daakour S, Mystikou A, Arnoux M, Sultana M et al. 2019. Potential for heightened sulfur-metabolic capacity in coastal subtropical microalgae. iScience 11: 450–465.

Ness RW, Kraemer SA, Colegrave N, Keightley PD. 2016. Direct estimate of the spontaneous mutation rate uncovers the effects of drift and recombination in the *Chlamydomonas reinhardtii* plastid genome. Mol Biol Evol 33: 800–808.

Ness RW, Morgan AD, Colegrave N, Keightley PD. 2012. Estimate of the spontaneous mutation rate in *Chlamydomonas reinhardtii*. Genetics 192: 1447–1454.

Ness RW, Morgan AD, Vasanthakrishnan RB, Colegrave N, Keightley PD. 2015. Extensive de novo mutation rate variation between individuals and across the genome of *Chlamydomonas reinhardtii*. Genome Res 25: 1739–1749.

Nguyen LT, Schmidt HA, von Haeseler A, Minh BQ. 2015. IQ-TREE: a fast and effective stochastic algorithm for estimating maximum-likelihood phylogenies. Mol Biol Evol 32: 268–274.

Peska V, Garcia S. 2020. Origin, diversity, and evolution of telomere sequences in plants. Front Plant Sci 11: 117.

Plecenikova A, Slaninova M, Riha K. 2014. Characterization of DNA repair deficient strains of *Chlamydomonas reinhardtii* generated by insertional mutagenesis. Plos One 9.

Popescu CE, Borza T, Bielawski JP, Lee RW. 2006. Evolutionary rates and expression level in *Chlamydomonas*. Genetics 172: 1567–1576.

Poulter RTM, Butler MI. 2015. Tyrosine recombinase retrotransposons and transposons. Microbiol Spectr 3: MDNA3-0036-2014.

Prochnik SE, Umen J, Nedelcu AM, Hallmann A, Miller SM, Nishii I, Ferris P, Kuo A, Mitros T, Fritz-Laylin LK et al. 2010. Genomic analysis of organismal complexity in the multicellular green alga *Volvox carteri*. Science 329: 223–226.

Pröschold T, Darienko T, Krienitz L, Coleman AW. 2018. *Chlamydomonas schloesseri* sp nov (Chlamydophyceae, Chlorophyta) revealed by morphology, autolysin cross experiments, and multiple gene analyses. Phytotaxa 362: 21–38.

Pröschold T, Harris EH, Coleman AW. 2005. Portrait of a species: *Chlamydomonas reinhardtii*. Genetics 170: 1601–1610.

Pröschold T, Marin B, Schlösser UG, Melkonian M. 2001. Molecular phylogeny and taxonomic revision of *Chlamydomonas* (Chlorophyta). I. Emendation of *Chlamydomonas* Ehrenberg and *Chloromonas* Gobi, and description of *Oogamochlamys* gen. nov. and *Lobochlamys* gen. nov. Protist 152: 265–300.

Quinlan AR, Hall IM. 2010. BEDTools: a flexible suite of utilities for comparing genomic features. Bioinformatics 26: 841–842.

Raj-Kumar PK, Vallon O, Liang C. 2017. *In silico* analysis of the sequence features responsible for alternatively spliced introns in the model green alga *Chlamydomonas reinhardtii*. Plant Mol Biol 94: 253–265.

Richards S, Liu Y, Bettencourt BR, Hradecky P, Letovsky S, Nielsen R, Thornton K, Hubisz MJ, Chen R, Meisel RP et al. 2005. Comparative genome sequencing of *Drosophila pseudoobscura*: chromosomal, gene, and cis-element evolution. Genome Res 15: 1–18.

Rose AB. 2018. Introns as gene regulators: a brick on the accelerator. Front Genet 9: 672.

Roth MS, Cokus SJ, Gallaher SD, Walter A, Lopez D, Erickson E, Endelman B, Westcott D, Larabell CA, Merchant SS et al. 2017. Chromosome-level genome assembly and transcriptome of the green alga *Chromochloris zofingiensis* illuminates astaxanthin production. Proc Natl Acad Sci U S A 114: E4296–E4305.

Salomé PA, Merchant SS. 2019. A Series of fortunate events: Introducing *Chlamydomonas* as a reference organism. Plant Cell 31: 1682–1707.

Sasso S, Stibor H, Mittag M, Grossman AR. 2018. From molecular manipulation of domesticated *Chlamydomonas reinhardtii* to survival in nature. eLife 7.

Sharp PM, Li WH. 1987. The codon adaptation index - a measure of directional synonymous codon usage bias, and its potential applications. Nucleic Acids Res 15: 1281–1295.

Siepel A, Bejerano G, Pedersen JS, Hinrichs AS, Hou M, Rosenbloom K, Clawson H, Spieth J, Hillier LW, Richards S et al. 2005. Evolutionarily conserved elements in vertebrate, insect, worm, and yeast genomes. Genome Res 15: 1034–1050.

Smit AFA, Hubley R. 2008-2015. RepeatModeler Open-1.0. http://www.repeatmasker.org.

Smit AFA, Hubley R, Green P. 2013-2015. RepeatMasker Open-4.0. http://www.repeatmasker.org.

Smith DR, Lee RW. 2008. Nucleotide diversity in the mitochondrial and nuclear compartments of *Chlamydomonas reinhardtii*: investigating the origins of genome architecture. BMC Evol Biol 8: 156.

Stanke M, Diekhans M, Baertsch R, Haussler D. 2008. Using native and syntenically mapped cDNA alignments to improve *de novo* gene finding. Bioinformatics 24: 637–644.

Stanke M, Schoffmann O, Morgenstern B, Waack S. 2006. Gene prediction in eukaryotes with a generalized hidden Markov model that uses hints from external sources. BMC Bioinformatics 7: 62.

Stark A, Lin MF, Kheradpour P, Pedersen JS, Parts L, Carlson JW, Crosby MA, Rasmussen MD, Roy S, Deoras AN et al. 2007. Discovery of functional elements in 12 *Drosophila* genomes using evolutionary signatures. Nature 450: 219–232.

Stein LD, Bao Z, Blasiar D, Blumenthal T, Brent MR, Chen N, Chinwalla A, Clarke L, Clee C, Coghlan A et al. 2003. The genome sequence of *Caenorhabditis briggsae*: a platform for comparative genomics. PLoS Biol 1: E45.

Stevens L, Felix MA, Beltran T, Braendle C, Caurcel C, Fausett S, Fitch D, Frezal L, Gosse C, Kaur T et al. 2019. Comparative genomics of 10 new *Caenorhabditis* species. Evol Lett 3: 217–236.

Strenkert D, Schmollinger S, Gallaher SD, Salome PA, Purvine SO, Nicora CD, Mettler-Altmann T, Soubeyrand E, Weber APM, Lipton MS et al. 2019. Multiomics resolution of molecular events during a day in the life of *Chlamydomonas*. Proc Natl Acad Sci U S A 116: 2374–2383.

Suh A, Churakov G, Ramakodi MP, Platt RN, 2nd, Jurka J, Kojima KK, Caballero J, Smit AF, Vliet KA, Hoffmann FG et al. 2014. Multiple lineages of ancient CR1 retroposons shaped the early genome evolution of amniotes. Genome Biol Evol 7: 205–217.

Sun Y, Whittle CA, Corcoran P, Johannesson H. 2015. Intron evolution in *Neurospora*: the role of mutational bias and selection. Genome Res 25: 100–110.

Tang H, Bowers JE, Wang X, Ming R, Alam M, Paterson AH. 2008. Synteny and collinearity in plant genomes. Science 320: 486–488.

Valli AA, Santos BA, Hnatova S, Bassett AR, Molnar A, Chung BY, Baulcombe DC. 2016. Most microRNAs in the single-cell alga *Chlamydomonas reinhardtii* are produced by Dicer-like 3-mediated cleavage of introns and untranslated regions of coding RNAs. Genome Res 26: 519–529.

Walker BJ, Abeel T, Shea T, Priest M, Abouelliel A, Sakthikumar S, Cuomo CA, Zeng Q, Wortman J, Young SK et al. 2014. Pilon: an integrated tool for comprehensive microbial variant detection and genome assembly improvement. PLoS One 9: e112963.

Waterhouse RM, Seppey M, Simao FA, Manni M, Ioannidis P, Klioutchnikov G, Kriventseva EV, Zdobnov EM. 2018. BUSCO applications from quality assessments to gene prediction and phylogenomics. Mol Biol Evol 35: 543–548.

Wicker T, Sabot F, Hua-Van A, Bennetzen JL, Capy P, Chalhoub B, Flavell A, Leroy P, Morgante M, Panaud O et al. 2007. A unified classification system for eukaryotic transposable elements. Nat Rev Genet 8: 973–982.

Williamson RJ, Josephs EB, Platts AE, Hazzouri KM, Haudry A, Blanchette M, Wright SI. 2014. Evidence for widespread positive and negative selection in coding and conserved noncoding regions of *Capsella grandiflora*. PLoS Genet 10: e1004622.

Yamamoto K, Kawai-Toyooka H, Hamaji T, Tsuchikane Y, Mori T, Takahashi F, Sekimoto H, Ferris PJ, Nozaki H. 2017. Molecular evolutionary analysis of a gender-limited *MID* ortholog from the homothallic species *Volvox africanus* with male and monoecious spheroids. PLoS One 12: e0180313.

Zhang C, Rabiee M, Sayyari E, Mirarab S. 2018. ASTRAL-III: polynomial time species tree reconstruction from partially resolved gene trees. BMC Bioinformatics 19.

