## Supplementary material for "Comparative genomics of *Chlamydomonas*": All supplementary files: file_S1.pdf

#### Supplementary file 1 for:

### Comparative genomics of *Chlamydomonas*

**Rory J. Craig, Ahmed R. Hasan, Rob W. Ness & Peter D. Keightley**

### High molecular weight DNA extraction protocol

This protocol is a modified version of the *Chlamydomonas* high molecular weight DNA extraction protocol made available by the Joint Genome Institute here: https://www.pacb.com/wp-content/uploads/2015/09/DNA-extraction-chlamy-CTAB-JGI.pdf

The protocol is based on a CTAB and phenol:chloroform extraction, which is performed in two iterations to ensure maximum removal of RNA. The protocol should yield ~10 µg of high molecular weight DNA.

1. Inoculate a 6-well plate with *Chlamydomonas* from a slant culture in 5 ml Bold's basal medium per well and incubate on a plate shaker for 4 days. Up to nine wells can be used per sample (i.e. two plates and 45 ml total), but if higher yield is needed perform multiple extractions per sample.
2. Preheat aliquot of CTAB buffer (see below) to 65 °C.
3. Transfer cultures to a 50 ml Falcon tube, centrifuge to pellet cells and remove media. Resuspend cells in 1 ml of lysis buffer (see below) by gentle pipette mixing and hand mixing in a swirling circular motion.
4. Add 1ml hot CTAB buffer, mix by gentle hand mixing and incubate at 65 °C for 30 minutes.
5. Decant above to 15 ml phase-lock gel tube (e.g. QIAGEN MaXtract) and add 2 ml (1x) room temperature phenol:chloroform:isoamyl (25:24:1). Gently mix by inverting ~40 times per minute for 10 minutes.
6. Centrifuge for 8 minutes at 1,500g (or until phases are clearly separated). Decant aqueous phase to a new 15 ml phase-lock gel tube.
7. Add 5 µL RNAse A. Gently mix and incubate at 37 °C for 30 minutes.
8. Repeat steps 5 & 6 (i.e. phenol:chloroform extraction).
9. Add 1x volume chloroform:isoamyl (49:1) (~2 ml), gently mix by inverting ~40 times per minute for 10 minutes.
10. Centrifuge as in step 6 and decant aqueous phase to a new 15 ml Falcon tube.
11. Add 2x volume ice-cold 100% ethanol, slowly mix by inverting 10 times end over end and incubate on ice for at least an hour.
12. Carefully decant above to four 1.5 ml DNA lo-bind tubes and centrifuge at 13,000g for 15 minutes at 4°C. The DNA should be visible by eye, and in most cases the majority of the DNA will be in a single 1.5 ml tube as it would have formed a large clump in the previous 15 ml tube.
13. Discard supernatant in each tube and wash with 1 ml freshly prepared 70% ethanol (using DNAse-free water), invert gently and centrifuge at 13,000g for one minute. Discard supernatant and repeat.

14. After removing majority of supernatant, briefly spin down washed pellets and remove any remaining ethanol by pipette. Dry in flow hood for ~5 minutes (ensure all ethanol has evaporated but be careful not to over-dry pellet).
15. Resuspend pellets overnight in 45 µl TE buffer.
16. After checking that the DNA has completely eluted, combine four tubes by gently pipetting 45 µl from three of the tubes to one tube (ideally pipette into the tube containing the majority of the DNA to minimise shearing). Add 20 µl NEB3 buffer 10x.
17. Add 8 µl of RNAse $I_f$, mix by gentle flicking and inversion, spin-down, and incubate for 30 minutes at 37 °C.
18. Add 2 µl of RiboShredder, incubate for a further 30 minutes at 37 °C.
19. Add 340 µl of DNAse-free water to bring total volume to 550 µl.
20. Add 550 µl phenol:chloroform:isoamyl (25:24:1), mix briefly and decant to a 2 ml phase-lock gel tube. As before, mix gently for 10 minutes and spin-down at room temperature at 14,000g.
21. Decant to second 2 ml phase-lock gel tube, add 1x chloroform:isoamyl (49:1), mix and spin-down as before.
22. Using a P1000, pipette 450 µl of the aqueous phase to a new 1.5 ml DNA lo-bind tube. Add 50 µl 3 M sodium acetate (pH 5.2) and mix gently by inversion.
23. Add 2x ice-cold ethanol (1 ml), precipitate and wash as in steps 12-14. Elute DNA pellet in 183 µl of TE buffer overnight.
24. QC sample – Qubit 2 µl, Nanodrop 1µl. Pulse-field gel electrophoresis can be performed to check DNA fragment length distribution.

Lysis Buffer (10ml)

50 mM Tris-HCl (pH 8.0) – 0.5 ml 1 M stock

200 mM NaCl – 0.4 ml 5 M stock

20 mM EDTA – 0.4 ml 0.5 M stock

Nuclease-free water – 6.2 ml

2% SDS – 2 ml

Proteinase K (20 mg/ml) – 0.5 ml

CTAB buffer (10ml)

50 mM Tris-HCl (pH 8.0) – 1 ml 1 M stock

1.4 M NaCl – 2.8 ml 5 M stock

20 mM EDTA – 0.4 ml 0.5 M stock

2% CTAB – 0.2 g

1% PVP 40,000 – 0.1 g

Nuclease-free water to 10 ml
