## Supplementary material for "Comparative genomics of *Chlamydomonas*": All supplementary files: file_S2.pdf

### Supplementary file 2 for:

### Comparative genomics of *Chlamydomonas*

**Rory J. Craig, Ahmed R. Hasan, Rob W. Ness & Peter D. Keightley**

### Detailed genome assembly methods

#### *Chlamydomonas incerta* SAG 7.73

The *C. incerta* genome was assembled from 6.31 Gb of PacBio long-read data, with a mean read length of 7.69 kb and an N50 read length of 13.71 kb. In order to estimate genome size and assess the library for possible contaminants, a preliminary genome assembly was produced using miniasm. The taxonomic origin of the resulting contigs was determined by comparison to the NCBI nucleotide collection database (nt) by BLAST+ megablast, and a taxon-annotated GC-coverage plot was produced using Blobtools. This identified a single low-coverage Proteobacteria contaminant, and all reads mapping to contigs identified as bacterial were filtered. Canu was run using the genome size of the miniasm assembly (130.8 Mb), and as a precaution the Blobtools pipeline was re-run on the resulting assembly, and any further reads identified as bacterial were filtered. In total <1% of reads were identified as bacterial over both filtering steps. Canu was then re-run using the final contaminant-filtered dataset. The resulting assembly underwent three rounds of iterative polishing by mapping the PacBio reads using pbalign and performing error correction using the Arrow module of the GenomicConsensus tool.

Further error-correction was performed using ~86x coverage of genomic short-read data and 8.20 Gb of RNA-seq data. The genomic short-read data consisted of two libraries of 100 bp paired-end reads with an insert size of ~180 bp (i.e. overlapping read pairs), and two mate-pair libraries of 100 bp reads with an insert size of ~5000 bp. The short insert size libraries were pre-processed by trimming low-quality bases and adapter sequence using the BBtools program bbduk.sh, mapping the trimmed reads to the Arrow-polished assembly using bwa mem, and filtering putative PCR duplicates using Picard MarkDuplicates. For each of the two libraries, the resulting read pairs were then merged using bbmerge-auto.sh to create single reads from the overlapping read pairs where possible. For each library, this resulted in one dataset of unpaired reads (i.e. merged read pairs) and one dataset of paired reads (read pairs that could not be merged). The mate-pair libraries were trimmed of junction adapters and classified as genuine mate-pairs, paired-end, or unknowns using NxTrim. For each library and each classification, pre-processing was performed as described above. Final classification as genuine mate-pairs (i.e. reads pairs with outward facing orientations) or normal paired-end (reads with inward facing orientations) was achieved by assessing the read orientation in the appropriate BAM file. This resulted in one dataset of long-insert mate pairs and one dataset of short-insert paired-end reads per library. All genomic short-read datasets were mapped to the Arrow-polished assembly using bwa mem prior to polishing. The RNA-seq dataset consisted of a single library of stranded 150 bp paired-end reads. Quality and adapter trimming were performed with Trimmomatic, and reads were mapped to the Arrow-polished assembly using STAR in 2-pass mode. To perform error correction, Pilon was then run by providing all BAM files of aligned short-reads (2x merged single reads and 2x unmerged paired-end reads from the short insert libraries, 2x mate-pair reads and 2x paired-end reads from the long insert libraries, and 1x RNA-seq paired-end reads). The option "--fix bases" was used to avoid genuine introns being removed due to the spliced mapping of the RNA-seq data. Even with this option we noticed that Pilon corrected a number of large indels, and upon manually checking a sub-sample of such cases using IGV we found that the majority of indels were not supported by the PacBio reads and appeared to be caused by multiply mapping Illumina reads (i.e. the indels appeared to be heterozygous in the Illumina data). We therefore used a custom Perl script to restore all indels >5 bp to their unpolished form after running Pilon. Pilon was run iteratively three times, with all short-read data re-mapped using bwa mem/STAR between each iteration.

The optimal number of polishing iterations for Arrow and Pilon were determined using two metrics: the number of complete single-copy orthologs identified by BUSCO (genome mode, Eukaryota odb9 dataset), and the sequence similarity between an existing *C. incerta* expressed sequence tag library (Popescu et al. 2006) and the contigs as assessed by megablast. Polishing was deemed complete when there was no increase in both metrics between iterations.

Final processing was performed by removing any contigs supported by only a single PacBio read (i.e. "reads=1" in the Canu fasta definition line) and by filtering the two mitochondrion and plastid contigs (which will be described elsewhere). Contigs were ordered by size and given unique IDs with the format CXXXX, where XXXX represent ordered numbers (i.e. the largest contig was named C0001). Potential misassemblies were identified via synteny analysis to *C. reinhardtii* (see main text). All breakpoints between synteny blocks on a given contig that resulted in a transition between *C. reinhardtii* chromosomes were checked manually using IGV and alignments of the PacBio reads. This resulted in four contigs being split due to likely misassemblies, with split contigs having either "a" or "b" appended to their contig ID.

| Software/program | Version | Command line options | Reference(s) |
|---|---|---|---|
| miniasm | 0.3-r179 | | Li (2016) |
| Blobtools | v1.0 | | Laetsch and Blaxter (2017) |
| Canu | 1.7.1 | genomeSize=130.8m correctedErrorRate=0.065 corMhapSensitivity=normal | Koren et al. (2017) |
| pbalign | 0.3.1 | | https://github.com/PacificBiosciences/pbalign |
| Arrow | 2.3.2 | | https://github.com/PacificBiosciences/GenomicConsensus |
| bbduk.sh | 38.16 | ktrim=r k=23 mink=11 hdist=1 tbo | https://jgi.doe.gov/data-and-tools/bbtools/ |
| bbmerge-auto.sh | 38.16 | | https://jgi.doe.gov/data-and-tools/bbtools/ |
| bwa mem | 0.7.17-r1188 | | Li and Durbin (2009) |
| samtools | 1.9 | | Li et al. (2009) |
| picard MarkDuplicates | 2.18.11-SNAPSHOT | REMOVE_DUPLICATES=true | http://broadinstitute.github.io/picard/ |
| Nxtrim | v0.4.3-6eb8d5e | --rf | O'Connell et al. (2015) |
| trimmomatic | 0.38 | PE LEADING:3 TRAILING:3 SLIDINGWINDOW:4:3 MINLEN:25 | Bolger et al. (2014) |
| STAR | STAR_2.6.1a | --twopassMode Basic | Dobin et al. (2013) |
| Pilon | 1.22 | --fix bases | Walker et al. (2014) |
| IGV | 2.7.2 | | Robinson et al. (2011) |

Software and programs used for the assembly of *C. incerta*. Full pipeline is available at: https://github.com/rorycraig337/Chlamydomonas_comparative_genomics/tree/master/genome_assemblies/c_incerta/

#### *Chlamydomonas schloesseri* CCAP 11/173

The *C. schloesseri* genome was assembled from 6.18 Gb of PacBio long-read data, with a mean read length of 7.17 kb and an N50 read length of 12.52 kb. Preliminary assembly and contaminant assessment were performed as per *C. incerta*. No putative contaminant contigs were found for the miniasm assembly, although two were identified for the initial Canu assembly. Reads mapping to these contigs were filtered and Canu was subsequently re-run. Three iterative rounds of polishing were performed with Arrow.

Error correction with Pilon was performed using ~71x coverage of genomic Illumina data (four libraries, 125 bp paired-end) and 7.42 Gb of RNA-seq (one library, stranded 100 bp paired-end). Low quality bases and adapter sequences were trimmed using bbduk.sh (genomic data) or Trimmomatic (RNAseq), and putative PCR duplicates were removed from the genomic data using picard MarkDuplicates. The resulting datasets were mapped to the Arrow-polished assembly using bwa mem (genomic data) or STAR (RNAseq), and the resulting BAM files were passed to Pilon. Illumina based polishing was iterated three times.

Final processing was performed by removing any contigs supported by only a single PacBio read (i.e. "reads=1" in the Canu fasta definition line) and by filtering the mitochondrion and plastid contigs. Potential misassemblies were identified via synteny analysis to *C. reinhardtii* (see main text). All breakpoints between synteny blocks on a given contig that resulted in a transition between *C. reinhardtii* chromosomes were checked manually using IGV and alignments of the PacBio reads. This resulted in two contigs being split due to likely misassemblies. Contigs were named as per *C. incerta*.

| **Software/program** | **Version** | **Command line options** | **Reference(s)** |
|---|---|---|---|
| miniasm | 0.3-r179 | | Li (2016) |
| Blobtools | v1.0 | | Laetsch and Blaxter (2017) |
| Canu | 1.7.1 | genomeSize=130.5m correctedErrorRate=0.065 corMhapSensitivity=normal | Koren et al. (2017) |
| pbalign | 0.3.1 | | https://github.com/PacificBiosciences/pbalign |
| Arrow | 2.3.2 | | https://github.com/PacificBiosciences/GenomicConsensus |
| bbduk.sh | 38.16 | ktrim=r k=23 mink=11 hdist=1 tbo | https://jgi.doe.gov/data-and-tools/bbtools/ |
| bwa mem | 0.7.17-r1188 | | Li and Durbin (2009) |
| samtools | 1.9 | | Li et al. (2009) |
| picard MarkDuplicates | 2.18.11-SNAPSHOT | REMOVE_DUPLICATES=true | http://broadinstitute.github.io/picard/ |
| trimmomatic | 0.38 | PE LEADING:3 TRAILING:3 SLIDINGWINDOW:4:3 MINLEN:25 | Bolger et al. (2014) |
| STAR | STAR_2.6.1a | --twopassMode Basic | Dobin et al. (2013) |
| Pilon | 1.22 | --fix bases | Walker et al. (2014) |
| IGV | 2.7.2 | | Robinson et al. (2011) |

Software and programs used for the assembly of *C. schloesseri*. Full pipeline is available at: https://github.com/rorycraig337/Chlamydomonas_comparative_genomics/tree/master/genome_assemblies/c_schloesseri/

***Edaphochlamys debaryana* CCAP 11/70**

The *E. debaryana* genome was assembled from 5.70 Gb of PacBio long-read data, with a mean read length of 7.82 kb and an N50 read length of 13.46 kb. Using Blobtools as described above, no contaminant contigs were detected in either the preliminary miniasm assembly or the subsequent Canu assembly. We therefore proceeded with the initial Canu assembly, which was polished with Arrow (three iterations) using all available PacBio reads.

Error correction with Pilon was performed using ~43x coverage of genomic Illumina data (one library, 150 bp paired-end) and 7.48 Gb of RNA-seq (one library, stranded 100 bp paired-end). Pre-processing was performed as described for *C. incerta* and *C. schloesseri*, and Pilon was run iteratively three times.

Final processing was performed by removing any contigs supported by only a single PacBio read (i.e. “reads=1” in the Canu fasta definition line) and by filtering the plastid contig (no mitochondrial contigs were detected). Contigs were named as per *C. incerta*.

| Software/program | Version | Command line options | Reference(s) |
|---|---|---|---|
| miniasm | 0.3-r179 | | Li (2016) |
| Blobtools | v1.0 | | Laetsch and Blaxter (2017) |
| Canu | 1.7.1 | genomeSize=148.9m correctedErrorRate=0.065 corMhapSensitivity=normal | Koren et al. (2017) |
| pbalign | 0.3.1 | | https://github.com/PacificBiosciences/pbalign |
| Arrow | 2.3.2 | | https://github.com/PacificBiosciences/GenomicConsensus |
| bbduk.sh | 38.16 | ktrim=r k=23 mink=11 hdist=1 tbo | https://jgi.doe.gov/data-and-tools/bbtools/ |
| bwa mem | 0.7.17-r1188 | | Li and Durbin (2009) |
| samtools | 1.9 | | Li et al. (2009) |
| picard MarkDuplicates | 2.18.11-SNAPSHOT | REMOVE_DUPLICATES=true | http://broadinstitute.github.io/picard/ |
| trimmomatic | 0.38 | PE LEADING:3 TRAILING:3 SLIDINGWINDOW:4:3 MINLEN:25 | Bolger et al. (2014) |
| STAR | STAR_2.6.1a | --twopassMode Basic | Dobin et al. (2013) |
| Pilon | 1.22 | --fix bases | Walker et al. (2014) |

Software and programs used for the assembly of *E. debaryana*. Full pipeline is available at: https://github.com/rorycraig337/Chlamydomonas_comparative_genomics/tree/master/genome_assemblies/e_debaryana/

#### *Polytomella parva* SAG 63-3 (assembly & annotation)

In order to include a more appropriate outgroup to the core-*Reinhardtinia* clade for phylogenomic analyses, we assembled and annotated a draft genome for *P. parva* using publicly available data. *Polytomella* are a genus of non-photosynthetic algae within the *Reinhardtinia* clade, and thus *P. parva* is more closely related to the core-*Reinhartinina* than other available annotated genomes (e.g. *Dunaliella salina* and *Chlamydomonas eustigma*).

The draft assembly was produced using 2.64 Gb of genomic Illumina data (50 bp paired-end reads) sequenced by Smith and Lee (2014). Low-quality and adapter sequences were trimmed using bbduk.sh, and assembled using SPAdes. The assembly contained no contaminant contigs as assessed by Blobtools. Scaffolds <500 bp were filtered. The transcriptome assembly produced by Johnson et al. (2019), using data sequenced by Keeling et al. (2014), was aligned to the SPAdes assembly using BLAT, and the resulting alignments were used to perform additional scaffolding with SCUBAT2.

A *de novo* repeat library was produced using RepeatModeler, which was combined with all manually annotated repeat models for *Chlamydomonas reinhardtii*, *C. incerta*, *C. schloesseri*, and *E. debaryana*, and Repbase sequences for *Volvox carteri*. Interspersed repeats were then softmasked using RepeatMasker and the custom repeat library. The raw RNAseq data (Keeling et al. 2014) was trimmed using Trimmomatic, and aligned to the repeat-masked genome using STAR. The resulting alignments were used to perform gene annotation using BRAKER. Gene models containing internal stop codons, protein sequences <30 amino acids, or coding sequence overlap >=30% with interspersed repeats or >=70% with low-complexity/simple repeats were filtered.

| Software/program | Version | Command line options | Reference(s) |
|---|---|---|---|
| bbduk.sh | 38.16 | ktrim=r k=23 mink=11 hdist=1 tbo | https://jgi.doe.gov/data-and-tools/bbtools/ |
| SPAdes | v3.13.0 | --careful | Bankevich et al. (2012) |
| Blobtools | v1.0 | | Laetsch and Blaxter (2017) |
| BLAT | v. 36 | -t=dna -q=dna | Kent (2002) |
| SCUBAT2 | v2 | | https://github.com/GDKO/SCUBAT2 |
| RepeatModeler | 1.0.11 | | Smit and Hubley (2008-2015) |
| RepeatMasker | 4.0.9 | -nolow -a -xsmall -gccalc | Smit et al. (2013-2015) |
| trimmomatic | 0.38 | PE LEADING:3 TRAILING:3 SLIDINGWINDOW:4:3 MINLEN:25 | Bolger et al. (2014) |
| STAR | STAR_2.6.1a | --twopassMode Basic | Dobin et al. (2013) |
| BRAKER | 2.1.2 | --softmasking | Hoff et al. (2016); Hoff et al. (2019) |

Software and programs used for the assembly of *P. parva*.

#### References


Bankevich A, Nurk S, Antipov D, Gurevich AA, Dvorkin M, Kulikov AS, Lesin VM, Nikolenko SI, Pham S, Prjibelski AD et al. 2012. SPAdes: a new genome assembly algorithm and its applications to single-cell sequencing. *Journal of Computational Biology* **19**: 455-477.

Bolger AM, Lohse M, Usadel B. 2014. Trimmomatic: a flexible trimmer for Illumina sequence data. *Bioinformatics* **30**: 2114-2120.

Dobin A, Davis CA, Schlesinger F, Drenkow J, Zaleski C, Jha S, Batut P, Chaisson M, Gingeras TR. 2013. STAR: ultrafast universal RNA-seq aligner. *Bioinformatics* **29**: 15-21.

Hoff KJ, Lange S, Lomsadze A, Borodovsky M, Stanke M. 2016. BRAKER1: unsupervised RNA-Seq-based genome annotation with GeneMark-ET and AUGUSTUS. *Bioinformatics* **32**: 767-769.

Hoff KJ, Lomsadze A, Borodovsky M, Stanke M. 2019. Whole-genome annotation with BRAKER. *Methods in Molecular Biology* **1962**: 65-95.

Johnson LK, Alexander H, Brown CT. 2019. Re-assembly, quality evaluation, and annotation of 678 microbial eukaryotic reference transcriptomes. *Gigascience* **8**.

Keeling PJ, Burki F, Wilcox HM, Allam B, Allen EE, Amaral-Zettler LA, Armbrust EV, Archibald JM, Bharti AK, Bell CJ et al. 2014. The Marine Microbial Eukaryote Transcriptome Sequencing Project (MMETSP): illuminating the functional diversity of eukaryotic life in the oceans through transcriptome sequencing. *PLoS Biol* **12**.

Kent WJ. 2002. BLAT - The BLAST-like alignment tool. *Genome Res* **12**: 656-664.

Koren S, Walenz BP, Berlin K, Miller JR, Bergman NH, Phillippy AM. 2017. Canu: scalable and accurate long-read assembly via adaptive k-mer weighting and repeat separation. *Genome Res* **27**: 722-736.

Laetsch DR, Blaxter M. 2017. BlobTools: Interrogation of genome assemblies. *F1000Research* **6**.

Li H. 2016. Minimap and miniasm: fast mapping and de novo assembly for noisy long sequences. *Bioinformatics* **32**: 2103-2110.

Li H, Durbin R. 2009. Fast and accurate short read alignment with Burrows-Wheeler transform. *Bioinformatics* **25**: 1754-1760.

Li H, Handsaker B, Wysoker A, Fennell T, Ruan J, Homer N, Marth G, Abecasis G, Durbin R, Genome Project Data Processing S. 2009. The Sequence Alignment/Map format and SAMtools. *Bioinformatics* **25**: 2078-2079.

O'Connell J, Schulz-Trieglaff O, Carlson E, Hims MM, Gormley NA, Cox AJ. 2015. NxTrim: optimized trimming of Illumina mate pair reads. *Bioinformatics* **31**: 2035-2037.

Popescu CE, Borza T, Bielawski JP, Lee RW. 2006. Evolutionary rates and expression level in *Chlamydomonas*. *Genetics* **172**: 1567-1576.

Robinson JT, Thorvaldsdóttir H, Winckler W, Guttman M, Lander ES, Getz G, Mesirov JP. 2011. Integrative Genomics Viewer. *Nat Biotechnol* **29**: 24-26.

Smit AFA, Hubley R. 2008-2015. RepeatModeler Open-1.0. *http://wwwrepeatmasker.org*.

Smit AFA, Hubley R, Green P. 2013-2015. RepeatMasker Open-4.0. *http://wwwrepeatmasker.org*.

Smith DR, Lee RW. 2014. A plastid without a genome: evidence from the nonphotosynthetic green algal genus *Polytomella*. *Plant Physiology* **164**: 1812-1819.

Walker BJ, Abeel T, Shea T, Priest M, Abouelliel A, Sakthikumar S, Cuomo CA, Zeng Q, Wortman J, Young SK et al. 2014. Pilon: an integrated tool for comprehensive microbial variant detection and genome assembly improvement. *PLoS One* **9**: e112963.
